# Vipsania: Unsupervised Deep Gene Finding

**DOI:** 10.64898/2026.08.26.747235

**Authors:** Richard Krieg, Felix Becker, Stepan Saenko, Joscha Diehl, Mario Stanke

**Affiliations:** Universität Greifswald, Institut für Mathematik und Informatik

## Abstract

Scaling the structural annotation of protein-coding genes to all eukaryotic genomes remains a major challenge. While recent deep learning methods rival evidence-based pipelines without requiring RNA-seq or alignments, they are entirely supervised. They depend on large, high-quality training sets from diverse genomes, leaving many basal eukaryotic clades without an accurate *ab initio* gene finder. We present Vipsania, the first unsupervised deep gene finder. A differentiable hidden Markov layer inside a deep sequence model learns to predict gene structures from unannotated genomes alone. Vipsania is pretrained for virtually all eukaryotes and finetunes without supervision on the target genome. It is, on average, more accurate than supervised methods across most clades and avoids the accuracy drop that supervised models suffer on distant target genomes. Vipsania adapts to non-standard genetic codes and provides a fast and highly versatile tool for unbiased, pan-eukaryotic genome annotation. The source code is available at https://github.com/gaius-augustus/vipsania.

---

Determining where the protein-coding genes of a newly sequenced eukaryotic genome lie, and what their exon–intron structures are, is the foundation for nearly every downstream analysis, but still not solved [JPS26]. The most accurate approach today is to collect extrinsic evidence for each new genome: evidence-based pipelines such as **Braker3** [Gab+24a] run hidden Markov models (HMMs) directly on the DNA, with parameters trained from short-read RNA-seq and the annotated proteomes of related species. The recent **EviAnn** [Zim+26] focuses on evidence-rich genomes and builds gene structures directly from transcript and protein alignments, training Markov chains only to screen the splice sites of these candidates rather than to predict genes. Such pipelines scale poorly across all eukaryotes: RNA-seq, ideally from diverse tissues and conditions, must be produced for each species and homologous proteins collected, and the spliced alignment of both is computationally expensive. Their accuracy is also bounded by the evidence at hand, which is thinnest for genomes with no well-annotated relative.

The Earth BioGenome Project [Lew+18] aims to generate a reference genome for each of the about 1.67 million named eukaryotic species, with an intermediate goal of representative genomes for at least half of all accepted genera [Bla+25]. In August 2026, there were about 24 thousand eukaryotic organisms (1.4% of eukaryotes) with at least one scaffold-level assembly in the NCBI database. Annotation is already the limiting step rather than sequencing: fewer than 20% of the genomes available at NCBI Datasets carry a gene annotation [Gab+26]. The shortfall is not spread evenly across the tree of life; it is most severe exactly where the least prior knowledge exists, leaving entire branches such as the eukaryotic algae, where most assemblies still carry no structural annotation at all [KHH23]. Absolute accuracy leaves room as well: on the human genome the best *ab initio* predictor reaches a gene-level F1 score of 62% [Gab+24b].

Deep learning has produced two approaches, so far developed separately. The first is a generation of deep *ab initio* gene finders. **Helixer** [Hol+25] passes convolutional and recurrent representations to an HMM post-processor, **ANNEVO** [Zha+26] does the same from attention and mixture-of-experts layers, and **Tiberius** [Gab+24b] makes the HMM at the last layer *differentiable*, incorporating it into the computational graph during training rather than treating it as a decoding step afterwards. An HMM is the one component they all use, and what makes them gene finders rather than classifiers: it carries the grammar of a gene — reading frames maintained across introns, no in-frame stop codons — and so turns per-position scores into translatable structures. Needing no extrinsic evidence, these deep gene finders are fast: **Tiberius** runs on a GPU on average 80 times faster than **Braker3** while approaching its accuracy in several clades [Gab+26]. All of these deep gene finders, however, depend on supervision: they are trained on reference gene sets assumed to be accurate, so the ceiling of the approach is set by how many high-quality reference genes exist [JPS26]. The released clade models of **Helixer** and **ANNEVO** each cover, according to their designation, 96.1% of the eukaryotic species in the NCBI Taxonomy, those of **Tiberius** 79.6%; 50,218 species (2.9%) — among them all of Rhodophyta, Alveolata, Discoba and Amoebozoa — are covered by none of the three. We here benchmark that **Helixer** and **ANNEVO** have average gene-level F1 scores of only 15.1% and 20.7% on invertebrates that are not arthropods, which make up 135,036 species according to the NCBI Taxonomy, reducing their effective applicability to only about 88% of eukaryotic species. The unsupervised self-training of **GeneMark** [Lom+05] is the only broadly applicable alternative. Supervision also makes accuracy hard to read: each method has its own training set, and a reported score depends via phylogenetic leakage on how close the test species is to those particular genomes — a distance that differs between tools and is not controlled for. Systematic analyses of phylogenetic leakage, and of how accuracy decays with the distance between target and training species, are missing.

The second deep-learning approach uses a genome foundation model (GFM), trained self-supervised on raw DNA [Zho+23]. Transformers remain widely used, but attention scales quadratically [Vas+17] while eukaryotic genes can span hundreds of kilobases, and the models that reach the longest contexts at single-nucleotide resolution instead use subquadratic backbones: **Caduceus** [Sch+24] builds on bidirectional state space models, and **Evo 2** [Bri+26] reaches a unidirectional context of one million tokens and develops internal features for exon–intron boundaries without ever being shown an annotation. Yet the annotation ability of GFMs is almost always reported per base, as splice-site or coding/noncoding classification [Mar+24; Fen+25]. However, the first deliverable of a genome annotation method that scales across all eukaryotic genomes is a set of protein sequences and it should be accurate at interval-level metrics such as transcript sensitivity [PP20] as per-base accuracy does not imply them. Benchmarking the representations directly confirms this: pretrained DNA language model embeddings “do not capture the features necessary for precise gene segmentation, and […] task-specific fine-tuning remains essential” [Shm+26]. Where gene grammar is added, it is bolted on at the decoding stage: **GeneCAD** [Liu+25] decodes with a conditional random field while the foundation model is frozen. Further, it is restricted to flowering plants.

We present **Vipsania**, which combines the gene-structure HMM, as used by deep gene finders, with the unsupervised objective of foundation models, in a single, fully differentiable deep learning model. **Vipsania** is trained like GFMs to predict masked nucleotides [Dev+19], and its HMM layer [BS22; BS24; Gab+24b] is placed in the middle of the layer stack. Its structural constraint shapes the learned representations instead of being imposed afterwards, supplying by construction and during training what supervised finetuning or postprocessing otherwise must provide. It uses linear recurrent units (LRUs) [Orv+23], which cover both long-range and short-range dependencies. Convolutional layers serving as input encodings, as employed by other deep gene finders, are not used. **Vipsania** uses single nucleotide resolution throughout and its HMM tracks the reading frame explicitly. A novel loss is used, which prioritizes nucleotide prediction accuracy on *predicted* gene borders based on HMM posterior state probabilities. No reference annotation is used at any point. We release models pretrained on 17 clades that together account for 99.4% of eukaryotic taxa in the NCBI Taxonomy; the remaining genomes can be annotated after unsupervised finetuning of a general model for the remaining 0.6% of eukaryotic species; and non-standard genetic codes are accommodated by minimal specifications and single-species training. **Vipsania** achieves a higher mean locus-level F1 score than supervised deep gene finders in every clade except Vertebrata. We further quantify the distance between each target genome and each method’s training genomes, and show that supervised gene finders lose accuracy as that distance grows whereas **Vipsania** does not, and that **Vipsania** is immune to overfitting to the training set.

## Results

The unsupervised training objective of **Vipsania** enables the model to produce a meaningful genome annotation for nearly any eukaryotic species. We release pretrained models for 17 major taxonomic clades that together cover more than 99.4% of all eukaryotes represented in the NCBI taxonomy database. Figure 1 represents the coverage of **Vipsania** across the taxonomic eukaryotic tree of life. There are only 3 clades with at least 8 genomes of assembly level “scaffold” or higher that are not covered by the released checkpoints. We release an additional **Vipsania** model for this complement of all already covered eukaryotes as well.

**Figure 1.**
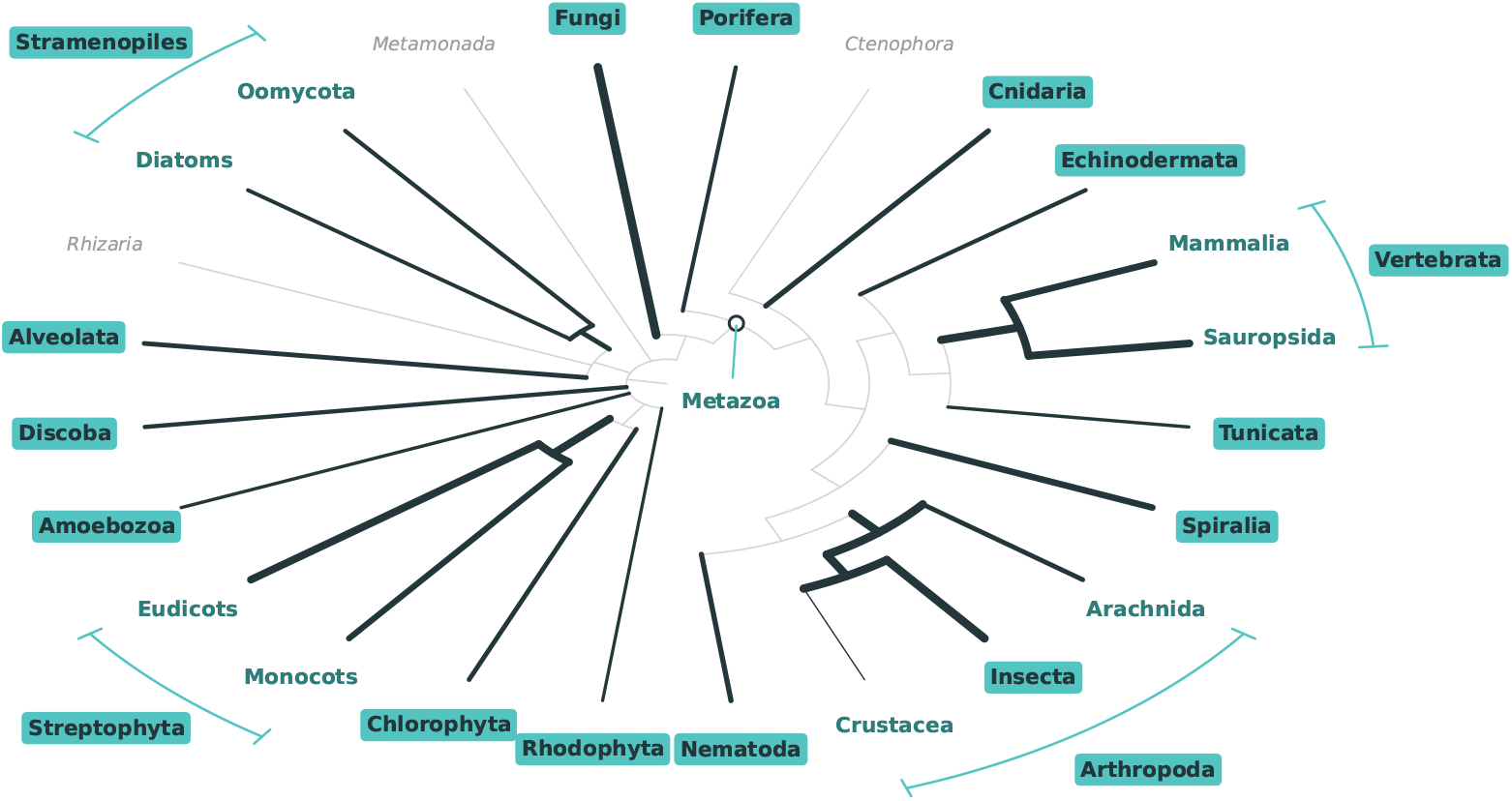
A simplified version of the taxonomic eukaryotic tree of life. We release **Vipsania** checkpoints for the highlighted clades. The branch widths are proportional to the logarithm of the number of available genomes in the clade with an assembly level of at least “scaffold”, inferred from the NCBI database. Clades that are not covered by the highlighted ones are shown if there are at least 8 genomes of the same quality available.

All **Vipsania** models share the same general architecture, see Figure 2. A residual stream connects LRU [Orv+23] and GLU [Sha20] layers, as well as a differentiable HMM layer. This layer yields state probabilities during training, which are funneled back into the residual stream, and discrete Viterbi paths during inference, which are formed into a genome annotation. By predicting masked nucleotides given a small surrounding genome context, **Vipsania** learns to adapt the HMM parameters to model gene structures. A special loss guides the model towards more accurate nucleotide predictions at gene borders, without using external gene structure information.

**Figure 2.**
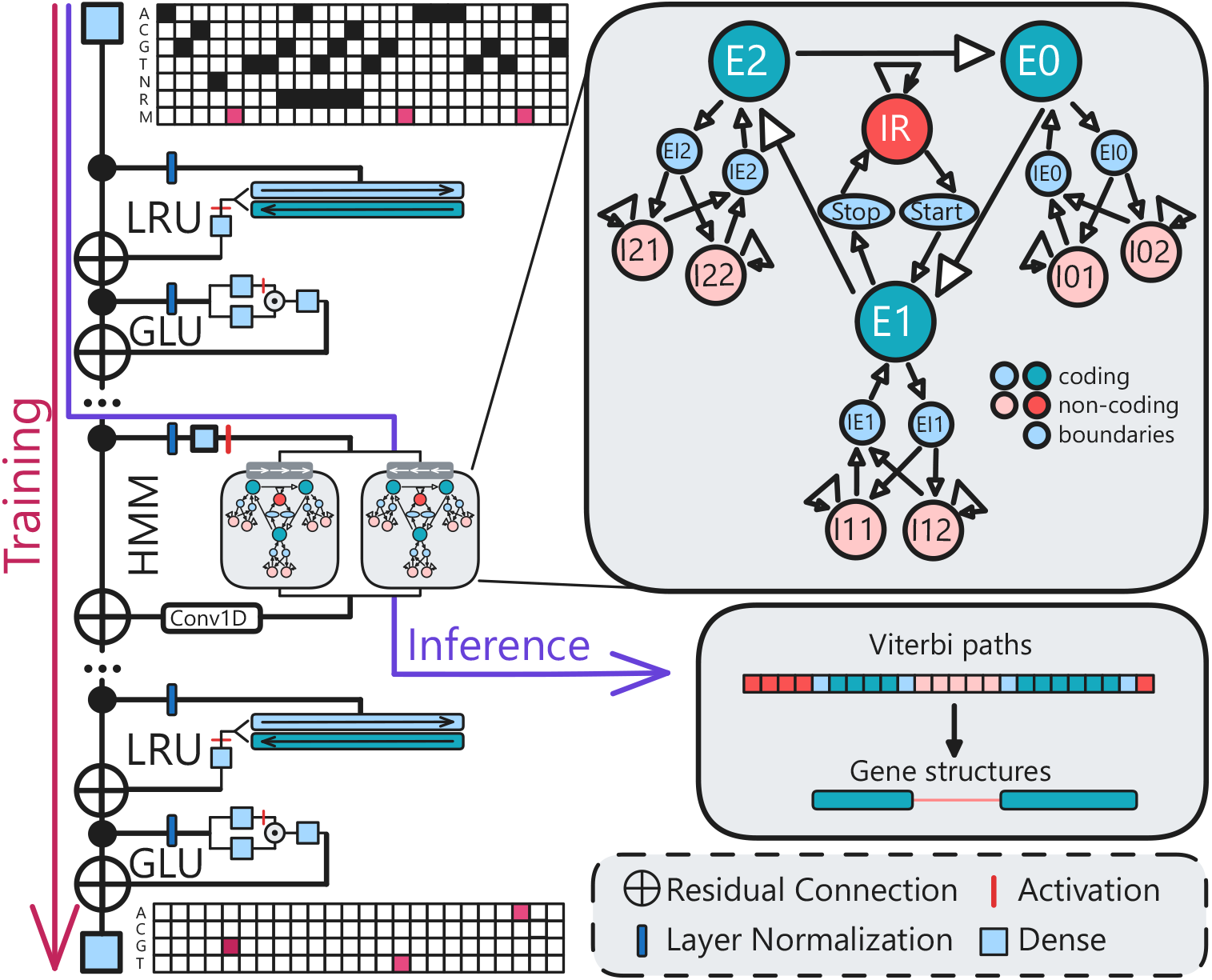
The model architecture of **Vipsania.** During training, masked nucleotides are predicted based on sequence context. For inference, Viterbi paths of the HMM are translated into gene structures. The HMM layer processes forward and backward strand simultaneously during training and inference.

For each clade, we finetune and test the corresponding **Vipsania** model on up to 13 test species. As no genome annotation data was used during training, each model can be evaluated without the risk of biased predictions or overfitting. We compare our results to **Helixer** [Hol+25], **Tiberius** [Gab+26], **ANNEVO** [Zha+26] and **GeneMark** [Lom+05] on clades for which they were trained (Figure 3). For each clade, we compute the mean of F1 scores for all test species for exon and locus level for the coding regions of protein-coding genes. A bar in each cell visualizes the accuracy gains of **Vipsania** over each specific tool.

**Figure 3.**
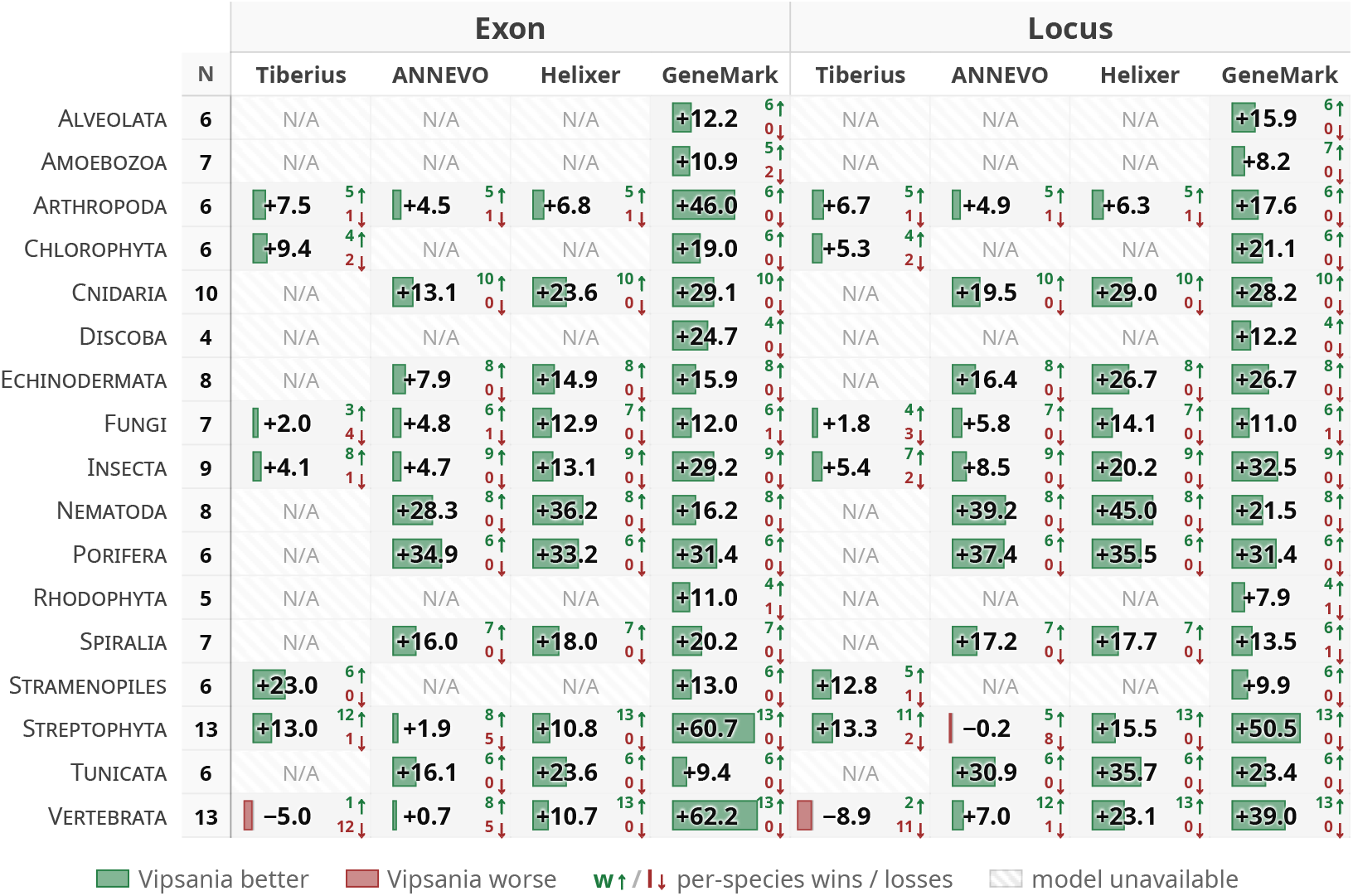
Mean of F1 score differences (**Vipsania** minus other program) across all clades. Reported are accuracy gains of finetuned **Vipsania** for exon and locus level. The green and red numbers show the number of the *N* test species where **Vipsania** had a higher or lower F1 score, respectively.

Across all clades, **Vipsania** shows a better average locus F1 score, with the exception of Vertebrata. **Vipsania** also wins the comparison for most individual species (numbers next to arrows in Figure 3); and the better average performance is not due to outlier poor performances of the other tools on particular test species. In Figure 4 absolute accuracies are compared for the three important clades of insects, fungi and land plants. Here, on 23 out of the 29 test species, **Vipsania** achieves a locus F1 score of 50% or higher. In 4 out of 17 clades, no appropriate checkpoint exists for any other deep learning gene finder. On the invertebrate clades Cnidaria, Echinodermata, Nematoda, Porifera, Spiralia and Tunicata, **Vipsania** exceeds **ANNEVO** annotations on average by 26 percentage points locus F1, and **Helixer** on average by 31 percentage points. The only global alternative that provides annotations across all tested clades is **GeneMark. Vipsania** is more accurate than **GeneMark** in all 17 clades, on average by 22 percentage points locus F1 (Figure 3). Detailed results can be found in Supplementary Tables 5–21.

**Figure 4.**
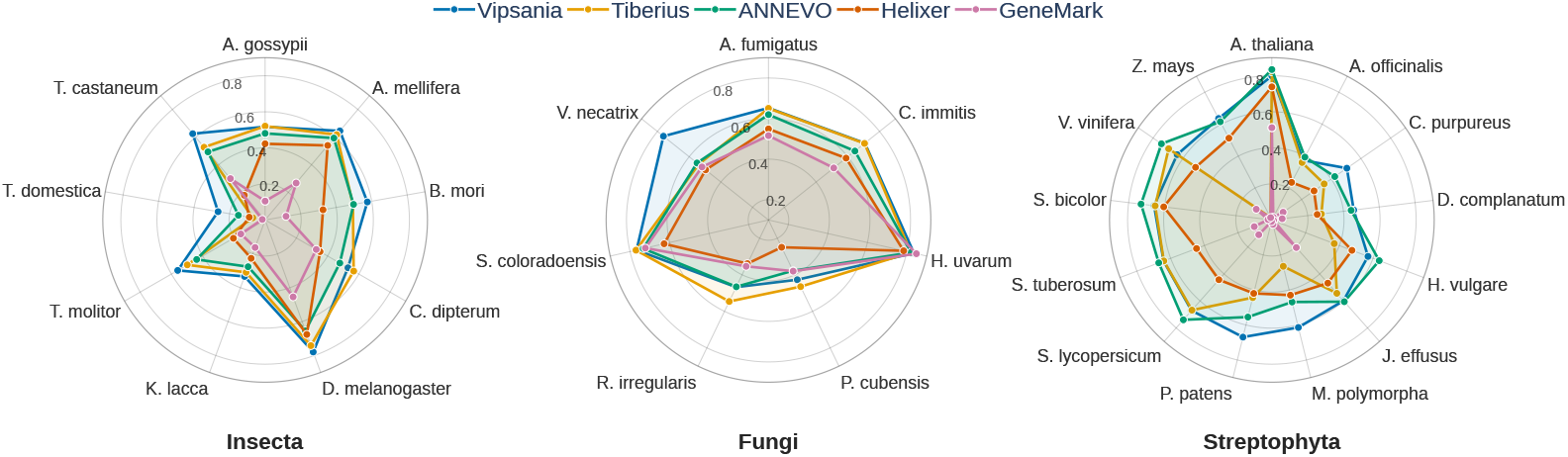
Locus F1 metrics of test species across the clades Insecta, Fungi and Streptophyta.

### Training species selection

We selected 17 clades that together cover 1,705,913 (99.4%) of the 1,716,067 eukaryotic taxa represented in the NCBI Taxonomy as of August 15, 2026. For each clade, test species were hand-picked for evaluations (Supplementary Figures 3–20). We here call these species *out-of-distribution* (OOD) for **Vipsania**, as all genomes from their entire genus were excluded from training. We then use the same automated process for each clade to find a set of diverse training species. We query all available genomes in the NCBI database that have at least an assembly status of “chromosome” (exception: “scaffold” for Amoebozoa and Rhodophyta, where too few chromosome-level assemblies were available), an assembly size of at least 2 Mb and at most 20,000 scaffolds, and that were not flagged as “atypical” by NCBI. Based on the number of available assemblies in a clade, we set the total number *N* of training species we want. We then maximize the diversity among training species for a given number of training species. As the leaves of the taxonomic tree generally have highly varying depths, we first made each subtree ultrametric, such that all leaves have the same time-distance from the root. We then use a custom script to select a set of leaves such that the subtree they induce has the maximal possible sum of branch lengths, i.e., the training genomes are maximally diverse. From this set of *N* species, we also selected up to 5 *part-of-distribution* (POD) test species that have a reference annotation available and are maximally diverse among them. The **Vipsania** Arthropoda model was trained on insects and other arthropods. The Arthropoda test species do not contain insects.

The median total genome size *M* ∈ ℕ (in bp) of all genomes in a clade is calculated. For each species in the training set of this specific clade, the genome is split into files of a set number of nucleotides each, totaling at most *M* base pairs. This reduces the overrepresentation of species with large genomes. In total, **Vipsania** was trained on 910 Gbp from 1,559 genomes across 17 clades.

According to NCBI’s taxonomy, 0.6% of eukaryotic species (9,792) fall into none of the 17 monophyletic clades for which we trained a model. We therefore trained an 18th **Vipsania** model on all 33 species with available genomes from this non-monophyletic set. The model was tested on seven species from the clades lancelets, comb jellies, hemichordates, parabasalids, xenacoelomorphs, horsehair worms and amoeboids. **Vipsania** achieved an average locus F1 score of 43.6% (Supplementary Table 22). Compared to the other clade-specific models, this performance is within the interquartile range of the average performance (41.61–59.09). Every eukaryotic species is thus covered by a **Vipsania** model, leaving no systematic gaps in applicability.

### Phylogenetic leakage

We studied whether the deep gene finders perform uniformly across the given clade or, conversely, whether they suffer from *phylogenetic leakage*, i.e. they are overfit to features of the training set and generalize poorly to distant species in the clade. For this, we measured the performance of the deep gene finders as a function of the distance between the test species and the training set. For each test species in one of the 17 clades marked in Figure 1, we selected a closest species in the taxonomic tree spanned by this species and the training set of one of the annotation tools. The proteome of this closest training species was then aligned against the test species’ proteome with DIAMOND [BRD21] and the percentage of exact matches of aligned positions was measured. Figure 5 shows a scatter plot of locus F1 values and mean percent identity to the taxonomically nearest species of the training set of each tool. A linear interpolation of these points was performed to measure the trend of annotation performance for increasing distance to the training set. The figure shows that **Vipsania** annotates species accurately even if they exhibit low similarities to those seen during pretraining. All other deep learning gene finders show a clear decrease in performance; **Helixer, ANNEVO** and **Tiberius** lose 3, 4 and 5 percentage points, respectively, for every reduction of 10 percentage points in percent identity between target and training species. The slope of the linear fit for **Tiberius, ANNEVO** and **Helixer** is significantly steeper than **Vipsania**’s; a two-sample t-test for equality of regression slopes yields *p* = 2 · 10^−4^, *p <* 10^−4^ and *p* = 0.001 respectively, for the alternative hypothesis that **Vipsania** has the smaller slope. Only clades where all compared tools have a corresponding checkpoint were included in the analyses: Arthropoda, Fungi, Insecta, Streptophyta, Vertebrata. We did not include other invertebrate clades, as test species in general were far away from the training sets of both **Helixer**’s and **ANNEVO**’s Invertebrate model and had low F1 scores.

**Figure 5.**
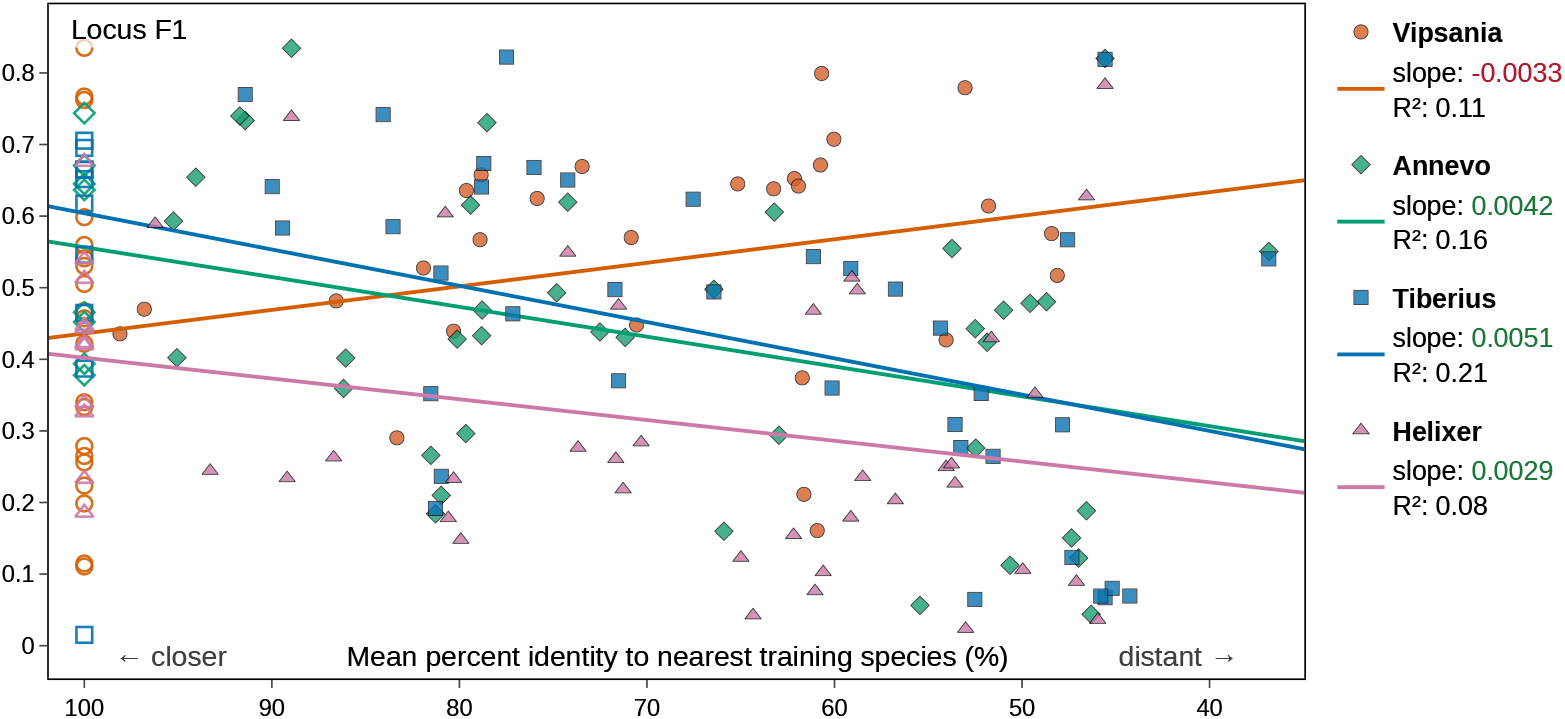
Annotation performance against test species distance to training set. The horizontal axis measures the percentage of aligned and matching positions of the proteomes of the test species with the taxonomically closest species in the training set of the corresponding model. Unfilled markers at 100% represent species included in the training set of the model. A linear regression shows the performance trend on distant species. Evaluations were conducted for 48 species across 5 major clades on all models.

Note that the training sets of **Tiberius, ANNEVO** or **Helixer** can overlap our set of test species. This may lead to major advantages that these deep gene finders might have on certain species that do not generalize to the whole clade. As an example, Figure 11a shows the performance of the Fungi deep learning models on our test species. Here, **Tiberius** has an average of 3.8 percentage points higher locus F1 than **Vipsania** on species that were included in its training set, but an average of 5.9 percentage points lower locus F1 on species that it was not trained on. Figure 11b shows the absolute number of species that overlap with the training set of any given tool. Both **Helixer** and **ANNEVO** have 4 Streptophyta species overlapping their training set, **Tiberius** includes 3 Vertebrata test species, and both **ANNEVO** and **Tiberius** include 3 Fungi test species. When also including close relatives in the training set, these numbers increase further. For example, for the annotation of *Drosophila melanogaster*, **Tiberius** can extrapolate from 6 different members of the *Drosophila* clade found in its training set. **Vipsania** excludes the *Drosophila* genus from its training set. **Vipsania** avoids the problem of overfitting to specific gene sets entirely, and can therefore give unbiased annotations regardless of species.

### BUSCO completeness score

We evaluated the BUSCO completeness score [Teg+25] of **Vipsania** and other models. Figure 6 compares the averaged scores across all 17 clades. **Vipsania** has the highest average BUSCO completeness (86.22%), followed by **Tiberius** (80.35%), **ANNEVO** (79.84%), **Helixer** (75.71%) and **GeneMark** (65.78%). **Vipsania** even surpasses the reference annotations on average by at least 1 percentage point in 9 of the 17 clades: Alveolata, Amoebozoa, Chlorophyta, Discoba, Fungi, Nematoda, Porifera, Rhodophyta and Stramenopiles. **Vipsania** exhibits high values throughout and can be trusted to find universal genes although it has not been shown examples of them during training. Note, however, that it has been argued that the BUSCO score should not be relied upon as a main metric for annotation tool evaluation [Gab+24a].

**Figure 6.**
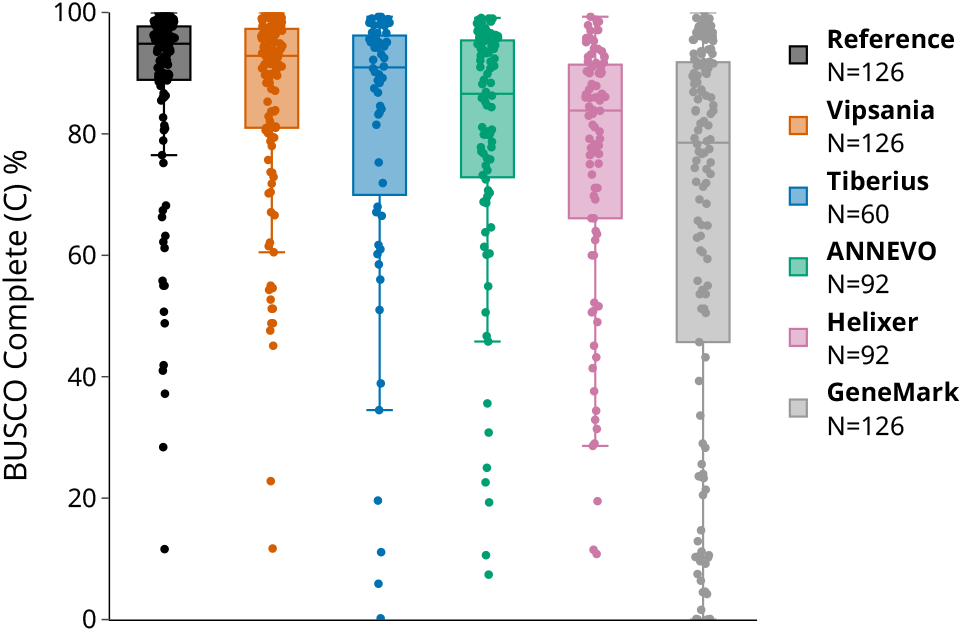
BUSCO completeness score of all tools, averaged across *N* species for each tool. All 17 clades were considered, and tools were evaluated if they had a matching model for that clade.

### Influence of repeat content

Repeats are a major driver of evolution, directly influence genome annotation [Gab+24b] and may dominate the pretraining objective of genome foundation models [ML26]. Repeat and coding region annotation are related, as repeats often make up large fractions of genomes (Figure 12), but only a small fraction of coding regions. We therefore assume that repeats are a useful source of information for deep learning gene finders. Across the clades for which all tools have a dedicated model, we plotted the percentage of repeat-masked positions in each individual chromosome or scaffold of each species against the normalized locus F1 score of an annotation in this sequence. We only considered sequences with at least 20 kb and 5 loci in the reference annotation. An exponential line *f* (*x*) = *a* · *e*^−*k*·*x*^, where *x* is the percentage of repeat-masked positions in the sequence and *f* (*x*) is fit to the locus-normalized F1 score, scaled by the fraction 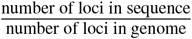 of loci in the reference annotation. Figure 12 shows that all tools deteriorate in mean performance for higher repeat content. On low repeat content sequences, only **Tiberius** performs better than **Vipsania**. However, **Vipsania** exhibits the slowest decay in annotation quality, and exceeds the other tools from just below 30% repeat content onwards.

### Non-standard genetic codes

We test whether eukaryotic species with a reported non-standard genetic code can be annotated by **Vipsania** and, at the same time, whether **Vipsania** can be trained successfully on a single genome alone. In **ANNEVO** and **Helixer**, the set of stop codons is hard-coded, and non-standard genetic codes are not supported. To this end, we train a small 2M parameter model on two test species for 500 epochs. Both *Paramecium tetraurelia* and *Tetrahymena thermophila* are known to follow the genetic code listed as translation table 6 in the NCBI taxonomy entry^1^. The change to **Vipsania** we make for a successful annotation is the removal of two stop codons (`TAG`, `TAA`) in the emission distribution of the HMM. Both codons get translated to glutamine in these species and do not serve as termination signals for genes. Figure 7a shows the training process of the standard **Vipsania** model as well as the model with the reduced set of stop codons. When the model’s internal genetic code is adapted, we see an immediate improvement in annotation quality, even in the very first epochs of training. The final locus level annotation accuracies reach above 50% (Figure 7b), while the standard model stays below 1%, both in sensitivity and precision. The adapted models converge after approximately 200 epochs, which we recommend as a general rule for single species training. This is equivalent to about 24 hours of training on an A100 GPU.

**Figure 7.**
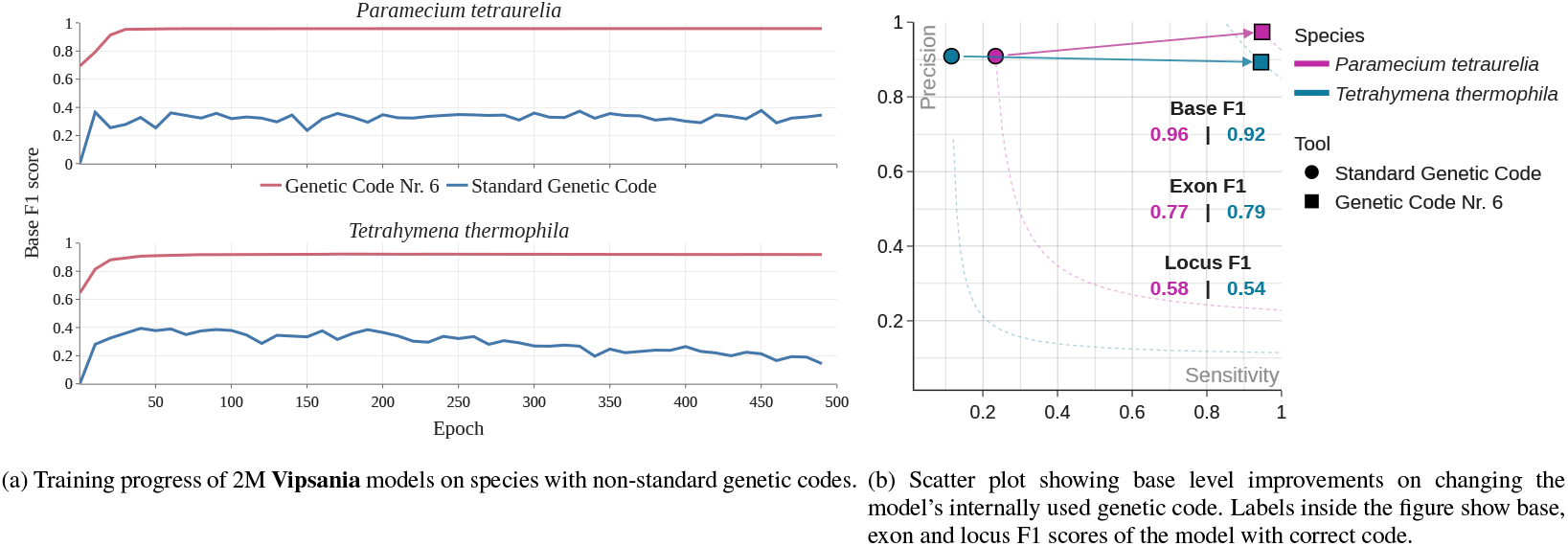
Training progress and final model annotations after completed training of two species with a non-standard genetic code.

### Predicted gene complexity

The profoundly different loss functions of supervised gene finders on the one hand and the masked language loss of **Vipsania** on the other hand could lead to different relative strengths with regard to gene complexity. Also, when multiple tools are used, e.g. with a combiner [Gab+21], a high complementary sensitivity of one tool with respect to another is desirable. As correctly predicting large genes is a particular challenge, we compared complementary accuracies as a function of gene size: Let *V* denote the set of genes found by **Vipsania** but not by a particular one of the four other tools. “Found” here means that one splice form of the reference gene structure is predicted by **Vipsania**, but none of its splice forms was predicted by the other tool. Define *W* as the set of genes found by that other tool but not by **Vipsania**. Figure 8 shows the difference |*V*| − |*W*| per other tool, stratified by either total gene length or length of the coding region, across 48 species in the 5 common clades as mentioned in prior analyses. Against every other gene finder, **Vipsania** correctly predicts genes that the other tool misses. As genomic gene span or total coding length increases, **Vipsania** gains an advantage over **Helixer, ANNEVO** and **GeneMark**, and catches up to **Tiberius**. In relative terms, long genes are a particular strength of **Vipsania**, even though it had to learn the gene boundaries from scratch, without any supervision. Clade specific results can be found in Supplementary Figures 1 and 2.

**Figure 8.**
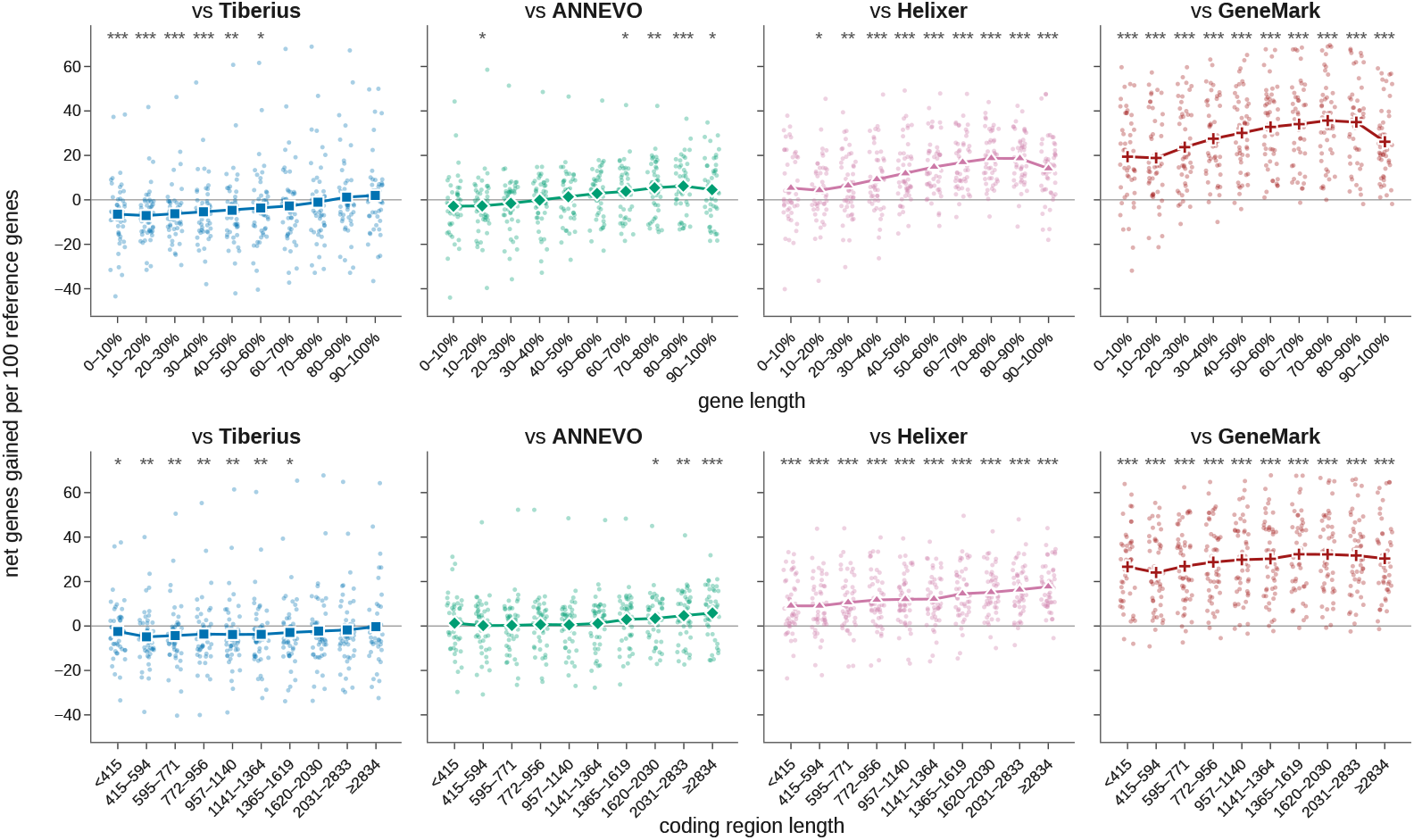
Difference of unique correctly predicted genes by **Vipsania** and another annotation tool. Length of a gene and the total coding region were used as gene complexity metrics. For gene length, the shown deciles are clade-specific. A Wilcoxon signed-rank test is performed and a bin is marked by one, two or three stars * if its p-value is less than 0.05, 0.01 or 0.001.

### Spliced loss

We deploy an extension to the default BERT-like loss in all **Vipsania** models except for the Vertebrata clade. This *spliced loss* is defined in (2). It increases the annotation quality of **Vipsania** drastically. Figure 9a shows the first 500 epochs of two **Vipsania** trainings, one with the spliced loss and one without, and their annotation quality on four test species of the Insecta clade. The spliced loss is activated at epoch 100, raising its weight factor *γ* from 0 to 0.05 over 5 epochs. This leads to an immediate increase in base F1 score, caused predominantly by an increase in precision, and drives the model towards more accurate annotations that it would not find with just the cross-entropy loss.

**Figure 9.**
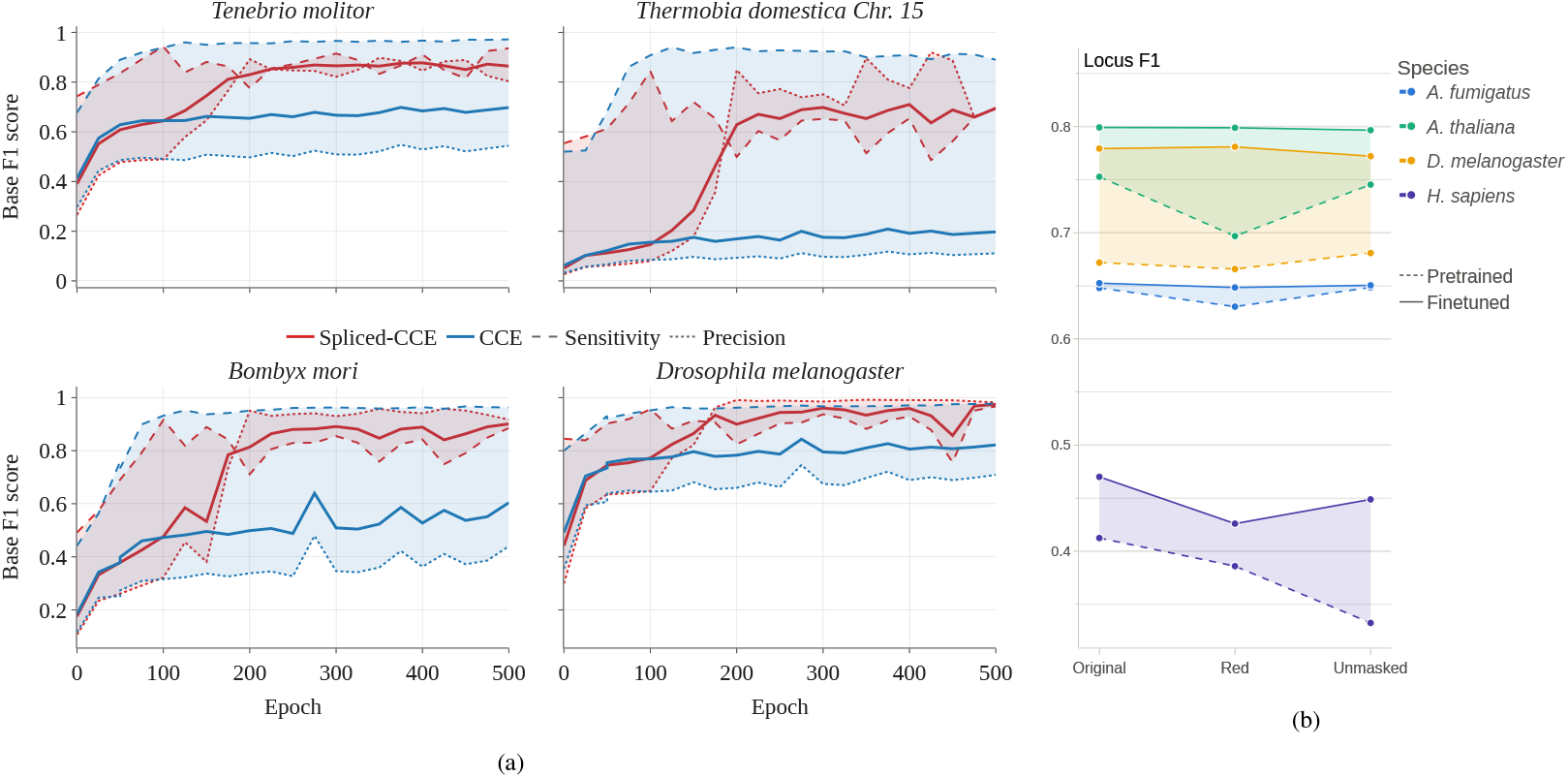
**(a)** Annotation quality for four species during two **Vipsania** Insecta training runs. One model uses ℒ_spliced-CCE_ and the other ℒ_CCE_. An annotation is produced every 25 epochs. The weight of the spliced loss in the red model increases from *γ* = 0 after epoch 100; see (2). **(b)** Stability of **Vipsania** finetuning across different repeat-masking approaches. Genomes are either unmasked, masked by **Red**, or left unchanged from their NCBI records.

### Finetuning

Here, “finetuning” refers to a continued training of a **Vipsania** model on a specific target genome. This is still an unsupervised training and does not use external gene structure data. To quantify the improvement by finetuning, Figure 13 shows the mean annotation locus sensitivity, precision and F1 increase over all test species per clade due to finetuning. Finetuning improves annotation quality on all clades with the exception of Arthropoda. An average F1 increase of 7.5% or more can be found in the clades Rhodophyta, Spiralia and Vertebrata. Finetuning on single genomes takes about 160 minutes on an A100 GPU with 80 GB of memory. For annotations across several closely related species, we recommend finetuning once on the set of genomes and then using this model to annotate multiple genomes. **Vipsania** is faster at the actual inference than the 4 other gene finders. As an example, annotating the genome of *Drosophila melanogaster* (144 Mb) with **Vipsania** takes just under 6 minutes on an A100 GPU. For more details on running times, we refer the reader to Supplementary Table 4.

### Repeat masking

We trained **Vipsania** on the most recent genome submissions to the NCBI database for our selected training species. The repeat annotations of these genomes are conducted using a variety of different approaches. Therefore, repeat masking quality may vary, which directly influences the performance of a **Vipsania** annotation. However, this effect can be dampened when finetuning on the specific genome version before annotating. We conducted analyses on 4 test species, whose genomes we either left unchanged from the NCBI submissions, stripped of their repeat masking information, or masked with the fast repeat masker **Red** [Gir15]. Figure 9b shows how finetuning can mitigate the effects of differently repeat-masked genomes before annotating. While the performance of a pretrained **Vipsania** model can heavily change depending on the species and its repeat masking, the finetuned models often arrive at a similar locus F1 score. If it is unclear whether a given genome has a high-quality repeat masking, we suggest unmasking the genome and finetuning before annotating.

### Pretraining

Recently, the effectiveness of pretraining has been challenged for genome foundation models [Vis+24]. Therefore, we analyzed whether the clade-wide training of **Vipsania** is actually needed to achieve the best annotation accuracy. On three test species from different clades, we trained 10M parameter models for 250 epochs on each genome and compared the resulting accuracy to the one from the corresponding clade model. Figure 10 shows the training progress of the 10M model and the results of the pretrained and pretrained+finetuned **Vipsania** model. On all test genomes, pretraining helps to model an annotation, although the difference is much more pronounced for larger genomes. Training the 10M parameter model took about 26 hours on an A100 GPU with 80 GB of memory, while the pretrained 25M parameter models trained for 10 days to reach 1000 epochs.

**Figure 10.**
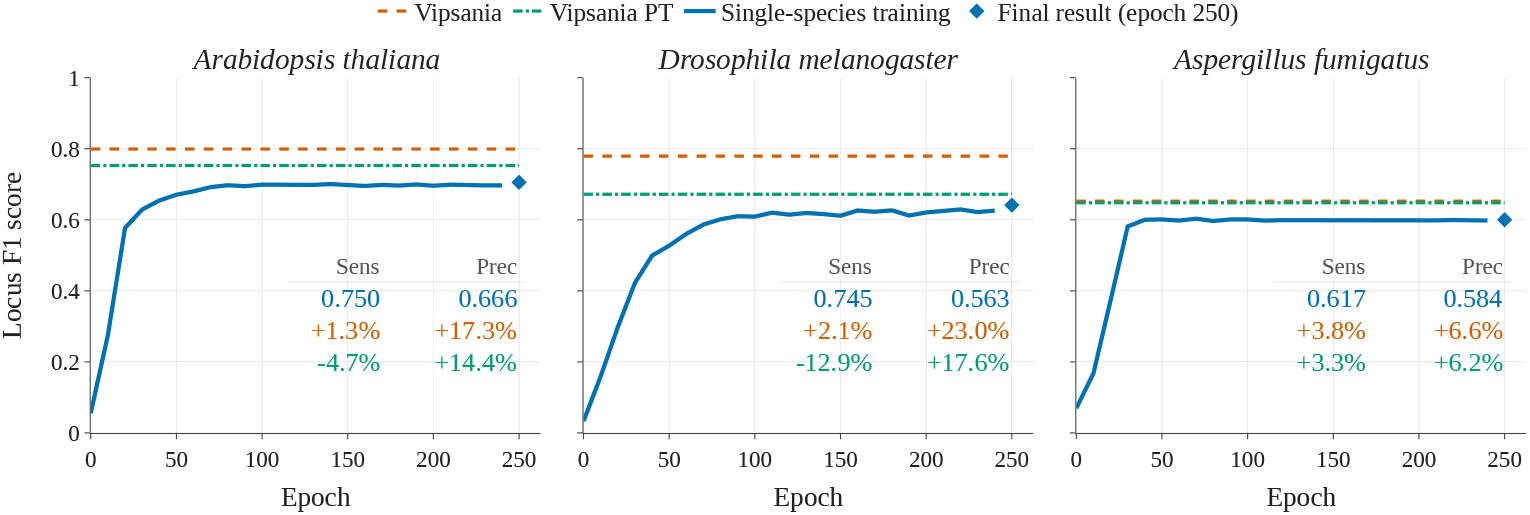
Comparison of locus level performance of single species training (blue), clade-wide pretraining only (Vipsania PT, green) and pretraining+finetuning (Vipsania, orange).

## Discussion

We release the first unsupervised deep gene finder, **Vipsania**. Throughout training, the model never saw examples of genes. Instead, it learned to reconstruct nucleotides from their surrounding context, with a differentiable HMM layer providing an inductive bias towards learning gene structures. Our demonstration — that unsupervised large-scale training can produce precise segmentations — qualifies the opposite conclusion recently made by Shmelev et al. [Shm+26] for general foundation models. The released 25M parameter **Vipsania** models were trained on close to 1 Tbp of genome sequence systematically sampled across the eukaryotic tree. **Vipsania**’s annotation quality matches or exceeds that of the supervised tools **Tiberius, ANNEVO** and **Helixer**, as well as that of the unsupervised gene finder **GeneMark**, on 38 of 60, 82 of 92, 91 of 92 and 123 of 126 test species, respectively. For clades for which supervised models provide no parameters, **Vipsania** is the only applicable modern deep learning gene finder, and it delivers strong performance. Its reach extends even to species with a non-standard genetic code: since no annotated relative is required, adapting **Vipsania** amounts to changing the set of stop codons in the emission distribution of the HMM and running a single-species training, whereas **ANNEVO** and **Helixer** have the standard code hardcoded.

Supervised gene finders require training annotations and thus suffer from four limitations by design. First, there may be insufficiently many or insufficiently diverse annotations available for the successful training of a deep gene finder. **Helixer, ANNEVO** and **Tiberius** were trained on 30–128, 21–306 and 9–306 genomes per clade, respectively, and such training sets are not available for the less well-studied clades of eukaryotes. Second, for all but a few exceptionally well-studied species (such as human, mouse, fruit fly and *Arabidopsis*), the available annotations are the product of evidence-based pipelines, which make a substantial number of errors [Gab+24a]. Third, the annotations available for training supervised methods are heterogeneous across species and were produced with different tool combinations at different times. It is not clear how to decide which annotations should be included in the training and which may decrease performance because they ‘teach’ the model too many, or particularly consequential, annotation mistakes. Fourth, the set of training gene structures is likely to introduce a bias towards certain species and potentially towards certain genes. We show that one such bias is phylogenetic feature leakage, introduced by the selection of the training set. Supervised methods extrapolate poorly to test species that are less similar to species from the training set. **Vipsania** avoids these problems by dispensing with supervision altogether.

Many tasks general eukaryotic genome foundation models are built for — promoter and regulatory element design, variant effect prediction, expression estimation — presuppose knowledge of proteincoding gene structure. A promoter is defined relative to a transcription start site; the consequence of a variant depends on whether it falls in a coding exon, at a splice site or in an untranslated region; expression is quantified per transcript. Structural annotation is thus not one downstream task among many but the coordinate system in which the others are expressed. We have shown that our HMM layer learns the gene structures already at an intermediate layer so that downstream layers can use gene structure information. We therefore expect that this layer will be broadly useful for genome foundation models.

The availability and suitability of tools for the annotation of protein-coding genes depend on the evidence available for the target species or even the target gene. In evidence-rich situations with highquality annotations of related species and diverse, deeply sequenced RNA-seq libraries from multiple tissues, conditions and developmental stages, **Vipsania** is not expected to outperform evidence-based pipelines on average but may be useful to complement their annotations with genes that have insufficient expression or lack homology evidence. For the bulk of species with limited available evidence, and with the exception of vertebrates, where supervised tools have particularly good training data, **Vipsania** can provide a state-of-the-art annotation. Due to its minimal input requirements and broad applicability across eukaryotes, **Vipsania** is the first genome annotation tool that can be used to generate annotations for virtually all eukaryotic organisms at a quality level that is competitive with state-of-the-art supervised methods that have narrower applicability. We expect that a combination of RNA-seq with deep *ab initio* gene prediction may become the best practice for general annotation of protein-coding genes.

## Methods

### Model architecture

Our deep learning model **Vipsania** consists of several layers connected by an uninterrupted residual stream with single base-pair resolution throughout, see Figure 2. The architecture is reminiscent of the original Transformer [Vas+17]. However, we replace attention with the Linear Recurrent Unit (LRU) [Orv+23], use a position-wise SwiGLU operation [Sha20] and normalize the input to each layer with LayerNorm [BKH16]. The most important component of our model is a differentiable HMM layer [BS22; Gab+24b] that allows it to infer a genome annotation while being trained fully unsupervised. In contrast to existing deep gene finders (**Helixer, Tiberius, ANNEVO**), **Vipsania** does not use convolutional layers.We use 16 layers with a residual stream of size 320. The HMM is placed after the 8th layer. The resulting model has 25M parameters.

We adapt the LRU specified in [Orv+23] by removing the exponential activation from the parameterization of the phase *θ* of the eigenvalues in the state transition matrix *A*. Following [BG25], we learn the LRU’s initial state by averaging across time steps and training epochs. Two LRUs with separate parameters are trained per layer to model the forward and backward strand of the DNA. Both directions are concatenated, ReLU-activated and linearly projected to fit the dimension of the residual stream. The LRU was implemented efficiently using a parallel scan [Ble90], which itself is compatible with TensorFlow’s [Mar+15] just-in-time (**jit**) compilation, allowing for a large speedup of the whole model.

### HMM layer

The core component of **Vipsania** is a differentiable HMM layer designed for gene finding. The state space is closely related to the one used by **Tiberius** [Gab+24b], with the exception that we extend the number of states to 18 by using two intron states per phase (see Figure 2). This design choice allows the model to learn short and long introns. We additionally use two copies of this HMM with shared parameters to simultaneously model the forward and backward strands of the DNA. The HMM is implemented using the packages `hidten`^2^ and `bricks2marble`^3^, which are compatible with TensorFlow’s **jit** compilation.

The HMM layer takes three inputs, all encoded as matrices. The nucleotide sequence is encoded as overlapping triplets to allow the state at each time step to see the two preceding (left) and two following (right) nucleotides. Let *X*_left_ ∈ ℝ^*T ×*64^ and *X*_right_ ∈ ℝ^*T ×*64^ be the one-hot encodings of triplets, and let 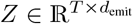 be a row-stochastic encoding over a learned emission alphabet of size *d*_emit_. Each row of *Z* is a probability distribution over the emission alphabet letters, predicted from the residual stream (see below). Let *S* ∈ ℝ^*T ×*18^ be a one-hot encoding of an HMM state sequence, and let *A* ∈ ℝ^18*×*18^ be the transition matrix, *B*_left_ ∈ ℝ^18*×*64^, *B*_right_ ∈ ℝ^18*×*64^ the triplet emission matrices and 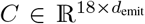 the emission alphabet matrix. The triplets and the emission alphabet symbols are assumed to be conditionally independent given the state at each time step. The HMM defines a probability distribution over *S, X*_left_, *X*_right_ and *Z*:

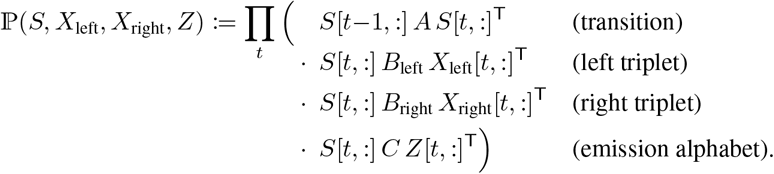

During training, the HMM layer computes the posterior state probabilities

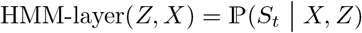

at each time step *t* using a differentiable and vectorized forward–backward algorithm. The posterior state probabilities are then added to the residual stream after a convolution with a learned kernel of width 9.

Unlike *X*, the matrix *Z* is not directly observed. Let 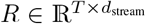 be the residual stream feeding into the HMM layer. We place a learned map *Z* = softmax(*RW* ) with a parameter matrix 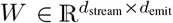 before the HMM, where the softmax is applied row-wise.

For the final **Vipsania** model, we chose *d*_emit_ = 160. The row-stochastic matrices *A* and *C* are learned, while *B*_left_ ∈ ℝ^18*×*64^ and *B*_right_ ∈ ℝ^18*×*64^ are static and contain hand-picked values to enforce canonical splice site patterns, start and stop codons and to prevent in-frame stop codons within exons.

While **Tiberius** does not adapt the parameters of the HMM during training, we observed that it is crucial to do so in the unsupervised case. **Vipsania** uses learnable parameters in the emission matrix, the transition matrix and the start distribution of *S*[0, :] and excludes them from weight decay throughout training. We found that a parameter sharing of the rows corresponding to noncoding states in the emission matrix can help to model long introns.

### Loss

The main training objective for **Vipsania** is to fill in missing nucleotides that have been masked in a target genome. In the following, we will refer to this as token-masking to avoid confusion with repeat-masking. We use the **BERT**-like [Dev+19] categorical cross-entropy loss

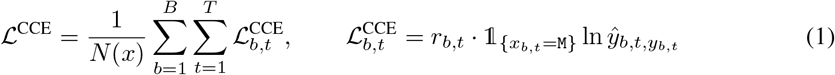

over batch size *B* and context length *T*, where *x*_*b,t*_ is the input and *y*_*b,t*_ the target nucleotide, and *ŷ*_*b,t*_ is the predicted distribution of the model. The indicator function 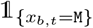 is 1 if the position *t* of sequence *b* is token-masked and otherwise 0. The additional term *r*_*b,t*_ is set to 10^−4^ if *y*_*b,t*_ is in a repeat-masked region and is otherwise 1. The loss is normalized by *N* (*x*), the number of token-masked positions in a batch *x*. We use a maximum of 5% token-masked base pairs for *T* = 20,000, all of which are replaced by the masked token `M`, as we did not see any annotation performance benefits of varying replacement tokens as mentioned in [Dev+19]. We do not token-mask ambiguous bases, represented by `N`.

The loss (1) indirectly incentivizes the model to learn the structure of genes via a higher accuracy of predicting the right nucleotides at gene-structure-dependent sites such as at exon boundaries and third bases in a codon. The use of loss (1) is by itself sufficient for the HMM to learn to produce reasonable full genome annotations after training. However, the annotation quality can be significantly boosted with an extension that considers current beliefs of the model about the location of exon boundaries. Let P_splice_(*x*_*b*_)_*t*_ be the probability — estimated by the HMM — of the current position *x*_*b,t*_ being an exon boundary state (start or end of a gene or a splice site, see Figure 2). Then, the augmented loss is

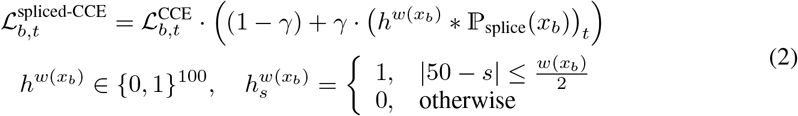

where * is the convolution of the probability of the splice site and a rectangular kernel on the time axis. When *γ >* 0, incorrect nucleotide predictions near exon boundary positions are penalized more. The convolution then also incorporates the *w*(*x*_*b*_) ∈ ℕ_*<*100_ positions around the boundary, which are put under the same pressure. This width *w* is a crucial parameter, which has an optimal value that changes for different clades and even species. We therefore calculate it context-dependently as

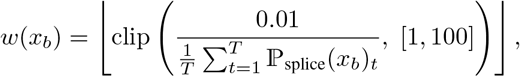

with a minimum value of 1 and a maximum value of 100. The factor 0.01 is chosen to work across all clades.

Note that the loss extension (2) does not violate the unsupervised criterion, as the probability of a splice site is directly taken from the HMM that is trained alongside all other components only on the objective of predicting nucleotides. As annotation quality is low at the start of any **Vipsania** training, we only increase *γ* = 0 after 100 warmup epochs to *γ* = 0.05 in a linear ascent over 5 epochs.

### Finetuning

The model checkpoint for a clade has been trained, typically for 1000 epochs, on a large number of species. A major advantage of an unsupervised training is the possibility of finetuning the model on a species of interest before generating an annotation. In our experiments we observed that such a specialized, short training on a single genome increases the annotation quality drastically. Finetuning consists of training the model for 10 additional epochs with a lower learning rate than in the pretraining stage. The spliced loss (2) is active, starting at epoch 0 with *γ* = 0.05. Large genomes often exhibit a higher quantity of repeat regions, which did negatively impact finetuning. Therefore, for genomes larger than one billion base pairs, we slightly change the data loader during finetuning to only train on sequences with less than 25% repeat-masked bases.

### Viterbi re-predictions

During training, the HMM layer in **Vipsania** outputs posterior probabilities. During inference, however, we use the Viterbi algorithm to generate an annotation sequence *s* for the (unmasked) input genome *x*. During inference, **Vipsania** infers state sequences independently and in parallel on nonoverlapping tiles of length 200,000, which is also the inference context length. Subsequently, incompatibilities at the seams between tiles are identified and fixed as follows. Let *s*^(*l*)^ and *s*^(*r*)^ be two predictions of the HMM for two consecutive genome regions *x*^(*l*)^, *x*^(*r*)^. If 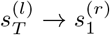 is an invalid state transition (we only allow intergenic regions or introns crossing prediction borders), then we make a re-prediction *s*^(*c*)^ spanning 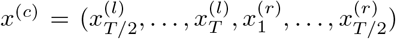 and replace the central part of the sequence. For that, we find indices

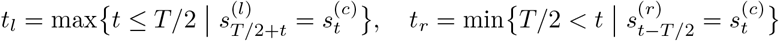

and replace 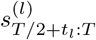 by 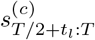 and 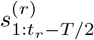 by 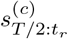 before merging. If these matching positions *t*_*l*_, *t*_*r*_ cannot be found, we set the surrounding area from the last to the next complete gene to an intergenic region.

### Vertebrata clade training

The complexity of Vertebrata genomes and, more specifically, their increased average gene length lead to difficulties for **Vipsania** regarding annotation performance. While supervised methods can learn structural differences between long introns and intergenic regions by extrapolating from given genes in the training set, this appears to be more difficult in an unsupervised setting without any annotation labels.

One reason for bad annotation performance in early experiments on Vertebrata genomes was the discrepancy between the expected lengths of intergenic regions and introns, modeled by the HMM. More specifically, let 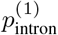 and 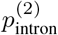 be the probabilities of staying in the two intron states `Ik1` and `Ik2`, which are shared parameters across indices *k*, see HMM in Figure 2. Let *p*_intergenic_ be the probability of staying in the intergenic state. While 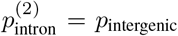 at initialization, they drift apart during training, which makes the prediction of long genes theoretically infeasible, even for small differences. We therefore include an auxiliary loss that penalizes large differences in the expected lengths of introns and intergenic regions modeled by the transition matrix of the HMM.

We use the parameter regularization

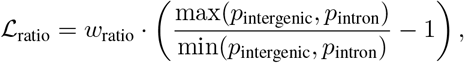

where *w*_ratio_ is a hyperparameter. Additionally, we increase the number of epochs trained to 3000.

## Extended Data Figures

**Figure 11.**
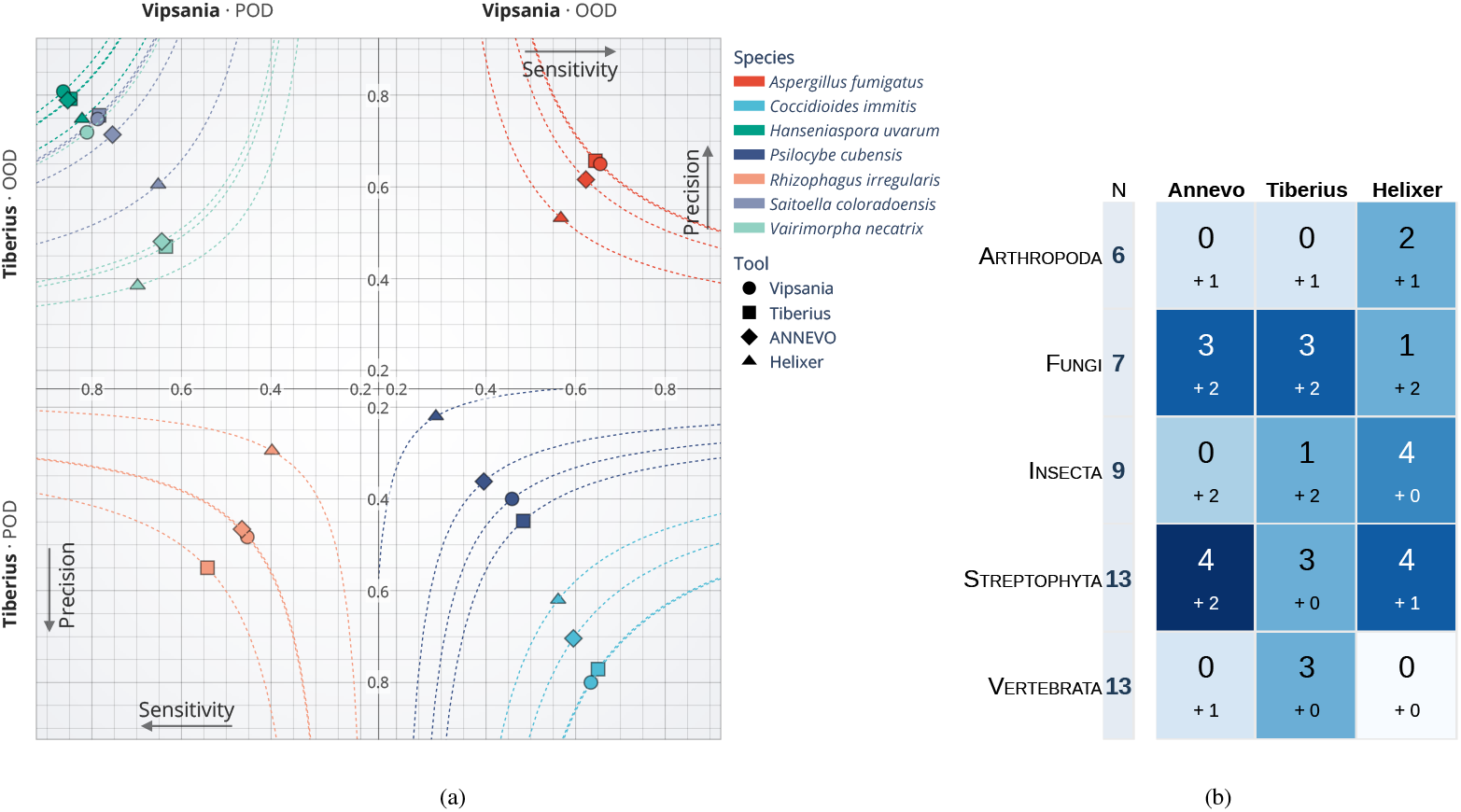
**(a)** Locus-level metrics for Fungi test species in four quadrants. A species is assigned to a quadrant based on whether it is part of (POD) or outside (OOD) a method’s training set. For example, the first row shows all species outside the training distribution of **Tiberius.** Because **Tiberius** and **ANNEVO** use the same fungi training set, this classification applies to both models. **(b)** Overlap of our test species with the training sets of other tools. The large number includes exact species name matches, while the “+ x” is the number of test species where a representative of the same clade exists in the training set.

**Figure 12.**
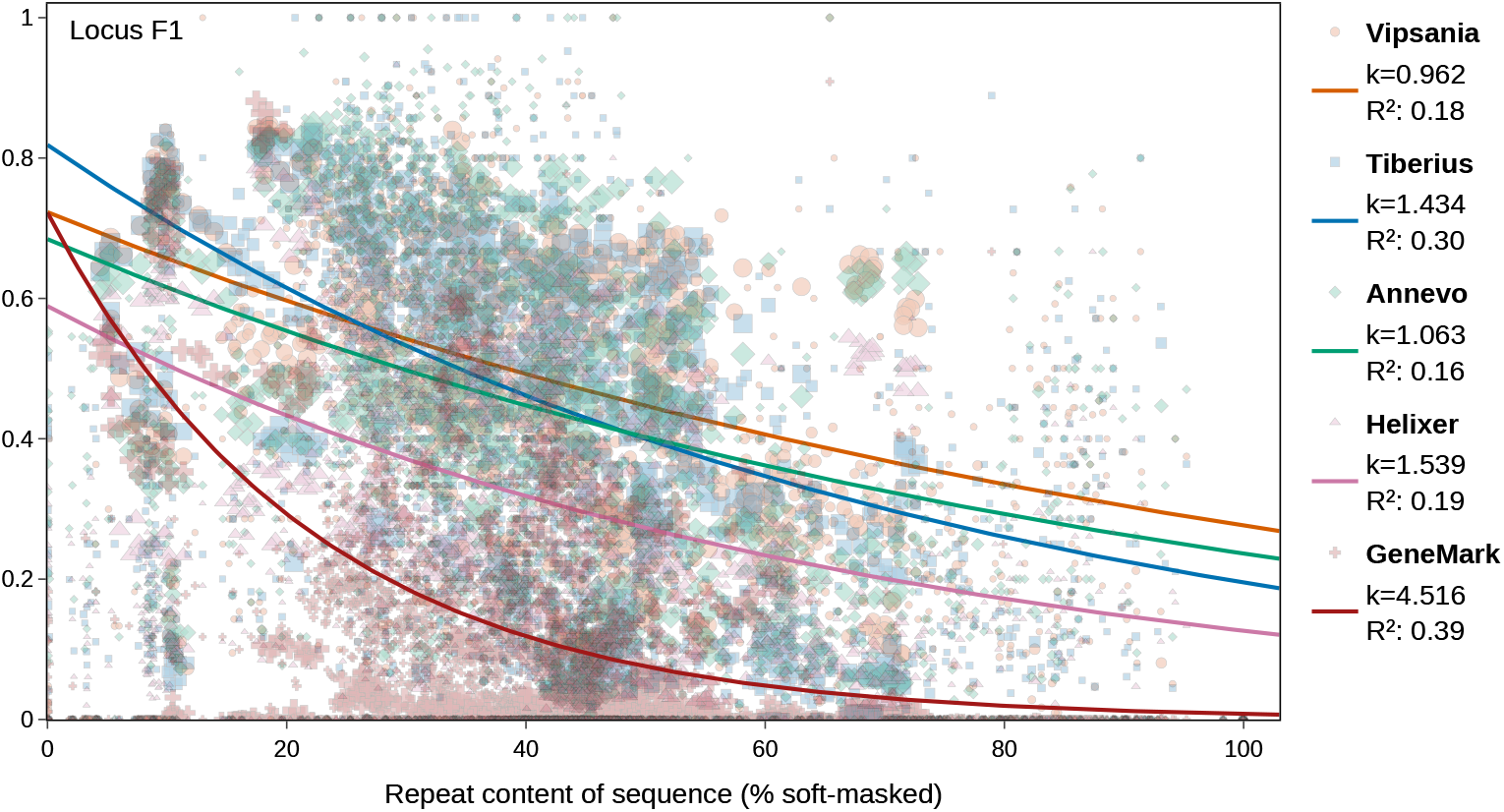
Annotation performance per chromosome/scaffold and its repeat content. Evaluation is spanning 2,869 sequences across 48 test species in the five major clades for which all tools are available.

**Figure 13.**
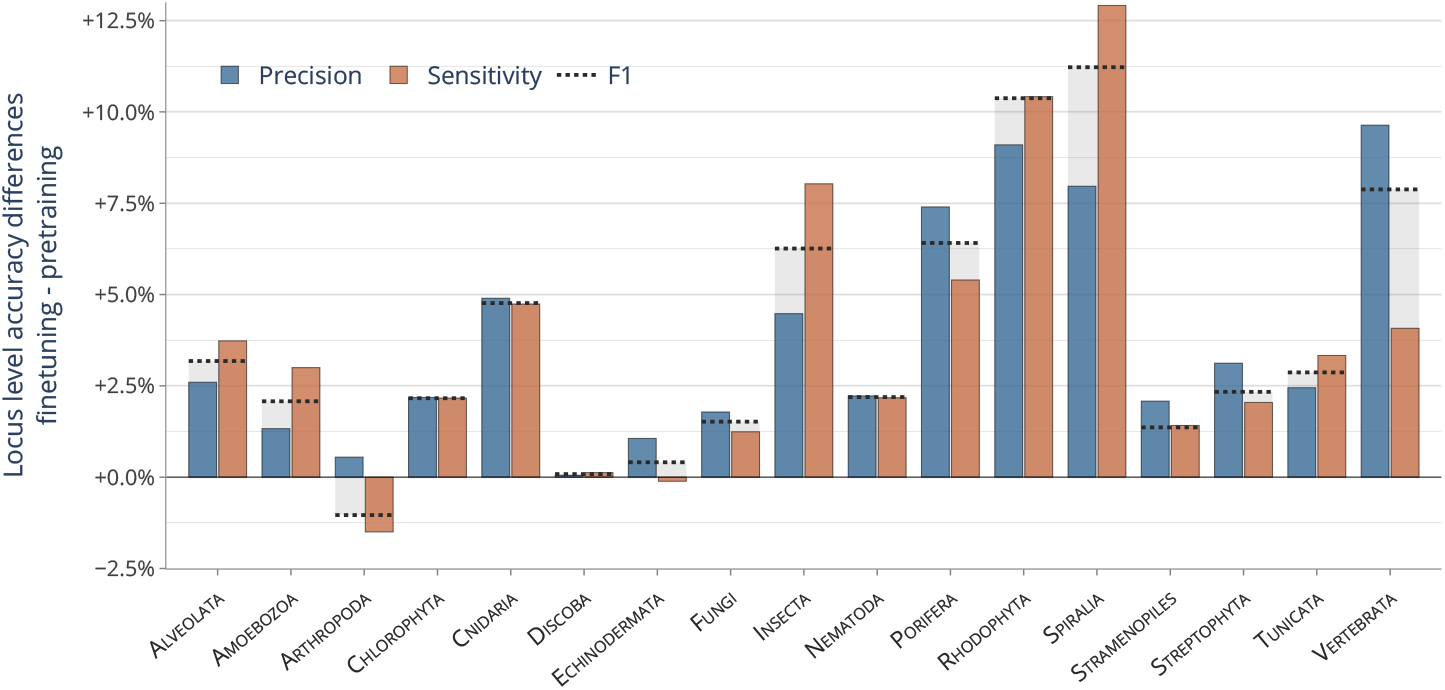
Locus sensitivity and precision gains of using species-specific finetuning instead of a raw pretrained **Vipsania** model. Shown are the means over all test species in each clade. The shown F1 value is the mean of F1 values across species.

## Acknowledgements

We thank Tomáš Brůna for the idea of comparing sensitivities to repeat contents. This work was funded by the Deutsche Forschungsgemeinschaft (DFG, German Research Foundation) – STA 1009/17-1.

## Footnotes

1 The Ciliate, Dasycladacean and Hexamita Nuclear Code

2 https://gaius-augustus.github.io/hidten-docs/

3 https://github.com/gaius-augustus/bricks2marble

