## Supplementary Material for "Vipsania: Unsupervised Deep Gene Finding"

Richard Krieg<sup>1</sup> Felix Becker<sup>1</sup> Stepan Saenko<sup>1</sup> Joscha Diehl<sup>1,§</sup> Mario Stanke<sup>1,§</sup>

<sup>1</sup>Universität Greifswald, Institut für Mathematik und Informatik
{richard.krieg, felix.becker, stepan.saenko, joscha.diehl, mario.stanke}
@uni-greifswald.de

§ joint senior authors

### 1 Tool usage

#### 1.1 Annotation tools

##### Vipsania

```
annotate.py \
  <model> \
  <genome.fa> \
  -o <output.gff3> \
  --finetune
```

The <model> choices for Vipsania, e.g. 58hsuobw for insects, as well as for Tiberius, ANNEVO and Helixer are in Table 2. For runs of **Vipsania** without finetuning, the flag `--finetune` has to be dropped from above command.

##### Tiberius

```
tiberius.py \
  --genome <genome.fa> \
  --model_cfg <model> \
  --min_genome_seqlen 1 \
  --out <output.gff3>
```

##### ANNEVO

```
annotation.py \
  --genome <genome.fa> \
  --model_path <model.pt> \
  --output <output_all.gff3> \
  --threads 64

grep -P "CDS\t" <output_all.gff3> > <output.gff3>
```

ANNEVO recently released a “boundary-aware” model type, that has to be executed differently:

```
annotation.py \
  --genome <genome.fa> \
  --model_path <model.pt> \
  --output <output_all.gff3> \
  --threads 64 \
  --num_classes 15 \
  --boundary-aware \
  --
```

```
--min_intron_length 20

grep -P "CDS\t" <output_all.gff3> > <output.gff3>
```

### Helixer

```
Helixer.py \
  --fasta-path <genome.fa> \
  --lineage <model> \
  --batch-size 16 \
  --gff-output-path <output_all.gff3>
grep -P "CDS\t" <output_all.gff3> > <output.gff3>
```

### GeneMark

```
gmes_petap.pl \
  -ES \
  --soft_mask auto \
  --work_dir <output_all.gtf> \
  --cores 64 \
  --v \
  --sequence <genome.fa>

awk 'BEGIN{FS="\t";OFS="\t"}!/^#/_{sub(/_./,_,_)}1' \
  <output_all.gtf> > <output.gtf>
```

### 1.2 Evaluation tools

#### gffcompare

All genome annotation files produced with the commands of Section 1.1 are compared to a reference annotation with the following execution of **gffcompare** (version 0.12.6) [PP20]. The reference .gtf file was filtered to only contain CDS entries. Notably, this also removes UTR regions from the annotation, which are not predicted by **Vipsania**. We extract exon and locus level metrics from the **gffcompare** output. None of the gene finders predict alternative isoforms. The locus level metric counts any match of a reference isoform to a predicted one as a correctly predicted gene.

```
gffcompare \
  --strict-match \
  -e 3 \
  -T \
  -o <gffcompare_output> \
  -r <reference_cds.gtf> \
  <output.gff3> # or: <output.gtf>
```

#### diamond+blastp

The evaluation of extrapolation performance of tools on species not existing in their training set required aligning each test species proteome to the taxonomically closest training proteome. We used the following **diamond** (version 2.1.9) [BRD21] commands to build the database from the training species proteome and comparing the two proteomes.

```
diamond makedb --in <train.faa> -d <workdir>/train_db --quiet
```

```
diamond blastp -q <test.faa> -d <workdir>/train_db \
  --very-sensitive \
  -e 1e-05 -k 1 -p 8 \
  --quiet \
  -f 6 qseqid sseqid pident length qcovhsp evaluate bitscore \
  -o <workdir>/hits.diamond.tsv
```

### BUSCO

For each tool’s annotation, we used **gffread** [PP20] to extract the proteome of a species. Then, we ran **BUSCO** (version 6.1.0) [Teg+25] for a predefined database, which was selected per clade, see Table 1.

```
busco -m proteins \
-i <test.faa> \
-l <lineage-database> \
-o reference \
--out_path work/busco \
-c 32 \
--download_path busco_downloads \
--offline
```

Table 1: BUSCO odb12 lineage dataset per clade.

| Clade | odb12 dataset |
| --- | --- |
| ALVEOLATA | alveolata_odb12 |
| AMOEBOZOA | eukaryota_odb12 |
| ARTHROPODA | arthropoda_odb12 |
| CHLOROPHYTA | chlorophyta_odb12 |
| CNIDARIA | metazoa_odb12 |
| DISCOBA | eukaryota_odb12 |
| ECHINODERMATA | metazoa_odb12 |
| FUNGI | fungi_odb12 |
| INSECTA | insecta_odb12 |
| NEMATODA | nematoda_odb12 |
| PORIFERA | metazoa_odb12 |
| RHODOPHYTA | eukaryota_odb12 |
| SPIRALIA | metazoa_odb12 |
| STRAMENOPILES | stramenopiles_odb12 |
| STREPTOPHYTA | embryophyta_odb12 |
| TUNICATA | metazoa_odb12 |
| VERTEBRATA | vertebrata_odb12 |

#### 1.3 Model selection

Table 2 shows the models used for each gene finder per clade. For **Vipsania** the eight letter alphanumeric code identifies the checkpoint and is to be specified on the command line. For VERTEBRATA, the best suited model was used for **Tiberius** and **ANNEVO**. That is, either a specialized Mammalia model or a general Vertebrate checkpoint. The Mammalia model was used for *Homo sapiens*, *Mus musculus* and *Delphinus delphis*. For ARTHROPODA, the ANNEVO Invertebrate checkpoint performed better than the ANNEVO Insecta checkpoint.

Table 2: Model names used for ab-initio gene finders across clades.

| Clade | Vipsania | Tiberius | ANNEVO | Helixer |
| --- | --- | --- | --- | --- |
| ALVEOLATA | sd2zcj7u | — | — | — |
| AMOEBOZOA | ezhpj2qm | — | — | — |
| ARTHROPODA | ihe1jk30 | insecta | Invertebrate | invertebrate |
| CHLOROPHYTA | faeijtnk | chlorophyta | — | — |
| CNIDARIA | b5vtieo0 | — | Invertebrate | invertebrate |
| DISCOBA | gcra9d9y | — | — | — |
| ECHINODERMATA | sx9zjl7p | — | Invertebrate | invertebrate |
| FUNGI | fh1kg88z | fungi | Fungi | fungi |
| INSECTA | 58hsuobw | insecta | Insecta | invertebrate |
| NEMATODA | zspca2rb | — | Invertebrate | invertebrate |
| PORIFERA | 7bjiexcu | — | Invertebrate | invertebrate |
| RHODOPHYTA | 6kmw3wme | — | — | — |
| SPIRALIA | v5ej8oyt | — | Invertebrate | invertebrate |
| STRAMENOPILES | r6p9z9jw | diatoms | — | — |
| STREPTOPHYTA | j9m0cdmk | angiosperms | Magnoliopsida | land_plant |
| TUNICATA | hcehc7ff | – | Invertebrate | invertebrate |
| VERTEBRATA | etb1go6q | vertebrate / mammalia | Vertebrate_other / Mammalia2 | vertebrate |
| Missing eukaryotes | cg6grhms | – | – | – |

Table 3 lists the number of epochs each **Vipsania** model was trained and how many species it was trained on.

Table 3: Model names used for ab-initio gene finders across clades.

| Clade | Number of epochs | Training species |
| --- | --- | --- |
| ALVEOLATA | 1000 | 30 |
| AMOEBOZOA | 1000 | 15 |
| ARTHROPODA | 1000 | 200 |
| CHLOROPHYTA | 1000 | 26 |
| CNIDARIA | 1000 | 82 |
| DISCOBA | 1000 | 20 |
| ECHINODERMATA | 1000 | 52 |
| FUNGI | 1000 | 200 |
| INSECTA | 1000 | 200 |
| NEMATODA | 1000 | 73 |
| PORIFERA | 1000 | 75 |
| RHODOPHYTA | 1000 | 20 |
| SPIRALIA | 1000 | 200 |
| STRAMENOPILES | 1000 | 32 |
| STREPTOPHYTA | 1000 | 200 |
| TUNICATA | 1000 | 34 |
| VERTEBRATA | 3000 | 200 |
| Missing eukaryotes | 2000 | 33 |

### 2 Experiments

#### 2.1 Predicted gene complexity

Let $V$ denote the set of genes found by **Vipsania**, but not by a particular other tool $T \in \{\text{Tiberius, ANNEVO, Helixer, GeneMark}\}$. Define $W$ accordingly as the set of genes correctly predicted by $T$ but not by **Vipsania**. We create these sets by using the .tmap file created by **gffcompare** evaluated on the reference annotation and each tool. Specifically, we use a custom solution based on **gffcompare** that is tolerant to differences in GFF conventions, that include the stop codon in the CDS or not. The difference $|V| - |W|$ is then plotted against gene length or CDS length. For the genomic span of a gene (= distance between start and stop codon in the genome), we took deciles for each clade separately, and assigned the quantile of each point in the multi-clade plot based on the clade-specific quantile. For CDS length, we made the simplified but reasonable assumption that the length of the coding (and therefore the protein) sequence is similarly distributed across clades, which lead us to use global deciles as a segmentation. Data is normalized by the total number of observations for each quantile separately. For plotting, we connect the mean of all points in each bin to visualize the trend.

We provide the clade-specific results in Figure 1 and 2.

#### 2.2 Annotation results

Table 4: Annotation wall time in minutes for all used tools across 3 diverse species. Vipsania finetuning timings exclude the actual annotation.

| Species | Vipsania finetuning | Vipsania annotation | ANNEVO | Tiberius | Helixer | GeneMark |
| --- | --- | --- | --- | --- | --- | --- |
| *Homo sapiens* | 158.6 | 92.1 | 115 | 93.2 | 838 | 338 |
| *Drosophila melanogaster* | 158.4 | 5.8 | 7.8 | 8.1 | 57.1 | 26.9 |
| *Arabidopsis thaliana* | 158.4 | 4.2 | 6.5 | 5.0 | 16.2 | 22.1 |

Table 4 presents annotation wall times for all methods. Deep learning models are evaluated on an A100 GPU with 80 GB of memory. **GeneMark** was run on a CPU with 64 threads and 128 GB

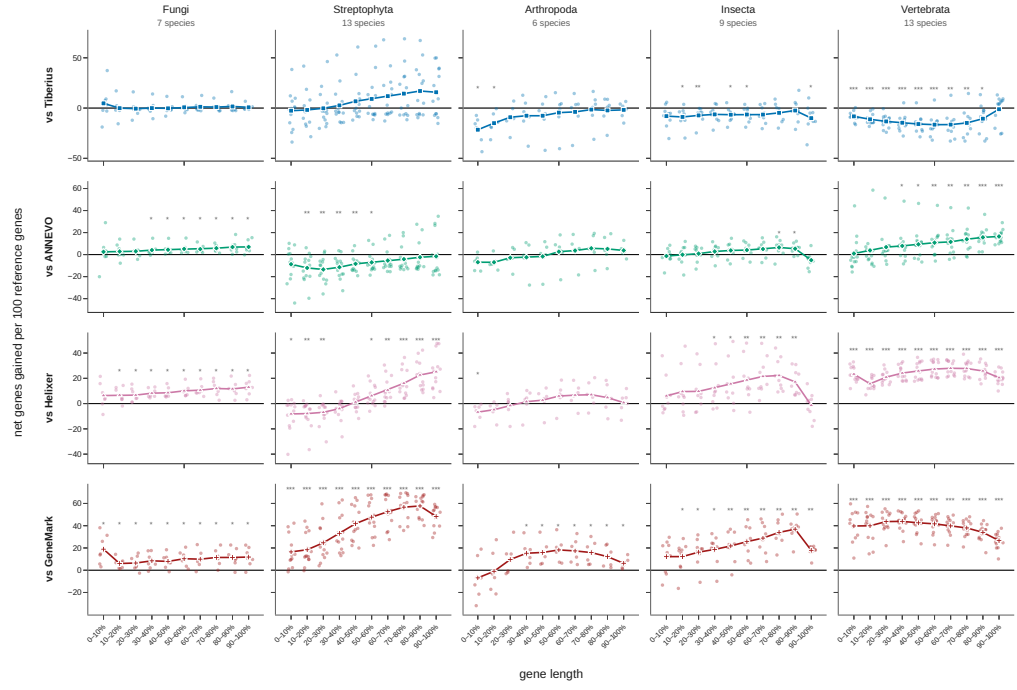

Figure 1: Clade-specific uniquely predicted genes by **Vipsania** against any other tool for genomic span decile.

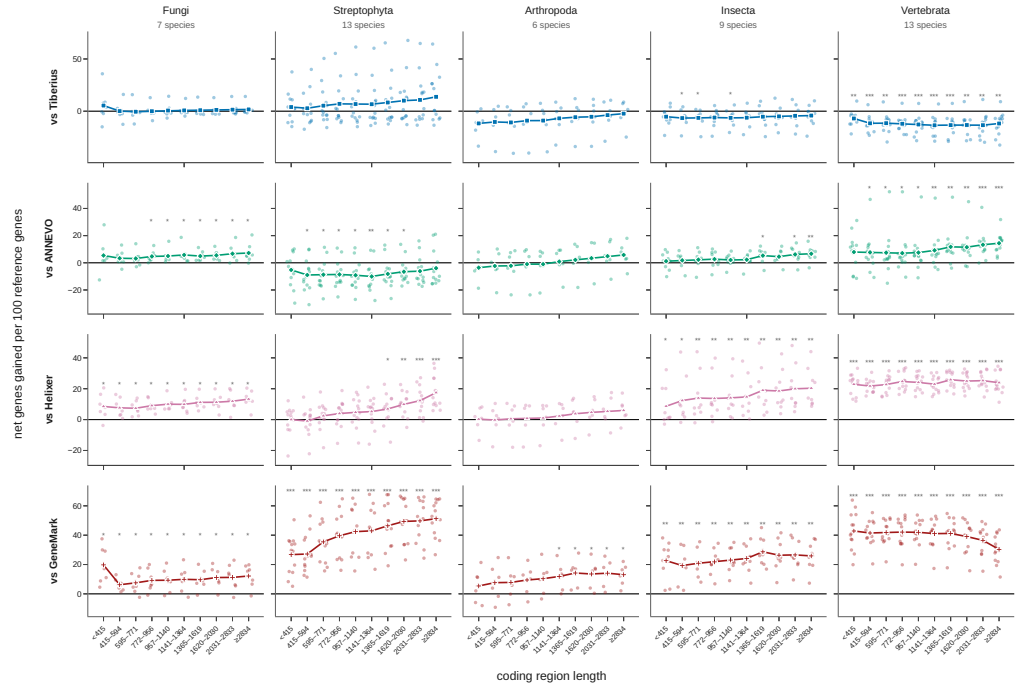

Figure 2: Clade-specific uniquely predicted genes by **Vipsania** against any other tool for length of the coding region.

of RAM. Tables 5 to 21 show sensitivity, precision and F1 metrics for base, exon and locus level comparisons. Vipsania refers to the model with finetuning, Vipsania PT is the pretrained variant without finetuning. Table 22 covers 7 test species in the complement of all eukaryotes and the 17 clade-specific models.

Table 5: Per-species sensitivity / precision for Alveolata. Rows list each tool’s S and P per metric; – denotes a missing measurement. Accession IDs appear below each species name; the final column gives each tool’s BUSCO Complete (C) percentage, and the reference annotation’s BUSCO C is shown under the accession ID.

| Species | Tool | Base |  | Exon |  | Locus |  | BUSCO |
| --- | --- | --- | --- | --- | --- | --- | --- | --- |
|  |  | S | P | S | P | S | P |  |
| Babesia bigemina<br>GCF_000981445.1<br>ref. BUSCO C 99.0 | Vipsania | 0.860 | 0.942 | 0.707 | 0.682 | 0.512 | 0.540 | 100.0 |
|  | Vipsania PT | 0.862 | 0.943 | 0.707 | 0.683 | 0.513 | 0.541 | 100.0 |
|  | GeneMark | 0.839 | 0.952 | 0.589 | 0.577 | 0.345 | 0.434 | 98.0 |
| Besnoitia besnoiti<br>GCF_002563875.1<br>ref. BUSCO C 100.0 | Vipsania | 0.958 | 0.938 | 0.825 | 0.822 | 0.484 | 0.499 | 100.0 |
|  | Vipsania PT | 0.946 | 0.952 | 0.811 | 0.839 | 0.464 | 0.509 | 100.0 |
|  | GeneMark | 0.793 | 0.978 | 0.670 | 0.800 | 0.273 | 0.399 | 93.9 |
| Cryptosporidium meleagridis<br>GCA_039657295.1<br>ref. BUSCO C 100.0 | Vipsania | 0.994 | 0.979 | 0.887 | 0.877 | 0.870 | 0.874 | 100.0 |
|  | Vipsania PT | 0.994 | 0.979 | 0.886 | 0.876 | 0.869 | 0.873 | 100.0 |
|  | GeneMark | 0.986 | 0.978 | 0.777 | 0.756 | 0.751 | 0.802 | 100.0 |
| Plasmodium falciparum<br>GCF_000002765.6<br>ref. BUSCO C 99.0 | Vipsania | 0.990 | 0.981 | 0.920 | 0.912 | 0.870 | 0.844 | 99.0 |
|  | Vipsania PT | 0.955 | 0.980 | 0.655 | 0.812 | 0.668 | 0.681 | 97.0 |
|  | GeneMark | 0.983 | 0.973 | 0.798 | 0.710 | 0.629 | 0.656 | 99.0 |
| Theileria annulata<br>GCF_000003225.4<br>ref. BUSCO C 97.0 | Vipsania | 0.972 | 0.961 | 0.769 | 0.680 | 0.514 | 0.491 | 100.0 |
|  | Vipsania PT | 0.973 | 0.960 | 0.769 | 0.680 | 0.513 | 0.489 | 100.0 |
|  | GeneMark | 0.934 | 0.965 | 0.691 | 0.630 | 0.396 | 0.441 | 96.0 |
| Theileria orientalis strain shintoku<br>GCF_000740895.1<br>ref. BUSCO C 94.9 | Vipsania | 0.974 | 0.930 | 0.683 | 0.695 | 0.430 | 0.434 | 100.0 |
|  | Vipsania PT | 0.974 | 0.930 | 0.683 | 0.694 | 0.429 | 0.433 | 100.0 |
|  | GeneMark | 0.822 | 0.930 | 0.476 | 0.536 | 0.156 | 0.217 | 89.9 |

Table 6: Per-species sensitivity / precision for Amoebozoa. Rows list each tool's S and P per metric; – denotes a missing measurement. Accession IDs appear below each species name; the final column gives each tool's BUSCO Complete (C) percentage, and the reference annotation's BUSCO C is shown under the accession ID.

| Species | Tool | Base |  | Exon |  | Locus |  | BUSCO |
| --- | --- | --- | --- | --- | --- | --- | --- | --- |
|  |  | S | P | S | P | S | P |  |
| Acanthamoeba castellanii str. neff<br>GCF_000313135.1<br>ref. BUSCO C 75.2 | Vipsania | 0.935 | 0.838 | 0.731 | 0.684 | 0.257 | 0.259 | 80.6 |
|  | Vipsania PT | 0.935 | 0.837 | 0.730 | 0.684 | 0.256 | 0.258 | 81.4 |
|  | GeneMark | 0.958 | 0.856 | 0.749 | 0.681 | 0.239 | 0.255 | 80.6 |
| Acytostelium subglobosum lb1<br>GCF_000787575.1<br>ref. BUSCO C 92.2 | Vipsania | 0.983 | 0.869 | 0.669 | 0.612 | 0.377 | 0.356 | 97.7 |
|  | Vipsania PT | 0.983 | 0.869 | 0.669 | 0.612 | 0.378 | 0.356 | 97.7 |
|  | GeneMark | 0.975 | 0.892 | 0.643 | 0.594 | 0.353 | 0.371 | 97.7 |
| Cavenderia fasciculata<br>GCF_000203815.1<br>ref. BUSCO C 89.9 | Vipsania | 0.965 | 0.943 | 0.599 | 0.685 | 0.430 | 0.418 | 97.7 |
|  | Vipsania PT | 0.965 | 0.943 | 0.599 | 0.686 | 0.431 | 0.418 | 97.7 |
|  | GeneMark | 0.966 | 0.950 | 0.597 | 0.705 | 0.409 | 0.424 | 95.3 |
| Dictyostelium discoideum<br>GCF_000004695.1<br>ref. BUSCO C 96.1 | Vipsania | 0.982 | 0.965 | 0.811 | 0.821 | 0.746 | 0.771 | 97.7 |
|  | Vipsania PT | 0.976 | 0.967 | 0.793 | 0.817 | 0.720 | 0.765 | 96.1 |
|  | GeneMark | 0.975 | 0.963 | 0.720 | 0.722 | 0.598 | 0.686 | 96.1 |
| Entamoeba invadens<br>GCF_000330505.1<br>ref. BUSCO C 48.8 | Vipsania | 0.949 | 0.778 | 0.562 | 0.409 | 0.549 | 0.436 | 48.8 |
|  | Vipsania PT | 0.875 | 0.792 | 0.335 | 0.283 | 0.366 | 0.352 | 36.4 |
|  | GeneMark | 0.976 | 0.775 | 0.549 | 0.281 | 0.494 | 0.434 | 51.2 |
| Heterostelium album pn500<br>GCF_000004825.1<br>ref. BUSCO C 89.1 | Vipsania | 0.963 | 0.923 | 0.536 | 0.642 | 0.371 | 0.358 | 97.7 |
|  | Vipsania PT | 0.962 | 0.924 | 0.536 | 0.643 | 0.371 | 0.360 | 97.7 |
|  | GeneMark | 0.948 | 0.928 | 0.494 | 0.603 | 0.292 | 0.324 | 96.1 |
| Pelomyxa schiedti<br>GCA_020536535.1<br>ref. BUSCO C 86.0 | Vipsania | 0.848 | 0.893 | 0.739 | 0.841 | 0.374 | 0.438 | 83.7 |
|  | Vipsania PT | 0.849 | 0.894 | 0.740 | 0.838 | 0.372 | 0.434 | 83.7 |
|  | GeneMark | 0.535 | 0.826 | 0.190 | 0.461 | 0.045 | 0.056 | 14.7 |

Table 7: Per-species sensitivity / precision for Arthropoda. Rows list each tool's S and P per metric; – denotes a missing measurement. Accession IDs appear below each species name; the final column gives each tool's BUSCO Complete (C) percentage, and the reference annotation's BUSCO C is shown under the accession ID.

| Species | Tool | Base |  | Exon |  | Locus |  | BUSCO |
| --- | --- | --- | --- | --- | --- | --- | --- | --- |
|  |  | S | P | S | P | S | P |  |
| Artemia franciscana<br>GCF_032884065.1<br>ref. BUSCO C 90.9 | Vipsania | 0.686 | 0.909 | 0.625 | 0.870 | 0.199 | 0.359 | 66.6 |
|  | Vipsania PT | 0.663 | 0.896 | 0.626 | 0.824 | 0.186 | 0.298 | 66.8 |
|  | Tiberius | 0.707 | 0.543 | 0.633 | 0.599 | 0.196 | 0.090 | 58.5 |
|  | ANNEVO | 0.750 | 0.829 | 0.637 | 0.548 | 0.117 | 0.108 | 63.8 |
|  | Helixer | 0.718 | 0.830 | 0.626 | 0.546 | 0.099 | 0.114 | 60.0 |
|  | GeneMark | 0.707 | 0.217 | 0.308 | 0.120 | 0.084 | 0.021 | 25.6 |
| Cherax quadricarinatus<br>GCF_038502225.1<br>ref. BUSCO C 87.9 | Vipsania | 0.716 | 0.902 | 0.627 | 0.845 | 0.247 | 0.320 | 71.8 |
|  | Vipsania PT | 0.690 | 0.903 | 0.597 | 0.841 | 0.227 | 0.300 | 70.8 |
|  | Tiberius | 0.664 | 0.325 | 0.600 | 0.481 | 0.181 | 0.041 | 60.2 |
|  | ANNEVO | 0.793 | 0.830 | 0.659 | 0.590 | 0.143 | 0.107 | 69.0 |
|  | Helixer | 0.771 | 0.687 | 0.627 | 0.560 | 0.144 | 0.107 | 62.5 |
|  | GeneMark | 0.025 | 0.004 | 0.000 | 0.000 | 0.000 | 0.000 | 0.1 |
| Dermacentor silvarum<br>GCF_013339745.2<br>ref. BUSCO C 93.6 | Vipsania | 0.675 | 0.742 | 0.650 | 0.706 | 0.194 | 0.264 | 75.7 |
|  | Vipsania PT | 0.622 | 0.757 | 0.583 | 0.716 | 0.158 | 0.238 | 66.6 |
|  | Tiberius | 0.753 | 0.562 | 0.693 | 0.577 | 0.314 | 0.138 | 83.2 |
|  | ANNEVO | 0.777 | 0.761 | 0.691 | 0.554 | 0.201 | 0.170 | 86.3 |
|  | Helixer | 0.660 | 0.817 | 0.618 | 0.630 | 0.164 | 0.214 | 73.3 |
|  | GeneMark | 0.653 | 0.103 | 0.175 | 0.036 | 0.050 | 0.007 | 23.6 |
| Parasteatoda tepidariorum<br>GCF_043381705.1<br>ref. BUSCO C 95.5 | Vipsania | 0.561 | 0.908 | 0.445 | 0.885 | 0.117 | 0.257 | 45.1 |
|  | Vipsania PT | 0.803 | 0.896 | 0.767 | 0.861 | 0.294 | 0.376 | 85.5 |
|  | Tiberius | 0.872 | 0.733 | 0.798 | 0.755 | 0.460 | 0.232 | 90.8 |
|  | ANNEVO | 0.860 | 0.913 | 0.805 | 0.748 | 0.298 | 0.290 | 91.9 |
|  | Helixer | 0.876 | 0.864 | 0.805 | 0.677 | 0.249 | 0.219 | 83.0 |
|  | GeneMark | 0.852 | 0.442 | 0.561 | 0.319 | 0.130 | 0.043 | 53.6 |
| Penaeus monodon<br>GCF_015228065.2<br>ref. BUSCO C 76.5 | Vipsania | 0.649 | 0.855 | 0.548 | 0.759 | 0.167 | 0.288 | 61.5 |
|  | Vipsania PT | 0.710 | 0.830 | 0.590 | 0.733 | 0.182 | 0.275 | 67.8 |
|  | Tiberius | 0.755 | 0.347 | 0.581 | 0.362 | 0.274 | 0.047 | 61.0 |
|  | ANNEVO | 0.760 | 0.829 | 0.605 | 0.583 | 0.185 | 0.192 | 69.7 |
|  | Helixer | 0.810 | 0.786 | 0.642 | 0.550 | 0.186 | 0.171 | 71.1 |
|  | GeneMark | 0.569 | 0.180 | 0.189 | 0.034 | 0.023 | 0.005 | 10.3 |

continued on next page

Table 7: Arthropoda (continued)

| Species | Tool | Base |  | Exon |  | Locus |  | BUSCO |
| --- | --- | --- | --- | --- | --- | --- | --- | --- |
|  |  | S | P | S | P | S | P |  |
| Pycnogonum litorale<br>GCA_964442445.1<br>ref. BUSCO C 41.0 | Vipsania | 0.601 | 0.847 | 0.544 | 0.715 | 0.098 | 0.127 | 89.6 |
|  | Vipsania PT | 0.572 | 0.834 | 0.507 | 0.705 | 0.065 | 0.095 | 80.9 |
|  | Tiberius | 0.541 | 0.681 | 0.452 | 0.609 | 0.084 | 0.059 | 67.1 |
|  | ANNEVO | 0.567 | 0.795 | 0.466 | 0.551 | 0.041 | 0.047 | 68.6 |
|  | Helixer | 0.553 | 0.675 | 0.445 | 0.429 | 0.040 | 0.033 | 52.2 |
|  | GeneMark | 0.650 | 0.549 | 0.504 | 0.525 | 0.085 | 0.052 | 78.8 |

Table 8: Per-species sensitivity / precision for Chlorophyta. Rows list each tool's S and P per metric; – denotes a missing measurement. Accession IDs appear below each species name; the final column gives each tool's BUSCO Complete (C) percentage, and the reference annotation's BUSCO C is shown under the accession ID.

| Species | Tool | Base |  | Exon |  | Locus |  | BUSCO |
| --- | --- | --- | --- | --- | --- | --- | --- | --- |
|  |  | S | P | S | P | S | P |  |
| Chlamydomonas reinhardtii<br>GCF_000002595.2<br>ref. BUSCO C 98.6 | Vipsania | 0.976 | 0.912 | 0.916 | 0.921 | 0.606 | 0.620 | 98.6 |
|  | Vipsania PT | 0.959 | 0.904 | 0.879 | 0.892 | 0.493 | 0.491 | 97.8 |
|  | Tiberius | 0.982 | 0.986 | 0.937 | 0.958 | 0.747 | 0.778 | 98.0 |
|  | GeneMark | 0.927 | 0.954 | 0.810 | 0.834 | 0.263 | 0.288 | 93.2 |
| Micromonas commoda<br>GCF_000090985.2<br>ref. BUSCO C 93.9 | Vipsania | 0.985 | 0.842 | 0.591 | 0.587 | 0.561 | 0.543 | 98.6 |
|  | Vipsania PT | 0.985 | 0.842 | 0.589 | 0.586 | 0.560 | 0.542 | 98.6 |
|  | Tiberius | 0.944 | 0.849 | 0.490 | 0.538 | 0.468 | 0.486 | 94.4 |
|  | GeneMark | 0.873 | 0.853 | 0.448 | 0.485 | 0.431 | 0.486 | 86.0 |
| Ostreococcus tauri<br>GCF_000214015.3<br>ref. BUSCO C 95.7 | Vipsania | 0.991 | 0.969 | 0.753 | 0.734 | 0.725 | 0.715 | 95.9 |
|  | Vipsania PT | 0.991 | 0.969 | 0.750 | 0.732 | 0.724 | 0.716 | 95.9 |
|  | Tiberius | 0.907 | 0.983 | 0.511 | 0.576 | 0.504 | 0.551 | 89.2 |
|  | GeneMark | 0.890 | 0.974 | 0.288 | 0.239 | 0.229 | 0.286 | 87.3 |
| Pseudoscurfieldia marina<br>GCA_049488355.1<br>ref. BUSCO C 92.1 | Vipsania | 0.950 | 0.961 | 0.644 | 0.679 | 0.668 | 0.710 | 92.9 |
|  | Vipsania PT | 0.943 | 0.961 | 0.640 | 0.679 | 0.663 | 0.710 | 92.9 |
|  | Tiberius | 0.834 | 0.979 | 0.416 | 0.466 | 0.426 | 0.510 | 84.1 |
|  | GeneMark | 0.917 | 0.971 | 0.544 | 0.443 | 0.558 | 0.625 | 91.6 |
| Pycnococcus provasolii<br>GCA_049487715.1<br>ref. BUSCO C 90.2 | Vipsania | 0.981 | 0.956 | 0.609 | 0.601 | 0.624 | 0.646 | 92.8 |
|  | Vipsania PT | 0.980 | 0.956 | 0.607 | 0.600 | 0.623 | 0.645 | 92.8 |
|  | Tiberius | 0.862 | 0.976 | 0.407 | 0.447 | 0.415 | 0.485 | 83.5 |
|  | GeneMark | 0.957 | 0.971 | 0.489 | 0.383 | 0.481 | 0.523 | 91.8 |
| Tetrademus obliquus<br>GCA_030272155.1<br>ref. BUSCO C 96.8 | Vipsania | 0.977 | 0.596 | 0.764 | 0.754 | 0.302 | 0.240 | 99.7 |
|  | Vipsania PT | 0.975 | 0.599 | 0.762 | 0.751 | 0.293 | 0.239 | 99.7 |
|  | Tiberius | 0.937 | 0.945 | 0.825 | 0.870 | 0.487 | 0.472 | 96.6 |
|  | GeneMark | 0.923 | 0.644 | 0.686 | 0.650 | 0.139 | 0.130 | 91.5 |

Table 9: Per-species sensitivity / precision for Cnidaria. Rows list each tool's S and P per metric; – denotes a missing measurement. Accession IDs appear below each species name; the final column gives each tool's BUSCO Complete (C) percentage, and the reference annotation's BUSCO C is shown under the accession ID.

| Species | Tool | Base |  | Exon |  | Locus |  | BUSCO |
| --- | --- | --- | --- | --- | --- | --- | --- | --- |
|  |  | S | P | S | P | S | P |  |
| Acropora digitifera<br>GCF_000222465.1<br>ref. BUSCO C 63.2 | Vipsania | 0.737 | 0.779 | 0.639 | 0.695 | 0.197 | 0.273 | 54.6 |
|  | Vipsania PT | 0.722 | 0.783 | 0.619 | 0.705 | 0.187 | 0.267 | 51.3 |
|  | ANNEVO | 0.753 | 0.788 | 0.594 | 0.531 | 0.142 | 0.170 | 46.7 |
|  | Helixer | 0.694 | 0.658 | 0.525 | 0.373 | 0.108 | 0.086 | 31.4 |
|  | GeneMark | 0.882 | 0.526 | 0.620 | 0.452 | 0.176 | 0.105 | 43.2 |
| Hydra vulgaris<br>GCF_038396675.1<br>ref. BUSCO C 93.6 | Vipsania | 0.636 | 0.920 | 0.703 | 0.878 | 0.257 | 0.484 | 78.0 |
|  | Vipsania PT | 0.442 | 0.928 | 0.473 | 0.879 | 0.118 | 0.317 | 43.6 |
|  | ANNEVO | 0.595 | 0.831 | 0.583 | 0.639 | 0.107 | 0.152 | 60.3 |
|  | Helixer | 0.473 | 0.775 | 0.415 | 0.591 | 0.086 | 0.131 | 32.9 |
|  | GeneMark | 0.716 | 0.157 | 0.168 | 0.043 | 0.080 | 0.015 | 23.5 |
| Hydractinia symbiolongicarpus<br>GCF_029227915.1<br>ref. BUSCO C 94.0 | Vipsania | 0.816 | 0.894 | 0.834 | 0.884 | 0.493 | 0.561 | 91.4 |
|  | Vipsania PT | 0.787 | 0.896 | 0.789 | 0.882 | 0.388 | 0.485 | 83.0 |
|  | ANNEVO | 0.799 | 0.859 | 0.717 | 0.707 | 0.238 | 0.254 | 79.8 |
|  | Helixer | 0.883 | 0.697 | 0.754 | 0.501 | 0.221 | 0.134 | 76.5 |
|  | GeneMark | 0.926 | 0.500 | 0.766 | 0.507 | 0.280 | 0.124 | 78.3 |

continued on next page

Table 9: Cnidaria (continued)

| Species | Tool | Base |  | Exon |  | Locus |  | BUSCO |
| --- | --- | --- | --- | --- | --- | --- | --- | --- |
|  |  | S | P | S | P | S | P |  |
| Montipora capricornis<br>GCF_036669925.1<br>ref. BUSCO C 95.7 | Vipsania | 0.656 | 0.882 | 0.745 | 0.856 | 0.291 | 0.486 | 87.1 |
|  | Vipsania PT | 0.696 | 0.858 | 0.759 | 0.834 | 0.328 | 0.455 | 89.1 |
|  | ANNEVO | 0.739 | 0.770 | 0.683 | 0.599 | 0.206 | 0.205 | 74.0 |
|  | Helixer | 0.662 | 0.706 | 0.598 | 0.484 | 0.128 | 0.118 | 41.4 |
|  | GeneMark | 0.806 | 0.316 | 0.403 | 0.242 | 0.163 | 0.062 | 28.3 |
| Montipora foliosa<br>GCF_036669935.1<br>ref. BUSCO C 95.8 | Vipsania | 0.679 | 0.891 | 0.771 | 0.867 | 0.330 | 0.516 | 91.2 |
|  | Vipsania PT | 0.703 | 0.873 | 0.770 | 0.855 | 0.347 | 0.479 | 90.6 |
|  | ANNEVO | 0.745 | 0.784 | 0.691 | 0.616 | 0.216 | 0.217 | 76.0 |
|  | Helixer | 0.669 | 0.720 | 0.602 | 0.497 | 0.132 | 0.123 | 43.2 |
|  | GeneMark | 0.814 | 0.326 | 0.424 | 0.259 | 0.169 | 0.064 | 29.0 |
| Nematostella vectensis<br>GCF_932526225.1<br>ref. BUSCO C 96.0 | Vipsania | 0.890 | 0.940 | 0.876 | 0.924 | 0.593 | 0.626 | 94.5 |
|  | Vipsania PT | 0.860 | 0.942 | 0.827 | 0.929 | 0.475 | 0.547 | 88.5 |
|  | ANNEVO | 0.909 | 0.929 | 0.821 | 0.790 | 0.382 | 0.380 | 90.8 |
|  | Helixer | 0.888 | 0.853 | 0.788 | 0.706 | 0.286 | 0.245 | 75.1 |
|  | GeneMark | 0.911 | 0.602 | 0.723 | 0.605 | 0.253 | 0.141 | 65.0 |
| Oculina patagonica<br>GCF_052425735.1<br>ref. BUSCO C 95.8 | Vipsania | 0.827 | 0.858 | 0.840 | 0.851 | 0.494 | 0.542 | 93.9 |
|  | Vipsania PT | 0.828 | 0.860 | 0.829 | 0.856 | 0.483 | 0.533 | 92.4 |
|  | ANNEVO | 0.883 | 0.825 | 0.798 | 0.696 | 0.355 | 0.322 | 84.7 |
|  | Helixer | 0.901 | 0.715 | 0.780 | 0.534 | 0.269 | 0.172 | 69.2 |
|  | GeneMark | 0.938 | 0.632 | 0.788 | 0.648 | 0.358 | 0.223 | 79.3 |
| Pocillopora verrucosa<br>GCF_036669915.1<br>ref. BUSCO C 96.4 | Vipsania | 0.897 | 0.869 | 0.867 | 0.842 | 0.542 | 0.542 | 93.3 |
|  | Vipsania PT | 0.866 | 0.883 | 0.837 | 0.860 | 0.476 | 0.524 | 91.8 |
|  | ANNEVO | 0.921 | 0.860 | 0.806 | 0.707 | 0.376 | 0.337 | 89.9 |
|  | Helixer | 0.921 | 0.760 | 0.789 | 0.544 | 0.258 | 0.176 | 78.1 |
|  | GeneMark | 0.922 | 0.697 | 0.796 | 0.669 | 0.356 | 0.238 | 83.3 |
| Porites lutea<br>GCF_958299795.1<br>ref. BUSCO C 95.8 | Vipsania | 0.807 | 0.826 | 0.845 | 0.817 | 0.432 | 0.483 | 93.5 |
|  | Vipsania PT | 0.817 | 0.821 | 0.837 | 0.814 | 0.427 | 0.463 | 91.2 |
|  | ANNEVO | 0.880 | 0.770 | 0.792 | 0.627 | 0.302 | 0.253 | 86.9 |
|  | Helixer | 0.860 | 0.688 | 0.753 | 0.487 | 0.209 | 0.139 | 69.8 |
|  | GeneMark | 0.932 | 0.551 | 0.775 | 0.572 | 0.312 | 0.161 | 77.1 |
| Rhopilema esculentum<br>GCF_013076305.1<br>ref. BUSCO C 92.7 | Vipsania | 0.866 | 0.928 | 0.882 | 0.916 | 0.536 | 0.569 | 91.1 |
|  | Vipsania PT | 0.853 | 0.928 | 0.858 | 0.913 | 0.462 | 0.522 | 87.5 |
|  | ANNEVO | 0.846 | 0.912 | 0.782 | 0.751 | 0.281 | 0.295 | 80.4 |
|  | Helixer | 0.865 | 0.814 | 0.773 | 0.591 | 0.191 | 0.153 | 66.1 |
|  | GeneMark | 0.937 | 0.701 | 0.803 | 0.714 | 0.315 | 0.197 | 75.3 |

Table 10: Per-species sensitivity / precision for Discoba. Rows list each tool's S and P per metric; – denotes a missing measurement. Accession IDs appear below each species name; the final column gives each tool's BUSCO Complete (C) percentage, and the reference annotation's BUSCO C is shown under the accession ID.

| Species | Tool | Base |  | Exon |  | Locus |  | BUSCO |
| --- | --- | --- | --- | --- | --- | --- | --- | --- |
|  |  | S | P | S | P | S | P |  |
| Angomonas deanei<br>GCA_903995115.1<br>ref. BUSCO C 41.9 | Vipsania | 0.965 | 0.773 | 0.470 | 0.379 | 0.470 | 0.506 | 51.2 |
|  | Vipsania PT | 0.964 | 0.771 | 0.469 | 0.377 | 0.469 | 0.504 | 51.2 |
|  | GeneMark | 0.951 | 0.800 | 0.341 | 0.176 | 0.341 | 0.451 | 51.2 |
| Leishmania major<br>GCF_000002725.2<br>ref. BUSCO C 55.0 | Vipsania | 0.970 | 0.958 | 0.742 | 0.726 | 0.742 | 0.741 | 55.0 |
|  | Vipsania PT | 0.970 | 0.956 | 0.739 | 0.712 | 0.739 | 0.737 | 55.0 |
|  | GeneMark | 0.972 | 0.935 | 0.581 | 0.402 | 0.581 | 0.617 | 55.8 |
| Porcisia hertigi<br>GCF_017918235.1<br>ref. BUSCO C 55.8 | Vipsania | 0.973 | 0.839 | 0.791 | 0.682 | 0.828 | 0.748 | 54.3 |
|  | Vipsania PT | 0.972 | 0.841 | 0.789 | 0.683 | 0.826 | 0.749 | 54.3 |
|  | GeneMark | 0.981 | 0.830 | 0.602 | 0.311 | 0.629 | 0.612 | 55.0 |
| Trypanosoma brucei brucei treu927<br>GCF_000002445.2<br>ref. BUSCO C 55.0 | Vipsania | 0.937 | 0.918 | 0.753 | 0.664 | 0.753 | 0.789 | 52.7 |
|  | Vipsania PT | 0.937 | 0.919 | 0.754 | 0.671 | 0.754 | 0.792 | 53.5 |
|  | GeneMark | 0.953 | 0.877 | 0.664 | 0.386 | 0.664 | 0.720 | 54.3 |

Table 11: Per-species sensitivity / precision for Echinodermata. Rows list each tool's S and P per metric; – denotes a missing measurement. Accession IDs appear below each species name; the final column gives each tool's BUSCO Complete (C) percentage, and the reference annotation's BUSCO C is shown under the accession ID.

| Species | Tool | Base |  | Exon |  | Locus |  | BUSCO C% |
| --- | --- | --- | --- | --- | --- | --- | --- | --- |
|  |  | S | P | S | P | S | P |  |
| Amphiura filiformis<br>GCF_039555335.1<br>ref. BUSCO C 98.5 | Vipsania | 0.709 | 0.678 | 0.735 | 0.744 | 0.242 | 0.288 | 82.1 |
|  | Vipsania PT | 0.810 | 0.672 | 0.801 | 0.724 | 0.323 | 0.313 | 91.8 |
|  | ANNEVO | 0.787 | 0.756 | 0.739 | 0.620 | 0.193 | 0.180 | 88.4 |
|  | Helixer | 0.848 | 0.645 | 0.769 | 0.503 | 0.166 | 0.114 | 81.1 |
|  | GeneMark | 0.927 | 0.403 | 0.726 | 0.415 | 0.181 | 0.070 | 74.4 |
| Antedon bifida<br>GCF_963402885.1<br>ref. BUSCO C 99.3 | Vipsania | 0.933 | 0.820 | 0.891 | 0.846 | 0.553 | 0.519 | 97.0 |
|  | Vipsania PT | 0.930 | 0.816 | 0.885 | 0.837 | 0.523 | 0.497 | 97.6 |
|  | ANNEVO | 0.916 | 0.852 | 0.838 | 0.769 | 0.354 | 0.343 | 94.0 |
|  | Helixer | 0.958 | 0.742 | 0.848 | 0.644 | 0.267 | 0.184 | 88.2 |
|  | GeneMark | 0.953 | 0.647 | 0.812 | 0.679 | 0.295 | 0.184 | 83.9 |
| Antedon mediterranea<br>GCF_964355755.1<br>ref. BUSCO C 99.1 | Vipsania | 0.894 | 0.832 | 0.880 | 0.842 | 0.502 | 0.509 | 95.4 |
|  | Vipsania PT | 0.919 | 0.809 | 0.884 | 0.827 | 0.508 | 0.484 | 97.6 |
|  | ANNEVO | 0.902 | 0.849 | 0.833 | 0.755 | 0.332 | 0.324 | 92.9 |
|  | Helixer | 0.942 | 0.723 | 0.839 | 0.626 | 0.244 | 0.164 | 84.5 |
|  | GeneMark | 0.954 | 0.607 | 0.817 | 0.652 | 0.299 | 0.171 | 86.3 |
| Apostichopus japonicus<br>GCF_037975245.1<br>ref. BUSCO C 99.1 | Vipsania | 0.891 | 0.907 | 0.848 | 0.899 | 0.521 | 0.566 | 96.0 |
|  | Vipsania PT | 0.889 | 0.905 | 0.841 | 0.892 | 0.482 | 0.537 | 95.2 |
|  | ANNEVO | 0.874 | 0.911 | 0.782 | 0.763 | 0.313 | 0.297 | 94.0 |
|  | Helixer | 0.923 | 0.808 | 0.797 | 0.631 | 0.224 | 0.169 | 87.9 |
|  | GeneMark | 0.931 | 0.678 | 0.779 | 0.678 | 0.267 | 0.163 | 89.6 |
| Asterias rubens<br>GCF_902459465.1<br>ref. BUSCO C 98.2 | Vipsania | 0.925 | 0.905 | 0.889 | 0.908 | 0.565 | 0.578 | 96.3 |
|  | Vipsania PT | 0.920 | 0.900 | 0.885 | 0.902 | 0.535 | 0.561 | 96.0 |
|  | ANNEVO | 0.930 | 0.905 | 0.850 | 0.781 | 0.346 | 0.333 | 93.3 |
|  | Helixer | 0.939 | 0.821 | 0.838 | 0.655 | 0.238 | 0.186 | 86.6 |
|  | GeneMark | 0.950 | 0.733 | 0.822 | 0.729 | 0.320 | 0.208 | 84.7 |
| Holothuria leucospilota<br>GCA_029531755.1<br>ref. BUSCO C 89.3 | Vipsania | 0.697 | 0.829 | 0.725 | 0.749 | 0.183 | 0.317 | 89.9 |
|  | Vipsania PT | 0.719 | 0.827 | 0.735 | 0.745 | 0.191 | 0.314 | 91.5 |
|  | ANNEVO | 0.653 | 0.836 | 0.640 | 0.613 | 0.090 | 0.139 | 86.3 |
|  | Helixer | 0.699 | 0.779 | 0.648 | 0.534 | 0.076 | 0.105 | 75.0 |
|  | GeneMark | 0.839 | 0.555 | 0.663 | 0.521 | 0.135 | 0.097 | 77.5 |
| Lytechinus pictus<br>GCF_037042905.1<br>ref. BUSCO C 93.5 | Vipsania | 0.875 | 0.900 | 0.864 | 0.877 | 0.442 | 0.511 | 96.6 |
|  | Vipsania PT | 0.894 | 0.891 | 0.874 | 0.864 | 0.452 | 0.502 | 97.2 |
|  | ANNEVO | 0.871 | 0.879 | 0.809 | 0.717 | 0.298 | 0.285 | 94.6 |
|  | Helixer | 0.914 | 0.790 | 0.811 | 0.628 | 0.214 | 0.166 | 83.3 |
|  | GeneMark | 0.950 | 0.599 | 0.800 | 0.582 | 0.241 | 0.122 | 82.1 |
| Strongylocentrotus purpuratus<br>GCF_000002235.5<br>ref. BUSCO C 99.0 | Vipsania | 0.864 | 0.773 | 0.841 | 0.770 | 0.417 | 0.436 | 97.3 |
|  | Vipsania PT | 0.878 | 0.767 | 0.847 | 0.765 | 0.420 | 0.431 | 97.8 |
|  | ANNEVO | 0.888 | 0.840 | 0.813 | 0.739 | 0.334 | 0.327 | 97.3 |
|  | Helixer | 0.933 | 0.683 | 0.812 | 0.567 | 0.229 | 0.161 | 89.9 |
|  | GeneMark | 0.952 | 0.562 | 0.779 | 0.533 | 0.207 | 0.105 | 86.9 |

Table 12: Per-species sensitivity / precision for Fungi. Rows list each tool's S and P per metric; – denotes a missing measurement. Accession IDs appear below each species name; the final column gives each tool's BUSCO Complete (C) percentage, and the reference annotation's BUSCO C is shown under the accession ID.

| Species | Tool | Base |  | Exon |  | Locus |  | BUSCO C% |
| --- | --- | --- | --- | --- | --- | --- | --- | --- |
|  |  | S | P | S | P | S | P |  |
| Aspergillus fumigatus<br>GCF_000002655.1<br>ref. BUSCO C 98.5 | Vipsania | 0.964 | 0.956 | 0.816 | 0.774 | 0.655 | 0.650 | 99.1 |
|  | Vipsania PT | 0.965 | 0.956 | 0.813 | 0.773 | 0.650 | 0.646 | 99.1 |
|  | Tiberius | 0.956 | 0.965 | 0.803 | 0.796 | 0.644 | 0.657 | 99.0 |
|  | ANNEVO | 0.965 | 0.960 | 0.792 | 0.750 | 0.623 | 0.616 | 99.1 |
|  | Helixer | 0.974 | 0.941 | 0.750 | 0.671 | 0.567 | 0.532 | 99.3 |
|  | GeneMark | 0.938 | 0.963 | 0.693 | 0.717 | 0.506 | 0.526 | 96.7 |
| Coccidioides immitis<br>GCF_000149335.2<br>ref. BUSCO C 99.2 | Vipsania | 0.908 | 0.981 | 0.741 | 0.908 | 0.634 | 0.800 | 99.4 |
|  | Vipsania PT | 0.910 | 0.975 | 0.738 | 0.898 | 0.627 | 0.785 | 99.3 |
|  | Tiberius | 0.921 | 0.970 | 0.748 | 0.894 | 0.650 | 0.771 | 99.3 |
|  | ANNEVO | 0.926 | 0.976 | 0.722 | 0.836 | 0.595 | 0.704 | 99.0 |
|  | Helixer | 0.931 | 0.947 | 0.703 | 0.773 | 0.561 | 0.620 | 98.8 |
|  | GeneMark | 0.911 | 0.960 | 0.619 | 0.772 | 0.463 | 0.571 | 97.4 |

continued on next page

Table 12: Fungi (continued)

| Species | Tool | Base |  | Exon |  | Locus |  | BUSCO |
| --- | --- | --- | --- | --- | --- | --- | --- | --- |
|  |  | S | P | S | P | S | P |  |
| Hanseniaspora uvarum<br>GCA_050947715.1<br>ref. BUSCO C 50.7 | Vipsania | 0.995 | 0.955 | 0.815 | 0.791 | 0.864 | 0.808 | 51.2 |
|  | Vipsania PT | 0.996 | 0.955 | 0.812 | 0.793 | 0.862 | 0.807 | 51.3 |
|  | Tiberius | 0.993 | 0.955 | 0.796 | 0.776 | 0.848 | 0.792 | 51.0 |
|  | ANNEVO | 0.990 | 0.956 | 0.801 | 0.773 | 0.854 | 0.789 | 50.6 |
|  | Helixer | 0.994 | 0.949 | 0.768 | 0.702 | 0.822 | 0.748 | 50.6 |
|  | GeneMark | 0.993 | 0.962 | 0.830 | 0.795 | 0.854 | 0.841 | 50.5 |
| Psilocybe cubensis<br>GCF_017499595.1<br>ref. BUSCO C 94.7 | Vipsania | 0.965 | 0.822 | 0.826 | 0.711 | 0.458 | 0.400 | 99.8 |
|  | Vipsania PT | 0.965 | 0.825 | 0.824 | 0.712 | 0.454 | 0.400 | 99.8 |
|  | Tiberius | 0.963 | 0.888 | 0.830 | 0.754 | 0.483 | 0.448 | 99.3 |
|  | ANNEVO | 0.968 | 0.903 | 0.784 | 0.695 | 0.395 | 0.362 | 98.8 |
|  | Helixer | 0.975 | 0.792 | 0.732 | 0.563 | 0.288 | 0.220 | 98.5 |
|  | GeneMark | 0.947 | 0.847 | 0.768 | 0.710 | 0.392 | 0.369 | 96.7 |
| Rhizophagus irregularis<br>GCF_026210795.1<br>ref. BUSCO C 95.5 | Vipsania | 0.871 | 0.709 | 0.652 | 0.683 | 0.453 | 0.483 | 97.0 |
|  | Vipsania PT | 0.890 | 0.693 | 0.649 | 0.640 | 0.431 | 0.449 | 96.6 |
|  | Tiberius | 0.913 | 0.815 | 0.717 | 0.761 | 0.542 | 0.550 | 96.7 |
|  | ANNEVO | 0.935 | 0.902 | 0.679 | 0.681 | 0.465 | 0.466 | 96.9 |
|  | Helixer | 0.953 | 0.664 | 0.642 | 0.493 | 0.398 | 0.295 | 95.8 |
|  | GeneMark | 0.953 | 0.631 | 0.679 | 0.503 | 0.381 | 0.330 | 95.5 |
| Saitoella coloradoensis<br>GCA_051599425.1<br>ref. BUSCO C 97.7 | Vipsania | 0.990 | 0.948 | 0.871 | 0.812 | 0.787 | 0.748 | 99.2 |
|  | Vipsania PT | 0.993 | 0.947 | 0.869 | 0.811 | 0.789 | 0.747 | 99.1 |
|  | Tiberius | 0.990 | 0.950 | 0.863 | 0.825 | 0.784 | 0.756 | 98.9 |
|  | ANNEVO | 0.989 | 0.950 | 0.845 | 0.783 | 0.754 | 0.714 | 99.0 |
|  | Helixer | 0.987 | 0.932 | 0.777 | 0.688 | 0.652 | 0.605 | 98.3 |
|  | GeneMark | 0.970 | 0.970 | 0.776 | 0.798 | 0.718 | 0.727 | 95.5 |
| Vairimorpha necatrix<br>GCF_036630325.1<br>ref. BUSCO C 11.6 | Vipsania | 0.934 | 0.710 | 0.676 | 0.682 | 0.811 | 0.719 | 11.7 |
|  | Vipsania PT | 0.936 | 0.666 | 0.638 | 0.593 | 0.762 | 0.649 | 11.5 |
|  | Tiberius | 0.902 | 0.639 | 0.532 | 0.395 | 0.635 | 0.470 | 11.1 |
|  | ANNEVO | 0.883 | 0.717 | 0.537 | 0.410 | 0.644 | 0.481 | 10.6 |
|  | Helixer | 0.973 | 0.546 | 0.589 | 0.260 | 0.698 | 0.385 | 11.5 |
|  | GeneMark | 0.920 | 0.488 | 0.550 | 0.137 | 0.631 | 0.441 | 10.5 |

Table 13: Per-species sensitivity / precision for Insecta. Rows list each tool's S and P per metric; – denotes a missing measurement. Accession IDs appear below each species name; the final column gives each tool's BUSCO Complete (C) percentage, and the reference annotation's BUSCO C is shown under the accession ID.

| Species | Tool | Base |  | Exon |  | Locus |  | BUSCO |
| --- | --- | --- | --- | --- | --- | --- | --- | --- |
|  |  | S | P | S | P | S | P |  |
| Aphis gossypii<br>GCF_020184175.1<br>ref. BUSCO C 93.7 | Vipsania | 0.772 | 0.944 | 0.770 | 0.901 | 0.455 | 0.599 | 89.9 |
|  | Vipsania PT | 0.787 | 0.907 | 0.782 | 0.854 | 0.433 | 0.545 | 88.3 |
|  | Tiberius | 0.916 | 0.727 | 0.846 | 0.792 | 0.679 | 0.422 | 92.5 |
|  | ANNEVO | 0.917 | 0.837 | 0.821 | 0.792 | 0.529 | 0.440 | 92.1 |
|  | Helixer | 0.854 | 0.868 | 0.800 | 0.779 | 0.435 | 0.411 | 90.0 |
|  | GeneMark | 0.885 | 0.522 | 0.570 | 0.391 | 0.148 | 0.081 | 73.4 |
| Apis mellifera<br>GCF_003254395.2<br>ref. BUSCO C 97.8 | Vipsania | 0.953 | 0.968 | 0.814 | 0.920 | 0.640 | 0.650 | 96.8 |
|  | Vipsania PT | 0.948 | 0.969 | 0.802 | 0.916 | 0.574 | 0.621 | 95.1 |
|  | Tiberius | 0.960 | 0.903 | 0.843 | 0.879 | 0.756 | 0.522 | 98.1 |
|  | ANNEVO | 0.966 | 0.966 | 0.821 | 0.867 | 0.610 | 0.577 | 98.2 |
|  | Helixer | 0.962 | 0.953 | 0.808 | 0.831 | 0.574 | 0.509 | 95.8 |
|  | GeneMark | 0.949 | 0.761 | 0.749 | 0.596 | 0.396 | 0.202 | 93.2 |
| Bombyx mori<br>GCF_030269925.1<br>ref. BUSCO C 98.4 | Vipsania | 0.861 | 0.947 | 0.825 | 0.918 | 0.553 | 0.600 | 95.9 |
|  | Vipsania PT | 0.821 | 0.952 | 0.789 | 0.916 | 0.448 | 0.553 | 89.3 |
|  | Tiberius | 0.839 | 0.735 | 0.778 | 0.856 | 0.602 | 0.419 | 94.8 |
|  | ANNEVO | 0.900 | 0.887 | 0.837 | 0.834 | 0.517 | 0.480 | 97.7 |
|  | Helixer | 0.872 | 0.867 | 0.801 | 0.730 | 0.346 | 0.308 | 90.5 |
|  | GeneMark | 0.879 | 0.487 | 0.643 | 0.487 | 0.174 | 0.089 | 76.9 |
| Cloeon dipterum<br>GCF_949628265.1<br>ref. BUSCO C 96.0 | Vipsania | 0.958 | 0.785 | 0.865 | 0.758 | 0.607 | 0.471 | 94.6 |
|  | Vipsania PT | 0.931 | 0.802 | 0.812 | 0.775 | 0.496 | 0.438 | 91.4 |
|  | Tiberius | 0.939 | 0.795 | 0.837 | 0.789 | 0.665 | 0.494 | 95.1 |
|  | ANNEVO | 0.957 | 0.848 | 0.837 | 0.756 | 0.534 | 0.433 | 94.8 |
|  | Helixer | 0.971 | 0.759 | 0.814 | 0.655 | 0.446 | 0.290 | 92.6 |
|  | GeneMark | 0.956 | 0.729 | 0.802 | 0.665 | 0.400 | 0.277 | 91.2 |

continued on next page

Table 13: Insecta (continued)

| Species | Tool | Base |  | Exon |  | Locus |  | BUSCO |
| --- | --- | --- | --- | --- | --- | --- | --- | --- |
|  |  | S | P | S | P | S | P |  |
| *Drosophila melanogaster*<br>GCF_000001215.4<br>ref. BUSCO C 98.8 | Vipsania | 0.956 | 0.991 | 0.807 | 0.938 | 0.766 | 0.793 | 97.5 |
|  | Vipsania PT | 0.913 | 0.992 | 0.748 | 0.926 | 0.616 | 0.739 | 93.1 |
|  | Tiberius | 0.937 | 0.919 | 0.811 | 0.907 | 0.795 | 0.695 | 97.8 |
|  | ANNEVO | 0.953 | 0.916 | 0.800 | 0.781 | 0.736 | 0.589 | 97.5 |
|  | Helixer | 0.959 | 0.965 | 0.792 | 0.843 | 0.688 | 0.664 | 97.0 |
|  | GeneMark | 0.913 | 0.825 | 0.680 | 0.632 | 0.507 | 0.411 | 93.1 |
| *Kerria lacca*<br>GCA_045014175.1<br>ref. BUSCO C 80.6 | Vipsania | 0.910 | 0.774 | 0.797 | 0.695 | 0.319 | 0.349 | 90.8 |
|  | Vipsania PT | 0.886 | 0.781 | 0.778 | 0.700 | 0.268 | 0.333 | 86.9 |
|  | Tiberius | 0.914 | 0.731 | 0.800 | 0.672 | 0.346 | 0.279 | 89.8 |
|  | ANNEVO | 0.918 | 0.766 | 0.788 | 0.644 | 0.280 | 0.272 | 91.0 |
|  | Helixer | 0.914 | 0.750 | 0.777 | 0.592 | 0.238 | 0.216 | 87.3 |
|  | GeneMark | 0.932 | 0.538 | 0.769 | 0.441 | 0.247 | 0.121 | 81.7 |
| *Tenebrio molitor*<br>GCF_963966145.1<br>ref. BUSCO C 98.9 | Vipsania | 0.909 | 0.876 | 0.789 | 0.821 | 0.583 | 0.538 | 96.3 |
|  | Vipsania PT | 0.895 | 0.880 | 0.753 | 0.810 | 0.502 | 0.489 | 93.0 |
|  | Tiberius | 0.904 | 0.772 | 0.773 | 0.767 | 0.625 | 0.414 | 95.2 |
|  | ANNEVO | 0.932 | 0.831 | 0.771 | 0.720 | 0.505 | 0.387 | 97.3 |
|  | Helixer | 0.931 | 0.777 | 0.608 | 0.504 | 0.257 | 0.168 | 90.1 |
|  | GeneMark | 0.818 | 0.507 | 0.554 | 0.343 | 0.240 | 0.115 | 71.3 |
| *Thermobia domestica*<br>GCF_964235325.1<br>ref. BUSCO C 92.0 | Vipsania | 0.589 | 0.788 | 0.618 | 0.789 | 0.220 | 0.333 | 72.9 |
|  | Vipsania PT | 0.567 | 0.794 | 0.604 | 0.776 | 0.188 | 0.284 | 69.6 |
|  | Tiberius | 0.579 | 0.328 | 0.577 | 0.441 | 0.164 | 0.044 | 61.7 |
|  | ANNEVO | 0.761 | 0.626 | 0.743 | 0.502 | 0.208 | 0.118 | 80.7 |
|  | Helixer | 0.703 | 0.485 | 0.653 | 0.383 | 0.123 | 0.070 | 60.0 |
|  | GeneMark | 0.571 | 0.073 | 0.052 | 0.010 | 0.077 | 0.009 | 10.2 |
| *Tribolium castaneum*<br>GCF_031307605.1<br>ref. BUSCO C 98.6 | Vipsania | 0.915 | 0.930 | 0.799 | 0.865 | 0.640 | 0.610 | 96.5 |
|  | Vipsania PT | 0.911 | 0.928 | 0.754 | 0.850 | 0.535 | 0.538 | 93.8 |
|  | Tiberius | 0.883 | 0.831 | 0.746 | 0.818 | 0.604 | 0.467 | 91.1 |
|  | ANNEVO | 0.929 | 0.909 | 0.761 | 0.775 | 0.522 | 0.467 | 96.1 |
|  | Helixer | 0.922 | 0.867 | 0.532 | 0.481 | 0.208 | 0.157 | 86.5 |
|  | GeneMark | 0.932 | 0.684 | 0.706 | 0.542 | 0.404 | 0.238 | 92.3 |

Table 14: Per-species sensitivity / precision for Nematoda. Rows list each tool's S and P per metric; – denotes a missing measurement. Accession IDs appear below each species name; the final column gives each tool's BUSCO Complete (C) percentage, and the reference annotation's BUSCO C is shown under the accession ID.

| Species | Tool | Base |  | Exon |  | Locus |  | BUSCO |
| --- | --- | --- | --- | --- | --- | --- | --- | --- |
|  |  | S | P | S | P | S | P |  |
| *Acanthocheilonema viteae*<br>GCA_046563165.1<br>ref. BUSCO C 98.0 | Vipsania | 0.967 | 0.924 | 0.906 | 0.907 | 0.603 | 0.612 | 99.2 |
|  | Vipsania PT | 0.962 | 0.921 | 0.897 | 0.901 | 0.560 | 0.579 | 98.8 |
|  | ANNEVO | 0.940 | 0.903 | 0.731 | 0.668 | 0.137 | 0.139 | 95.3 |
|  | Helixer | 0.951 | 0.824 | 0.710 | 0.550 | 0.084 | 0.072 | 93.1 |
|  | GeneMark | 0.949 | 0.860 | 0.822 | 0.782 | 0.250 | 0.252 | 97.8 |
| *Caenorhabditis elegans*<br>GCF_000002985.6<br>ref. BUSCO C 100.0 | Vipsania | 0.959 | 0.965 | 0.879 | 0.926 | 0.748 | 0.756 | 99.7 |
|  | Vipsania PT | 0.959 | 0.957 | 0.870 | 0.917 | 0.717 | 0.720 | 99.5 |
|  | ANNEVO | 0.809 | 0.966 | 0.595 | 0.710 | 0.212 | 0.295 | 88.9 |
|  | Helixer | 0.941 | 0.910 | 0.554 | 0.509 | 0.139 | 0.122 | 94.1 |
|  | GeneMark | 0.934 | 0.921 | 0.792 | 0.816 | 0.435 | 0.451 | 97.7 |
| *Meloidogyne hapla*<br>GCA_051171035.1<br>ref. BUSCO C 89.1 | Vipsania | 0.976 | 0.737 | 0.929 | 0.729 | 0.620 | 0.476 | 93.6 |
|  | Vipsania PT | 0.976 | 0.736 | 0.927 | 0.727 | 0.613 | 0.471 | 93.8 |
|  | ANNEVO | 0.796 | 0.806 | 0.609 | 0.578 | 0.102 | 0.115 | 78.7 |
|  | Helixer | 0.846 | 0.743 | 0.430 | 0.350 | 0.050 | 0.040 | 76.7 |
|  | GeneMark | 0.963 | 0.742 | 0.875 | 0.687 | 0.432 | 0.363 | 91.9 |
| *Necator americanus*<br>GCF_031761385.1<br>ref. BUSCO C 97.8 | Vipsania | 0.681 | 0.950 | 0.701 | 0.923 | 0.290 | 0.502 | 99.2 |
|  | Vipsania PT | 0.678 | 0.947 | 0.688 | 0.917 | 0.262 | 0.461 | 98.0 |
|  | ANNEVO | 0.433 | 0.923 | 0.340 | 0.593 | 0.022 | 0.053 | 61.4 |
|  | Helixer | 0.529 | 0.817 | 0.418 | 0.519 | 0.027 | 0.041 | 71.1 |
|  | GeneMark | 0.048 | 0.585 | 0.000 | 0.002 | 0.000 | 0.001 | 0.0 |
| *Nippostrongylus brasiliensis*<br>GCA_030553155.1<br>ref. BUSCO C 99.2 | Vipsania | 0.916 | 0.776 | 0.831 | 0.910 | 0.555 | 0.523 | 99.2 |
|  | Vipsania PT | 0.921 | 0.756 | 0.829 | 0.896 | 0.531 | 0.487 | 99.2 |
|  | ANNEVO | 0.644 | 0.914 | 0.454 | 0.628 | 0.054 | 0.078 | 73.2 |
|  | Helixer | 0.732 | 0.808 | 0.528 | 0.577 | 0.062 | 0.061 | 77.7 |
|  | GeneMark | 0.881 | 0.613 | 0.732 | 0.722 | 0.236 | 0.151 | 90.9 |

continued on next page

Table 14: Nematoda (continued)

| Species | Tool | Base |  | Exon |  | Locus |  | BUSCO |
| --- | --- | --- | --- | --- | --- | --- | --- | --- |
|  |  | S | P | S | P | S | P |  |
| Pristionchus pacificus<br>GCA_000180635.4<br>ref. BUSCO C 97.8 | Vipsania | 0.902 | 0.822 | 0.819 | 0.765 | 0.241 | 0.211 | 98.3 |
|  | Vipsania PT | 0.906 | 0.813 | 0.817 | 0.763 | 0.236 | 0.203 | 98.5 |
|  | ANNEVO | 0.611 | 0.835 | 0.442 | 0.555 | 0.028 | 0.038 | 72.5 |
|  | Helixer | 0.797 | 0.752 | 0.489 | 0.462 | 0.036 | 0.028 | 81.5 |
|  | GeneMark | 0.825 | 0.843 | 0.741 | 0.757 | 0.137 | 0.148 | 91.8 |
| Steinernema hermaphroditum<br>GCA_030435675.2<br>ref. BUSCO C 98.2 | Vipsania | 0.940 | 0.923 | 0.732 | 0.901 | 0.714 | 0.598 | 99.3 |
|  | Vipsania PT | 0.941 | 0.924 | 0.730 | 0.894 | 0.700 | 0.586 | 99.5 |
|  | ANNEVO | 0.641 | 0.967 | 0.317 | 0.573 | 0.117 | 0.160 | 79.9 |
|  | Helixer | 0.821 | 0.900 | 0.221 | 0.299 | 0.076 | 0.062 | 86.1 |
|  | GeneMark | 0.892 | 0.941 | 0.678 | 0.892 | 0.585 | 0.567 | 95.5 |
| Strongyloides ratti<br>GCF_001040885.1<br>ref. BUSCO C 96.1 | Vipsania | 0.980 | 0.960 | 0.743 | 0.811 | 0.641 | 0.623 | 97.5 |
|  | Vipsania PT | 0.978 | 0.961 | 0.734 | 0.801 | 0.619 | 0.616 | 96.3 |
|  | ANNEVO | 0.896 | 0.970 | 0.569 | 0.649 | 0.384 | 0.462 | 95.0 |
|  | Helixer | 0.934 | 0.961 | 0.470 | 0.517 | 0.261 | 0.272 | 95.5 |
|  | GeneMark | 0.980 | 0.967 | 0.719 | 0.780 | 0.582 | 0.614 | 97.0 |

Table 15: Per-species sensitivity / precision for Porifera. Rows list each tool's S and P per metric; – denotes a missing measurement. Accession IDs appear below each species name; the final column gives each tool's BUSCO Complete (C) percentage, and the reference annotation's BUSCO C is shown under the accession ID.

| Species | Tool | Base |  | Exon |  | Locus |  | BUSCO |
| --- | --- | --- | --- | --- | --- | --- | --- | --- |
|  |  | S | P | S | P | S | P |  |
| Corticium candelabrum<br>GCF_963422355.1<br>ref. BUSCO C 89.3 | Vipsania | 0.849 | 0.841 | 0.843 | 0.816 | 0.448 | 0.501 | 91.5 |
|  | Vipsania PT | 0.807 | 0.840 | 0.744 | 0.793 | 0.301 | 0.349 | 78.9 |
|  | ANNEVO | 0.551 | 0.850 | 0.312 | 0.481 | 0.069 | 0.118 | 22.6 |
|  | Helixer | 0.715 | 0.785 | 0.429 | 0.480 | 0.114 | 0.122 | 28.6 |
|  | GeneMark | 0.963 | 0.439 | 0.665 | 0.285 | 0.120 | 0.045 | 64.9 |
| Dysidea avara<br>GCF_963678975.1<br>ref. BUSCO C 90.0 | Vipsania | 0.844 | 0.828 | 0.826 | 0.851 | 0.496 | 0.520 | 89.3 |
|  | Vipsania PT | 0.846 | 0.819 | 0.827 | 0.828 | 0.476 | 0.475 | 88.7 |
|  | ANNEVO | 0.579 | 0.851 | 0.399 | 0.526 | 0.093 | 0.119 | 30.8 |
|  | Helixer | 0.721 | 0.761 | 0.450 | 0.476 | 0.130 | 0.121 | 29.0 |
|  | GeneMark | 0.943 | 0.490 | 0.729 | 0.482 | 0.246 | 0.110 | 69.2 |
| Ephydatia muelleri<br>GCA_049114765.1<br>ref. BUSCO C 78.9 | Vipsania | 0.604 | 0.844 | 0.601 | 0.767 | 0.160 | 0.261 | 89.7 |
|  | Vipsania PT | 0.606 | 0.835 | 0.598 | 0.753 | 0.151 | 0.233 | 87.2 |
|  | ANNEVO | 0.451 | 0.860 | 0.300 | 0.500 | 0.032 | 0.065 | 35.6 |
|  | Helixer | 0.533 | 0.816 | 0.350 | 0.493 | 0.041 | 0.064 | 37.6 |
|  | GeneMark | 0.658 | 0.547 | 0.118 | 0.063 | 0.023 | 0.013 | 20.5 |
| Halichondria panicea<br>GCF_963675165.1<br>ref. BUSCO C 89.0 | Vipsania | 0.937 | 0.865 | 0.883 | 0.848 | 0.616 | 0.551 | 88.4 |
|  | Vipsania PT | 0.941 | 0.852 | 0.887 | 0.819 | 0.609 | 0.504 | 88.5 |
|  | ANNEVO | 0.716 | 0.907 | 0.584 | 0.642 | 0.125 | 0.171 | 54.9 |
|  | Helixer | 0.903 | 0.824 | 0.695 | 0.584 | 0.228 | 0.184 | 64.0 |
|  | GeneMark | 0.973 | 0.755 | 0.839 | 0.666 | 0.403 | 0.266 | 81.4 |
| Oscarella lobularis<br>GCF_947507565.1<br>ref. BUSCO C 88.8 | Vipsania | 0.968 | 0.870 | 0.910 | 0.860 | 0.658 | 0.549 | 89.7 |
|  | Vipsania PT | 0.961 | 0.871 | 0.893 | 0.850 | 0.595 | 0.493 | 89.1 |
|  | ANNEVO | 0.594 | 0.914 | 0.313 | 0.539 | 0.072 | 0.107 | 25.0 |
|  | Helixer | 0.766 | 0.868 | 0.378 | 0.491 | 0.101 | 0.093 | 34.4 |
|  | GeneMark | 0.971 | 0.792 | 0.849 | 0.744 | 0.408 | 0.306 | 85.7 |
| Sycon ciliatum<br>GCF_964019385.1<br>ref. BUSCO C 86.3 | Vipsania | 0.821 | 0.818 | 0.879 | 0.875 | 0.527 | 0.520 | 85.3 |
|  | Vipsania PT | 0.814 | 0.790 | 0.859 | 0.837 | 0.449 | 0.404 | 83.6 |
|  | ANNEVO | 0.716 | 0.863 | 0.661 | 0.657 | 0.159 | 0.185 | 60.1 |
|  | Helixer | 0.758 | 0.690 | 0.626 | 0.553 | 0.180 | 0.144 | 51.6 |
|  | GeneMark | 0.876 | 0.374 | 0.627 | 0.395 | 0.188 | 0.065 | 53.6 |

Table 16: Per-species sensitivity / precision for Rhodophyta. Rows list each tool's S and P per metric; – denotes a missing measurement. Accession IDs appear below each species name; the final column gives each tool's BUSCO Complete (C) percentage, and the reference annotation's BUSCO C is shown under the accession ID.

| Species | Tool | Base |  | Exon |  | Locus |  | BUSCO |
| --- | --- | --- | --- | --- | --- | --- | --- | --- |
|  |  | S | P | S | P | S | P |  |
| Chondrus crispus<br>GCF_000350225.1<br>ref. BUSCO C 67.4 | Vipsania | 0.719 | 0.476 | 0.254 | 0.249 | 0.259 | 0.270 | 73.6 |
|  | Vipsania PT | 0.819 | 0.240 | 0.280 | 0.102 | 0.286 | 0.152 | 77.5 |
|  | GeneMark | 0.839 | 0.203 | 0.199 | 0.041 | 0.235 | 0.087 | 76.7 |

continued on next page

Table 16: Rhodophyta (continued)

| Species | Tool | Base |  | Exon |  | Locus |  | BUSCO |
| --- | --- | --- | --- | --- | --- | --- | --- | --- |
|  |  | S | P | S | P | S | P |  |
| Cyanidioschyzon merolae | Vipsania | 0.971 | 0.969 | 0.835 | 0.802 | 0.836 | 0.811 | 60.5 |
| GCF_000091205.1 | Vipsania PT | 0.868 | 0.967 | 0.544 | 0.576 | 0.544 | 0.643 | 52.7 |
| ref. BUSCO C 61.2 | GeneMark | 0.939 | 0.924 | 0.648 | 0.494 | 0.652 | 0.644 | 60.5 |
| Galdieria sulphuraria | Vipsania | 0.933 | 0.945 | 0.665 | 0.725 | 0.472 | 0.484 | 82.9 |
| GCF_000341285.1 | Vipsania PT | 0.824 | 0.944 | 0.357 | 0.533 | 0.228 | 0.319 | 69.8 |
| ref. BUSCO C 81.4 | GeneMark | 0.948 | 0.961 | 0.713 | 0.772 | 0.516 | 0.563 | 83.7 |
| Gracilaria domingensis | Vipsania | 0.813 | 0.395 | 0.242 | 0.125 | 0.307 | 0.202 | 83.7 |
| GCA_022539475.1 | Vipsania PT | 0.816 | 0.386 | 0.240 | 0.122 | 0.305 | 0.197 | 82.2 |
| ref. BUSCO C 68.2 | GeneMark | 0.837 | 0.371 | 0.224 | 0.087 | 0.296 | 0.173 | 79.1 |
| Porphyra umbilicalis | Vipsania | 0.680 | 0.364 | 0.263 | 0.135 | 0.266 | 0.182 | 48.8 |
| GCA_002049455.2 | Vipsania PT | 0.646 | 0.362 | 0.254 | 0.137 | 0.256 | 0.183 | 48.1 |
| ref. BUSCO C 37.2 | GeneMark | 0.663 | 0.411 | 0.118 | 0.040 | 0.118 | 0.085 | 45.7 |

Table 17: Per-species sensitivity / precision for Spiralia. Rows list each tool's S and P per metric; – denotes a missing measurement. Accession IDs appear below each species name; the final column gives each tool's BUSCO Complete (C) percentage, and the reference annotation's BUSCO C is shown under the accession ID.

| Species | Tool | Base |  | Exon |  | Locus |  | BUSCO |
| --- | --- | --- | --- | --- | --- | --- | --- | --- |
|  |  | S | P | S | P | S | P |  |
| Adineta vaga | Vipsania | 0.943 | 0.862 | 0.833 | 0.816 | 0.578 | 0.556 | 94.3 |
| GCA_021613535.1 | Vipsania PT | 0.618 | 0.846 | 0.344 | 0.690 | 0.136 | 0.278 | 35.0 |
| ref. BUSCO C 91.4 | ANNEVO | 0.730 | 0.890 | 0.599 | 0.678 | 0.196 | 0.294 | 80.1 |
|  | Helixer | 0.952 | 0.825 | 0.678 | 0.599 | 0.316 | 0.276 | 90.3 |
|  | GeneMark | 0.970 | 0.849 | 0.848 | 0.794 | 0.555 | 0.540 | 93.6 |
| Neoechinorhynchus agilis | Vipsania | 0.700 | 0.907 | 0.434 | 0.624 | 0.229 | 0.370 | 22.8 |
| GCA_051530055.1 | Vipsania PT | 0.421 | 0.916 | 0.175 | 0.520 | 0.083 | 0.283 | 8.9 |
| ref. BUSCO C 28.4 | ANNEVO | 0.297 | 0.920 | 0.174 | 0.447 | 0.051 | 0.219 | 7.4 |
|  | Helixer | 0.618 | 0.858 | 0.222 | 0.262 | 0.125 | 0.198 | 19.5 |
|  | GeneMark | 0.933 | 0.670 | 0.575 | 0.404 | 0.341 | 0.302 | 33.6 |
| Octopus bimaculoides | Vipsania | 0.707 | 0.935 | 0.650 | 0.886 | 0.198 | 0.325 | 73.7 |
| GCF_001194135.2 | Vipsania PT | 0.795 | 0.921 | 0.733 | 0.858 | 0.263 | 0.352 | 84.8 |
| ref. BUSCO C 93.0 | ANNEVO | 0.837 | 0.917 | 0.724 | 0.684 | 0.189 | 0.188 | 79.9 |
|  | Helixer | 0.866 | 0.894 | 0.762 | 0.711 | 0.241 | 0.240 | 83.9 |
|  | GeneMark | 0.222 | 0.090 | 0.008 | 0.001 | 0.031 | 0.005 | 4.2 |
| Patella vulgata | Vipsania | 0.924 | 0.716 | 0.854 | 0.721 | 0.490 | 0.382 | 93.5 |
| GCF_932274485.2 | Vipsania PT | 0.916 | 0.719 | 0.843 | 0.716 | 0.443 | 0.367 | 93.6 |
| ref. BUSCO C 89.7 | ANNEVO | 0.893 | 0.737 | 0.794 | 0.636 | 0.299 | 0.229 | 85.9 |
|  | Helixer | 0.950 | 0.630 | 0.823 | 0.491 | 0.221 | 0.121 | 83.8 |
|  | GeneMark | 0.941 | 0.503 | 0.786 | 0.563 | 0.281 | 0.122 | 86.9 |
| Tubulanus polymorphus | Vipsania | 0.906 | 0.738 | 0.853 | 0.807 | 0.457 | 0.390 | 91.8 |
| GCF_964204645.1 | Vipsania PT | 0.883 | 0.719 | 0.801 | 0.796 | 0.331 | 0.324 | 86.6 |
| ref. BUSCO C 99.0 | ANNEVO | 0.896 | 0.795 | 0.795 | 0.740 | 0.305 | 0.284 | 88.2 |
|  | Helixer | 0.935 | 0.683 | 0.828 | 0.629 | 0.273 | 0.183 | 91.1 |
|  | GeneMark | 0.948 | 0.582 | 0.823 | 0.644 | 0.361 | 0.189 | 88.5 |
| Watersipora subatra | Vipsania | 0.823 | 0.672 | 0.812 | 0.668 | 0.312 | 0.289 | 91.5 |
| GCF_963576615.1 | Vipsania PT | 0.798 | 0.652 | 0.781 | 0.638 | 0.233 | 0.230 | 86.8 |
| ref. BUSCO C 98.7 | ANNEVO | 0.788 | 0.660 | 0.709 | 0.513 | 0.125 | 0.102 | 70.7 |
|  | Helixer | 0.784 | 0.516 | 0.655 | 0.382 | 0.082 | 0.045 | 49.0 |
|  | GeneMark | 0.886 | 0.320 | 0.647 | 0.304 | 0.110 | 0.037 | 59.4 |

Table 18: Per-species sensitivity / precision for Stramenopiles. Rows list each tool's S and P per metric; – denotes a missing measurement. Accession IDs appear below each species name; the final column gives each tool's BUSCO Complete (C) percentage, and the reference annotation's BUSCO C is shown under the accession ID.

| Species | Tool | Base |  | Exon |  | Locus |  | BUSCO |
| --- | --- | --- | --- | --- | --- | --- | --- | --- |
|  |  | S | P | S | P | S | P |  |
| Bremia lactucae | Vipsania | 0.922 | 0.900 | 0.657 | 0.699 | 0.529 | 0.597 | 96.8 |
| GCF_004359215.1 | Vipsania PT | 0.907 | 0.898 | 0.622 | 0.680 | 0.496 | 0.572 | 95.8 |
| ref. BUSCO C 91.4 | Tiberius | 0.785 | 0.793 | 0.308 | 0.441 | 0.375 | 0.360 | 75.3 |
|  | GeneMark | 0.948 | 0.616 | 0.588 | 0.281 | 0.453 | 0.328 | 96.8 |

continued on next page

Table 18: Stramenopiles (continued)

| Species | Tool | Base |  | Exon |  | Locus |  | BUSCO |
| --- | --- | --- | --- | --- | --- | --- | --- | --- |
|  |  | S | P | S | P | S | P |  |
| Ectocarpus siliculosus<br>GCA_000310025.1<br>ref. BUSCO C 91.8 | Vipsania | 0.924 | 0.775 | 0.782 | 0.741 | 0.246 | 0.262 | 94.5 |
|  | Vipsania PT | 0.900 | 0.793 | 0.758 | 0.758 | 0.218 | 0.261 | 93.5 |
|  | Tiberius | 0.395 | 0.578 | 0.083 | 0.230 | 0.025 | 0.019 | 5.9 |
|  | GeneMark | 0.922 | 0.684 | 0.727 | 0.621 | 0.137 | 0.113 | 90.8 |
| Peronosclerospora sorghi<br>GCA_026184515.1<br>ref. BUSCO C 86.7 | Vipsania | 0.527 | 0.812 | 0.209 | 0.481 | 0.147 | 0.392 | 79.6 |
|  | Vipsania PT | 0.584 | 0.767 | 0.222 | 0.413 | 0.161 | 0.333 | 86.7 |
|  | Tiberius | 0.697 | 0.418 | 0.181 | 0.164 | 0.193 | 0.111 | 81.5 |
|  | GeneMark | 0.807 | 0.277 | 0.154 | 0.043 | 0.126 | 0.068 | 89.5 |
| Phaeodactylum tricornutum<br>GCF_000150955.2<br>ref. BUSCO C 89.8 | Vipsania | 0.955 | 0.807 | 0.523 | 0.542 | 0.518 | 0.481 | 99.1 |
|  | Vipsania PT | 0.960 | 0.805 | 0.522 | 0.539 | 0.516 | 0.479 | 99.3 |
|  | Tiberius | 0.946 | 0.801 | 0.499 | 0.515 | 0.489 | 0.446 | 98.3 |
|  | GeneMark | 0.968 | 0.800 | 0.497 | 0.504 | 0.492 | 0.446 | 99.0 |
| Phytophthora ramorum<br>GCF_020800215.1<br>ref. BUSCO C 99.6 | Vipsania | 0.956 | 0.802 | 0.783 | 0.763 | 0.706 | 0.637 | 99.4 |
|  | Vipsania PT | 0.952 | 0.744 | 0.750 | 0.738 | 0.669 | 0.596 | 99.3 |
|  | Tiberius | 0.892 | 0.686 | 0.469 | 0.518 | 0.491 | 0.382 | 87.5 |
|  | GeneMark | 0.953 | 0.717 | 0.726 | 0.583 | 0.632 | 0.497 | 99.1 |
| Thalassiosira pseudonana ccmp1335<br>GCF_000149405.2<br>ref. BUSCO C 89.5 | Vipsania | 0.969 | 0.739 | 0.497 | 0.461 | 0.391 | 0.337 | 99.1 |
|  | Vipsania PT | 0.965 | 0.741 | 0.496 | 0.465 | 0.392 | 0.340 | 99.1 |
|  | Tiberius | 0.950 | 0.745 | 0.484 | 0.470 | 0.393 | 0.343 | 98.3 |
|  | GeneMark | 0.971 | 0.742 | 0.484 | 0.451 | 0.374 | 0.325 | 98.1 |

Table 19: Per-species sensitivity / precision for Streptophyta. Rows list each tool's S and P per metric; – denotes a missing measurement. Accession IDs appear below each species name; the final column gives each tool's BUSCO Complete (C) percentage, and the reference annotation's BUSCO C is shown under the accession ID.

| Species | Tool | Base |  | Exon |  | Locus |  | BUSCO |
| --- | --- | --- | --- | --- | --- | --- | --- | --- |
|  |  | S | P | S | P | S | P |  |
| Arabidopsis thaliana<br>GCF_000001735.4<br>ref. BUSCO C 98.9 | Vipsania | 0.933 | 0.984 | 0.826 | 0.958 | 0.763 | 0.839 | 98.4 |
|  | Vipsania PT | 0.916 | 0.985 | 0.807 | 0.956 | 0.703 | 0.810 | 96.8 |
|  | Tiberius | 0.935 | 0.978 | 0.824 | 0.960 | 0.785 | 0.863 | 98.3 |
|  | ANNEVO | 0.926 | 0.967 | 0.824 | 0.960 | 0.795 | 0.878 | 98.2 |
|  | Helixer | 0.955 | 0.953 | 0.822 | 0.895 | 0.734 | 0.743 | 97.9 |
|  | GeneMark | 0.956 | 0.778 | 0.777 | 0.724 | 0.554 | 0.476 | 97.0 |
| Asparagus officinalis<br>GCF_001876935.1<br>ref. BUSCO C 88.9 | Vipsania | 0.617 | 0.942 | 0.620 | 0.877 | 0.297 | 0.505 | 70.4 |
|  | Vipsania PT | 0.668 | 0.925 | 0.648 | 0.853 | 0.326 | 0.467 | 74.0 |
|  | Tiberius | 0.741 | 0.850 | 0.675 | 0.768 | 0.375 | 0.346 | 71.9 |
|  | ANNEVO | 0.776 | 0.726 | 0.714 | 0.700 | 0.457 | 0.346 | 76.8 |
|  | Helixer | 0.781 | 0.709 | 0.700 | 0.545 | 0.275 | 0.206 | 79.2 |
|  | GeneMark | 0.596 | 0.121 | 0.124 | 0.024 | 0.052 | 0.010 | 12.9 |
| Ceratodon purpureus<br>GCA_014871385.1<br>ref. BUSCO C 80.9 | Vipsania | 0.826 | 0.916 | 0.758 | 0.870 | 0.415 | 0.646 | 81.0 |
|  | Vipsania PT | 0.812 | 0.866 | 0.747 | 0.837 | 0.393 | 0.590 | 80.5 |
|  | Tiberius | 0.731 | 0.818 | 0.633 | 0.771 | 0.316 | 0.398 | 66.5 |
|  | ANNEVO | 0.811 | 0.644 | 0.719 | 0.719 | 0.407 | 0.442 | 75.8 |
|  | Helixer | 0.861 | 0.522 | 0.733 | 0.489 | 0.327 | 0.251 | 78.3 |
|  | GeneMark | 0.848 | 0.306 | 0.569 | 0.215 | 0.117 | 0.056 | 72.1 |
| Diphysiastrum complanatum<br>GCA_029204225.1<br>ref. BUSCO C 85.5 | Vipsania | 0.775 | 0.912 | 0.631 | 0.850 | 0.402 | 0.530 | 62.1 |
|  | Vipsania PT | 0.754 | 0.906 | 0.556 | 0.828 | 0.372 | 0.488 | 44.1 |
|  | Tiberius | 0.711 | 0.800 | 0.476 | 0.658 | 0.304 | 0.254 | 34.5 |
|  | ANNEVO | 0.900 | 0.820 | 0.728 | 0.729 | 0.484 | 0.408 | 68.8 |
|  | Helixer | 0.801 | 0.807 | 0.596 | 0.547 | 0.268 | 0.240 | 50.8 |
|  | GeneMark | 0.787 | 0.271 | 0.326 | 0.116 | 0.145 | 0.036 | 23.3 |
| Hordeum vulgare<br>GCF_904849725.1<br>ref. BUSCO C 96.7 | Vipsania | 0.758 | 0.921 | 0.763 | 0.883 | 0.496 | 0.670 | 87.4 |
|  | Vipsania PT | 0.831 | 0.899 | 0.782 | 0.855 | 0.583 | 0.633 | 89.9 |
|  | Tiberius | 0.816 | 0.837 | 0.643 | 0.695 | 0.448 | 0.315 | 56.0 |
|  | ANNEVO | 0.929 | 0.783 | 0.856 | 0.760 | 0.741 | 0.558 | 96.3 |
|  | Helixer | 0.932 | 0.700 | 0.832 | 0.581 | 0.628 | 0.382 | 95.1 |
|  | GeneMark | 0.389 | 0.063 | 0.034 | 0.003 | 0.054 | 0.006 | 9.6 |
| Juncus effusus<br>GCA_027726005.1<br>ref. BUSCO C 88.6 | Vipsania | 0.914 | 0.788 | 0.899 | 0.772 | 0.655 | 0.551 | 96.9 |
|  | Vipsania PT | 0.901 | 0.798 | 0.886 | 0.777 | 0.622 | 0.545 | 95.2 |
|  | Tiberius | 0.930 | 0.748 | 0.859 | 0.760 | 0.642 | 0.471 | 88.4 |
|  | ANNEVO | 0.931 | 0.754 | 0.896 | 0.772 | 0.691 | 0.539 | 94.4 |
|  | Helixer | 0.936 | 0.709 | 0.881 | 0.652 | 0.587 | 0.389 | 92.8 |
|  | GeneMark | 0.962 | 0.402 | 0.819 | 0.335 | 0.386 | 0.139 | 90.1 |

continued on next page

Table 19: Streptophyta (continued)

| Species | Tool | Base |  | Exon |  | Locus |  | BUSCO C% |
| --- | --- | --- | --- | --- | --- | --- | --- | --- |
|  |  | S | P | S | P | S | P |  |
| Marchantia polymorpha subsp. ruderalis<br>GCA_037833965.1<br>ref. BUSCO C 86.0 | Vipsania | 0.889 | 0.955 | 0.835 | 0.928 | 0.537 | 0.717 | 83.9 |
|  | Vipsania PT | 0.861 | 0.958 | 0.807 | 0.919 | 0.482 | 0.658 | 81.6 |
|  | Tiberius | 0.616 | 0.935 | 0.503 | 0.776 | 0.225 | 0.321 | 38.9 |
|  | ANNEVO | 0.826 | 0.935 | 0.737 | 0.865 | 0.411 | 0.545 | 70.3 |
|  | Helixer | 0.891 | 0.919 | 0.792 | 0.761 | 0.396 | 0.469 | 80.4 |
|  | GeneMark | 0.656 | 0.321 | 0.124 | 0.045 | 0.037 | 0.019 | 21.4 |
| Physcomitrium patens<br>GCF_000002425.5<br>ref. BUSCO C 82.7 | Vipsania | 0.924 | 0.967 | 0.818 | 0.933 | 0.637 | 0.705 | 81.0 |
|  | Vipsania PT | 0.888 | 0.970 | 0.788 | 0.931 | 0.564 | 0.662 | 78.8 |
|  | Tiberius | 0.804 | 0.948 | 0.661 | 0.862 | 0.420 | 0.470 | 68.0 |
|  | ANNEVO | 0.855 | 0.958 | 0.729 | 0.895 | 0.529 | 0.583 | 74.8 |
|  | Helixer | 0.916 | 0.934 | 0.752 | 0.768 | 0.416 | 0.423 | 77.5 |
|  | GeneMark | 0.934 | 0.456 | 0.434 | 0.151 | 0.012 | 0.007 | 68.5 |
| Solanum lycopersicum<br>GCF_036512215.1<br>ref. BUSCO C 98.6 | Vipsania | 0.830 | 0.939 | 0.808 | 0.907 | 0.636 | 0.711 | 98.0 |
|  | Vipsania PT | 0.809 | 0.952 | 0.788 | 0.919 | 0.590 | 0.711 | 95.2 |
|  | Tiberius | 0.836 | 0.904 | 0.810 | 0.891 | 0.673 | 0.663 | 97.9 |
|  | ANNEVO | 0.859 | 0.930 | 0.834 | 0.915 | 0.719 | 0.762 | 98.5 |
|  | Helixer | 0.849 | 0.777 | 0.806 | 0.683 | 0.547 | 0.372 | 97.1 |
|  | GeneMark | 0.900 | 0.320 | 0.578 | 0.170 | 0.218 | 0.073 | 71.9 |
| Solanum tuberosum<br>GCF_000226075.1<br>ref. BUSCO C 97.5 | Vipsania | 0.886 | 0.891 | 0.836 | 0.861 | 0.642 | 0.642 | 96.1 |
|  | Vipsania PT | 0.857 | 0.910 | 0.814 | 0.877 | 0.596 | 0.652 | 93.1 |
|  | Tiberius | 0.892 | 0.859 | 0.840 | 0.843 | 0.682 | 0.604 | 95.6 |
|  | ANNEVO | 0.912 | 0.865 | 0.863 | 0.842 | 0.729 | 0.621 | 96.6 |
|  | Helixer | 0.913 | 0.776 | 0.827 | 0.641 | 0.542 | 0.380 | 93.6 |
|  | GeneMark | 0.890 | 0.286 | 0.533 | 0.166 | 0.214 | 0.069 | 63.2 |
| Sorghum bicolor<br>GCF_000003195.3<br>ref. BUSCO C 96.3 | Vipsania | 0.822 | 0.953 | 0.806 | 0.920 | 0.601 | 0.726 | 92.3 |
|  | Vipsania PT | 0.851 | 0.941 | 0.805 | 0.909 | 0.617 | 0.697 | 92.3 |
|  | Tiberius | 0.880 | 0.925 | 0.808 | 0.882 | 0.666 | 0.638 | 86.8 |
|  | ANNEVO | 0.913 | 0.888 | 0.857 | 0.882 | 0.750 | 0.712 | 96.0 |
|  | Helixer | 0.914 | 0.885 | 0.836 | 0.794 | 0.637 | 0.574 | 95.3 |
|  | GeneMark | 0.450 | 0.240 | 0.048 | 0.015 | 0.036 | 0.015 | 10.6 |
| Vitis vinifera<br>GCF_030704535.1<br>ref. BUSCO C 98.8 | Vipsania | 0.867 | 0.951 | 0.811 | 0.905 | 0.605 | 0.675 | 94.5 |
|  | Vipsania PT | 0.871 | 0.947 | 0.805 | 0.898 | 0.593 | 0.654 | 92.6 |
|  | Tiberius | 0.927 | 0.895 | 0.858 | 0.876 | 0.737 | 0.658 | 98.8 |
|  | ANNEVO | 0.930 | 0.927 | 0.866 | 0.902 | 0.756 | 0.732 | 98.9 |
|  | Helixer | 0.923 | 0.894 | 0.828 | 0.761 | 0.542 | 0.489 | 95.7 |
|  | GeneMark | 0.910 | 0.372 | 0.544 | 0.208 | 0.187 | 0.073 | 60.8 |
| Zea mays<br>GCF_902167145.1<br>ref. BUSCO C 94.8 | Vipsania | 0.803 | 0.937 | 0.753 | 0.900 | 0.571 | 0.717 | 90.4 |
|  | Vipsania PT | 0.806 | 0.922 | 0.739 | 0.872 | 0.550 | 0.661 | 88.0 |
|  | Tiberius | 0.014 | 0.410 | 0.004 | 0.249 | 0.008 | 0.136 | 0.2 |
|  | ANNEVO | 0.901 | 0.678 | 0.820 | 0.742 | 0.700 | 0.549 | 95.8 |
|  | Helixer | 0.881 | 0.794 | 0.785 | 0.655 | 0.569 | 0.466 | 94.0 |
|  | GeneMark | 0.356 | 0.118 | 0.029 | 0.004 | 0.038 | 0.009 | 9.2 |

Table 20: Per-species sensitivity / precision for Tunicata. Rows list each tool's S and P per metric; – denotes a missing measurement. Accession IDs appear below each species name; the final column gives each tool's BUSCO Complete (C) percentage, and the reference annotation's BUSCO C is shown under the accession ID.

| Species | Tool | Base |  | Exon |  | Locus |  | BUSCO C% |
| --- | --- | --- | --- | --- | --- | --- | --- | --- |
|  |  | S | P | S | P | S | P |  |
| Botryllus schlosseri<br>GCF_051294905.1<br>ref. BUSCO C 95.8 | Vipsania | 0.889 | 0.847 | 0.855 | 0.850 | 0.534 | 0.521 | 94.0 |
|  | Vipsania PT | 0.888 | 0.831 | 0.835 | 0.830 | 0.491 | 0.456 | 93.2 |
|  | ANNEVO | 0.725 | 0.880 | 0.609 | 0.657 | 0.160 | 0.187 | 64.6 |
|  | Helixer | 0.824 | 0.742 | 0.665 | 0.521 | 0.185 | 0.139 | 76.9 |
|  | GeneMark | 0.918 | 0.611 | 0.756 | 0.677 | 0.332 | 0.204 | 90.0 |
| Ciona intestinalis<br>GCF_018327825.1<br>ref. BUSCO C 94.3 | Vipsania | 0.949 | 0.881 | 0.893 | 0.837 | 0.553 | 0.519 | 93.2 |
|  | Vipsania PT | 0.942 | 0.878 | 0.882 | 0.839 | 0.515 | 0.503 | 92.4 |
|  | ANNEVO | 0.887 | 0.897 | 0.777 | 0.729 | 0.236 | 0.264 | 77.1 |
|  | Helixer | 0.952 | 0.818 | 0.824 | 0.643 | 0.244 | 0.193 | 86.0 |
|  | GeneMark | 0.969 | 0.727 | 0.842 | 0.665 | 0.309 | 0.213 | 88.2 |
| Clavelina lepadiformis<br>GCF_947623445.1<br>ref. BUSCO C 96.1 | Vipsania | 0.953 | 0.906 | 0.900 | 0.901 | 0.639 | 0.609 | 95.7 |
|  | Vipsania PT | 0.937 | 0.912 | 0.876 | 0.907 | 0.563 | 0.571 | 93.9 |
|  | ANNEVO | 0.885 | 0.913 | 0.749 | 0.742 | 0.231 | 0.240 | 81.1 |
|  | Helixer | 0.962 | 0.795 | 0.795 | 0.616 | 0.240 | 0.160 | 85.9 |
|  | GeneMark | 0.939 | 0.761 | 0.809 | 0.758 | 0.338 | 0.252 | 83.8 |

continued on next page

Table 20: Tunicata (continued)

| Species | Tool | Base |  | Exon |  | Locus |  | BUSCO |
| --- | --- | --- | --- | --- | --- | --- | --- | --- |
|  |  | S | P | S | P | S | P |  |
| Oikopleura dioica<br>GCA_907165135.1<br>ref. BUSCO C 62.2 | Vipsania | 0.927 | 0.829 | 0.702 | 0.643 | 0.331 | 0.260 | 67.1 |
|  | Vipsania PT | 0.915 | 0.826 | 0.668 | 0.645 | 0.307 | 0.247 | 62.2 |
|  | ANNEVO | 0.670 | 0.905 | 0.352 | 0.500 | 0.069 | 0.104 | 45.8 |
|  | Helixer | 0.862 | 0.818 | 0.254 | 0.249 | 0.042 | 0.034 | 45.1 |
|  | GeneMark | 0.937 | 0.821 | 0.657 | 0.639 | 0.269 | 0.248 | 62.9 |
| Phallusia mammillata<br>GCF_965637545.1<br>ref. BUSCO C 95.7 | Vipsania | 0.903 | 0.803 | 0.894 | 0.787 | 0.572 | 0.501 | 95.1 |
|  | Vipsania PT | 0.903 | 0.803 | 0.892 | 0.788 | 0.568 | 0.494 | 95.4 |
|  | ANNEVO | 0.855 | 0.837 | 0.774 | 0.690 | 0.238 | 0.247 | 85.3 |
|  | Helixer | 0.928 | 0.737 | 0.824 | 0.588 | 0.250 | 0.170 | 86.2 |
|  | GeneMark | 0.958 | 0.698 | 0.844 | 0.681 | 0.374 | 0.260 | 88.7 |
| Styela clava<br>GCF_964204865.1<br>ref. BUSCO C 97.5 | Vipsania | 0.931 | 0.872 | 0.912 | 0.875 | 0.666 | 0.591 | 97.3 |
|  | Vipsania PT | 0.929 | 0.868 | 0.907 | 0.875 | 0.651 | 0.583 | 97.5 |
|  | ANNEVO | 0.845 | 0.905 | 0.793 | 0.769 | 0.289 | 0.317 | 77.8 |
|  | Helixer | 0.949 | 0.749 | 0.839 | 0.553 | 0.242 | 0.154 | 87.9 |
|  | GeneMark | 0.962 | 0.738 | 0.859 | 0.776 | 0.446 | 0.321 | 94.0 |

Table 21: Per-species sensitivity / precision for Vertebrata. Rows list each tool's S and P per metric; – denotes a missing measurement. Accession IDs appear below each species name; the final column gives each tool's BUSCO Complete (C) percentage, and the reference annotation's BUSCO C is shown under the accession ID.

| Species | Tool | Base |  | Exon |  | Locus |  | BUSCO |
| --- | --- | --- | --- | --- | --- | --- | --- | --- |
|  |  | S | P | S | P | S | P |  |
| Danio rerio<br>GCF_049306965.1<br>ref. BUSCO C 98.5 | Vipsania | 0.898 | 0.876 | 0.802 | 0.906 | 0.530 | 0.525 | 92.6 |
|  | Vipsania PT | 0.926 | 0.668 | 0.827 | 0.840 | 0.550 | 0.383 | 94.9 |
|  | Tiberius | 0.914 | 0.946 | 0.851 | 0.919 | 0.654 | 0.596 | 94.2 |
|  | ANNEVO | 0.883 | 0.957 | 0.807 | 0.842 | 0.435 | 0.431 | 95.7 |
|  | Helixer | 0.889 | 0.909 | 0.786 | 0.703 | 0.242 | 0.199 | 85.5 |
|  | GeneMark | 0.877 | 0.458 | 0.656 | 0.342 | 0.106 | 0.034 | 74.2 |
| Delphinus delphis<br>GCF_949987515.2<br>ref. BUSCO C 96.3 | Vipsania | 0.784 | 0.729 | 0.704 | 0.893 | 0.408 | 0.467 | 81.2 |
|  | Vipsania PT | 0.785 | 0.470 | 0.698 | 0.827 | 0.368 | 0.386 | 81.7 |
|  | Tiberius | 0.923 | 0.852 | 0.855 | 0.904 | 0.610 | 0.559 | 95.1 |
|  | ANNEVO | 0.927 | 0.945 | 0.840 | 0.819 | 0.431 | 0.377 | 95.5 |
|  | Helixer | 0.903 | 0.928 | 0.797 | 0.723 | 0.237 | 0.231 | 91.7 |
|  | GeneMark | 0.364 | 0.092 | 0.008 | 0.002 | 0.006 | 0.001 | 7.5 |
| Denticeps clupeioides<br>GCF_900700375.2<br>ref. BUSCO C 96.2 | Vipsania | 0.950 | 0.897 | 0.867 | 0.893 | 0.576 | 0.510 | 95.9 |
|  | Vipsania PT | 0.949 | 0.849 | 0.854 | 0.860 | 0.528 | 0.419 | 95.1 |
|  | Tiberius | 0.960 | 0.937 | 0.888 | 0.915 | 0.699 | 0.634 | 96.6 |
|  | ANNEVO | 0.946 | 0.945 | 0.841 | 0.835 | 0.444 | 0.418 | 96.1 |
|  | Helixer | 0.925 | 0.902 | 0.805 | 0.738 | 0.280 | 0.244 | 91.9 |
|  | GeneMark | 0.825 | 0.498 | 0.574 | 0.325 | 0.054 | 0.020 | 65.7 |
| Gallus gallus<br>GCF_016699485.2<br>ref. BUSCO C 96.1 | Vipsania | 0.862 | 0.904 | 0.770 | 0.881 | 0.479 | 0.484 | 91.1 |
|  | Vipsania PT | 0.840 | 0.887 | 0.744 | 0.874 | 0.427 | 0.449 | 89.4 |
|  | Tiberius | 0.900 | 0.938 | 0.825 | 0.899 | 0.600 | 0.571 | 95.1 |
|  | ANNEVO | 0.877 | 0.965 | 0.780 | 0.844 | 0.392 | 0.412 | 96.7 |
|  | Helixer | 0.884 | 0.903 | 0.776 | 0.727 | 0.266 | 0.260 | 92.7 |
|  | GeneMark | 0.239 | 0.165 | 0.005 | 0.002 | 0.002 | 0.001 | 4.5 |
| Heptranchias perlo<br>GCF_035084215.1<br>ref. BUSCO C 96.5 | Vipsania | 0.739 | 0.766 | 0.663 | 0.827 | 0.344 | 0.336 | 78.8 |
|  | Vipsania PT | 0.756 | 0.651 | 0.680 | 0.710 | 0.306 | 0.223 | 80.5 |
|  | Tiberius | 0.894 | 0.786 | 0.834 | 0.775 | 0.460 | 0.285 | 90.2 |
|  | ANNEVO | 0.890 | 0.883 | 0.818 | 0.720 | 0.301 | 0.238 | 91.2 |
|  | Helixer | 0.808 | 0.761 | 0.740 | 0.544 | 0.108 | 0.098 | 79.2 |
|  | GeneMark | 0.018 | 0.019 | 0.000 | 0.000 | 0.001 | 0.000 | 0.1 |
| Homo sapiens<br>GCF_000001405.40<br>ref. BUSCO C 95.7 | Vipsania | 0.754 | 0.889 | 0.609 | 0.878 | 0.450 | 0.492 | 80.1 |
|  | Vipsania PT | 0.752 | 0.839 | 0.606 | 0.844 | 0.407 | 0.418 | 79.5 |
|  | Tiberius | 0.906 | 0.912 | 0.761 | 0.891 | 0.678 | 0.608 | 95.1 |
|  | ANNEVO | 0.918 | 0.916 | 0.748 | 0.798 | 0.469 | 0.394 | 94.9 |
|  | Helixer | 0.861 | 0.885 | 0.695 | 0.678 | 0.252 | 0.237 | 90.7 |
|  | GeneMark | 0.332 | 0.095 | 0.007 | 0.002 | 0.008 | 0.001 | 6.4 |
| Kryptolebias marmoratus<br>GCF_001649575.2<br>ref. BUSCO C 97.7 | Vipsania | 0.949 | 0.928 | 0.874 | 0.921 | 0.584 | 0.551 | 94.9 |
|  | Vipsania PT | 0.954 | 0.846 | 0.872 | 0.855 | 0.542 | 0.397 | 95.0 |
|  | Tiberius | 0.959 | 0.958 | 0.902 | 0.934 | 0.700 | 0.649 | 95.8 |
|  | ANNEVO | 0.950 | 0.966 | 0.868 | 0.862 | 0.479 | 0.459 | 95.8 |
|  | Helixer | 0.936 | 0.935 | 0.837 | 0.768 | 0.289 | 0.265 | 91.7 |
|  | GeneMark | 0.869 | 0.553 | 0.655 | 0.387 | 0.082 | 0.034 | 75.6 |

continued on next page

Table 21: Vertebrata (continued)

| Species | Tool | Base |  | Exon |  | Locus |  | BUSCO C% |
| --- | --- | --- | --- | --- | --- | --- | --- | --- |
|  |  | S | P | S | P | S | P |  |
| Latimeria chalumnae<br>GCF_037176945.1<br>ref. BUSCO C 94.4 | Vipsania | 0.828 | 0.331 | 0.743 | 0.673 | 0.361 | 0.137 | 81.2 |
|  | Vipsania PT | 0.847 | 0.202 | 0.757 | 0.467 | 0.312 | 0.065 | 81.5 |
|  | Tiberius | 0.918 | 0.833 | 0.857 | 0.777 | 0.458 | 0.335 | 88.8 |
|  | ANNEVO | 0.908 | 0.798 | 0.801 | 0.588 | 0.205 | 0.131 | 77.8 |
|  | Helixer | 0.851 | 0.579 | 0.755 | 0.367 | 0.067 | 0.031 | 66.1 |
|  | GeneMark | 0.447 | 0.156 | 0.102 | 0.054 | 0.044 | 0.010 | 11.2 |
| Mus musculus<br>GCF_000001635.27<br>ref. BUSCO C 95.3 | Vipsania | 0.786 | 0.795 | 0.649 | 0.885 | 0.430 | 0.449 | 79.4 |
|  | Vipsania PT | 0.802 | 0.622 | 0.658 | 0.827 | 0.407 | 0.333 | 80.6 |
|  | Tiberius | 0.922 | 0.933 | 0.802 | 0.929 | 0.688 | 0.643 | 94.2 |
|  | ANNEVO | 0.935 | 0.923 | 0.788 | 0.819 | 0.491 | 0.419 | 93.8 |
|  | Helixer | 0.849 | 0.930 | 0.726 | 0.736 | 0.219 | 0.249 | 90.0 |
|  | GeneMark | 0.474 | 0.438 | 0.211 | 0.206 | 0.031 | 0.018 | 24.0 |
| Myxine glutinosa<br>GCF_964187855.1<br>ref. BUSCO C 66.3 | Vipsania | 0.716 | 0.255 | 0.612 | 0.606 | 0.203 | 0.080 | 47.6 |
|  | Vipsania PT | 0.793 | 0.173 | 0.665 | 0.465 | 0.184 | 0.040 | 47.3 |
|  | Tiberius | 0.650 | 0.501 | 0.525 | 0.531 | 0.115 | 0.045 | 19.6 |
|  | ANNEVO | 0.494 | 0.592 | 0.410 | 0.562 | 0.060 | 0.053 | 19.3 |
|  | Helixer | 0.483 | 0.411 | 0.374 | 0.382 | 0.031 | 0.019 | 10.8 |
|  | GeneMark | 0.240 | 0.065 | 0.006 | 0.001 | 0.025 | 0.003 | 1.6 |
| Pristiophorus japonicus<br>GCF_044704955.1<br>ref. BUSCO C 94.6 | Vipsania | 0.655 | 0.663 | 0.569 | 0.774 | 0.318 | 0.267 | 70.2 |
|  | Vipsania PT | 0.733 | 0.391 | 0.647 | 0.559 | 0.301 | 0.109 | 77.8 |
|  | Tiberius | 0.882 | 0.688 | 0.803 | 0.675 | 0.389 | 0.170 | 84.6 |
|  | ANNEVO | 0.876 | 0.856 | 0.785 | 0.658 | 0.260 | 0.176 | 84.4 |
|  | Helixer | 0.755 | 0.659 | 0.667 | 0.470 | 0.094 | 0.064 | 63.5 |
|  | GeneMark | 0.005 | 0.005 | 0.000 | 0.000 | 0.000 | 0.000 | 0.0 |
| Tiliqua scincoides<br>GCF_035046505.1<br>ref. BUSCO C 95.5 | Vipsania | 0.908 | 0.814 | 0.817 | 0.830 | 0.446 | 0.400 | 88.4 |
|  | Vipsania PT | 0.828 | 0.764 | 0.734 | 0.797 | 0.344 | 0.334 | 81.4 |
|  | Tiberius | 0.941 | 0.847 | 0.866 | 0.826 | 0.528 | 0.413 | 92.2 |
|  | ANNEVO | 0.944 | 0.838 | 0.843 | 0.719 | 0.336 | 0.265 | 92.4 |
|  | Helixer | 0.903 | 0.744 | 0.806 | 0.574 | 0.165 | 0.134 | 86.0 |
|  | GeneMark | 0.219 | 0.157 | 0.038 | 0.022 | 0.007 | 0.003 | 4.6 |
| Xenopus tropicalis<br>GCF_000004195.4<br>ref. BUSCO C 97.3 | Vipsania | 0.889 | 0.760 | 0.807 | 0.827 | 0.476 | 0.423 | 89.9 |
|  | Vipsania PT | 0.865 | 0.669 | 0.771 | 0.777 | 0.399 | 0.312 | 86.3 |
|  | Tiberius | 0.915 | 0.873 | 0.858 | 0.846 | 0.537 | 0.463 | 93.1 |
|  | ANNEVO | 0.921 | 0.874 | 0.837 | 0.777 | 0.390 | 0.333 | 93.4 |
|  | Helixer | 0.880 | 0.814 | 0.804 | 0.614 | 0.163 | 0.147 | 86.0 |
|  | GeneMark | 0.691 | 0.310 | 0.430 | 0.232 | 0.059 | 0.019 | 39.3 |

Table 22: Per-species sensitivity / precision for test species not represented by one of the other 17 clade-specific **Vipsania** models. Rows list each tool's S and P per metric. Accession IDs appear below each species name.

| Species | Tool | Base |  | Exon |  | Locus |  |
| --- | --- | --- | --- | --- | --- | --- | --- |
|  |  | S | P | S | P | S | P |
| Branchiostoma lanceolatum<br>GCF_035083965.1 | Vipsania | 0.956 | 0.816 | 0.874 | 0.824 | 0.527 | 0.475 |
|  | Vipsania PT | 0.946 | 0.820 | 0.866 | 0.824 | 0.510 | 0.471 |
| Capsaspora owczarzaki<br>GCF_000151315.2 | Vipsania | 0.971 | 0.953 | 0.891 | 0.875 | 0.705 | 0.705 |
|  | Vipsania PT | 0.955 | 0.956 | 0.869 | 0.870 | 0.659 | 0.672 |
| Convolutriloba macropyga<br>GCF_964194025.1 | Vipsania | 0.835 | 0.670 | 0.718 | 0.692 | 0.413 | 0.328 |
|  | Vipsania PT | 0.817 | 0.678 | 0.711 | 0.686 | 0.387 | 0.317 |
| Gordionus montsenyensis<br>GCF_954871325.1 | Vipsania | 0.850 | 0.760 | 0.775 | 0.730 | 0.342 | 0.326 |
|  | Vipsania PT | 0.855 | 0.722 | 0.781 | 0.667 | 0.330 | 0.282 |
| Mnemiopsis leidyi<br>GCA_048537945.1 | Vipsania | 0.720 | 0.868 | 0.793 | 0.854 | 0.403 | 0.442 |
|  | Vipsania PT | 0.713 | 0.873 | 0.784 | 0.862 | 0.386 | 0.450 |
| Saccoglossus kowalevskii<br>GCF_000003605.3 | Vipsania | 0.793 | 0.791 | 0.724 | 0.752 | 0.315 | 0.355 |
|  | Vipsania PT | 0.766 | 0.796 | 0.712 | 0.742 | 0.286 | 0.338 |
| Trichomonas vaginalis<br>GCF_026262505.1 | Vipsania | 0.870 | 0.704 | 0.415 | 0.306 | 0.415 | 0.372 |
|  | Vipsania PT | 0.896 | 0.678 | 0.426 | 0.272 | 0.426 | 0.337 |

#### 3 Sampled Clades

For each clade, the NCBI taxonomy of the sampled species was converted into a clock-like (ultra-metric) tree. A genome assembly was considered if its assembly level was *chromosome* or *complete*, it was not flagged as atypical by NCBI, it spanned at least 2 Mb and it consisted of at most 20,000 scaffolds. For AMOEBOZOA and RHODOPHYTA too few assemblies reached chromosome level, so for these two clades the requirement was relaxed to *scaffold* or better. From all species with such a genome assembly a subset of given size was constructed with a greedy algorithm that guarantees to maximize phylogenetic diversity. Figures 3–20 show the resulting trees, one per clade. Branch lengths are relative, the root of each tree being at 0 and all tips at 1; they are derived from taxonomic ranks and are a proxy for divergence time rather than dated divergence estimates.

Tips set in italics are the sampled species. Shaded rows mark clades that were deliberately held out of the sample, so that the case of a genome unseen at training time could be evaluated; a single representative tip, labeled with the clade name, stands in for the whole held-out clade. Trees with many tips are laid out in several columns on one page and are read left to right.

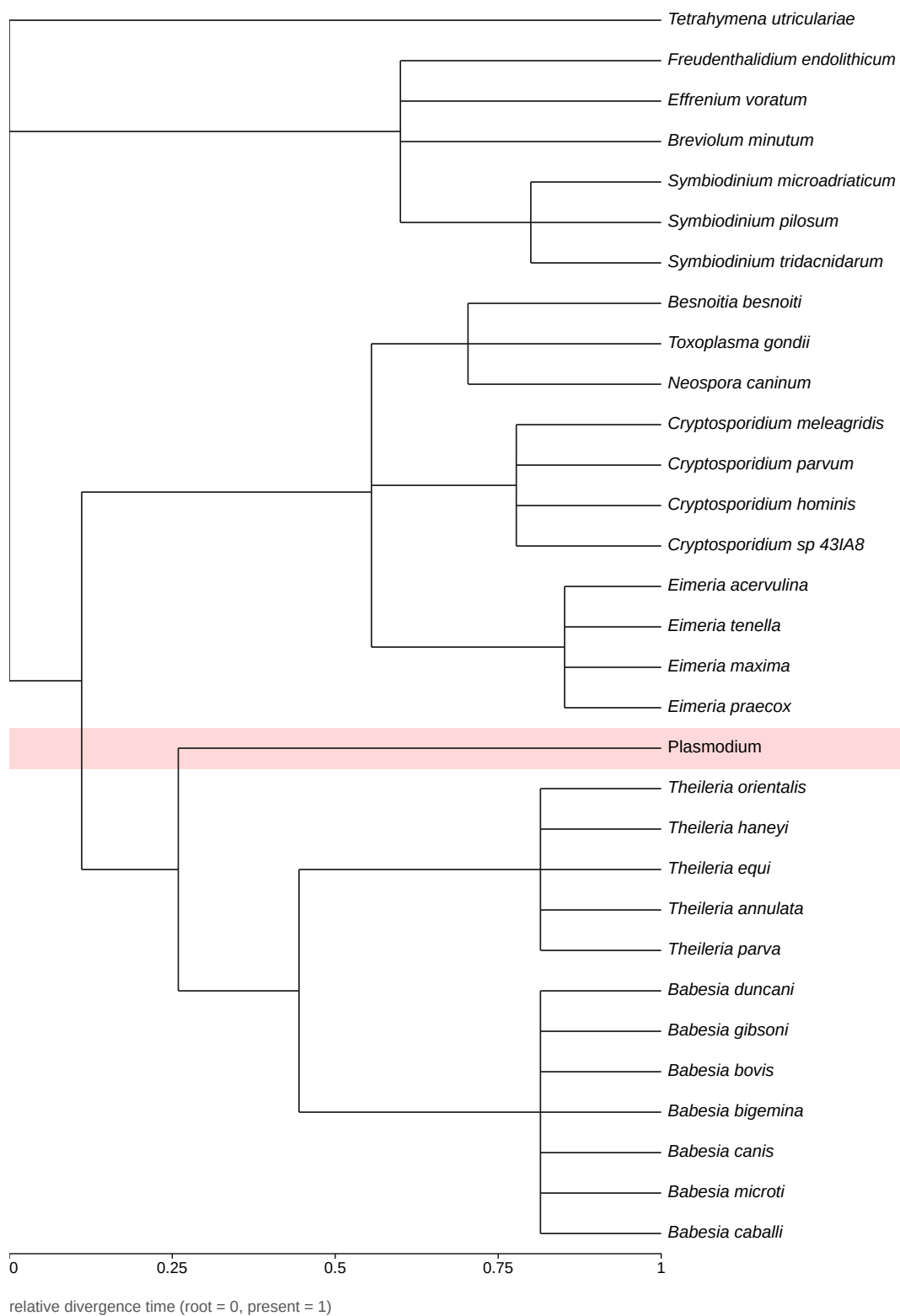

Figure 3: Clock-like tree of the 30 species sampled from ALVEOLATA (NCBI taxid 33630), with one held-out clade, shaded.

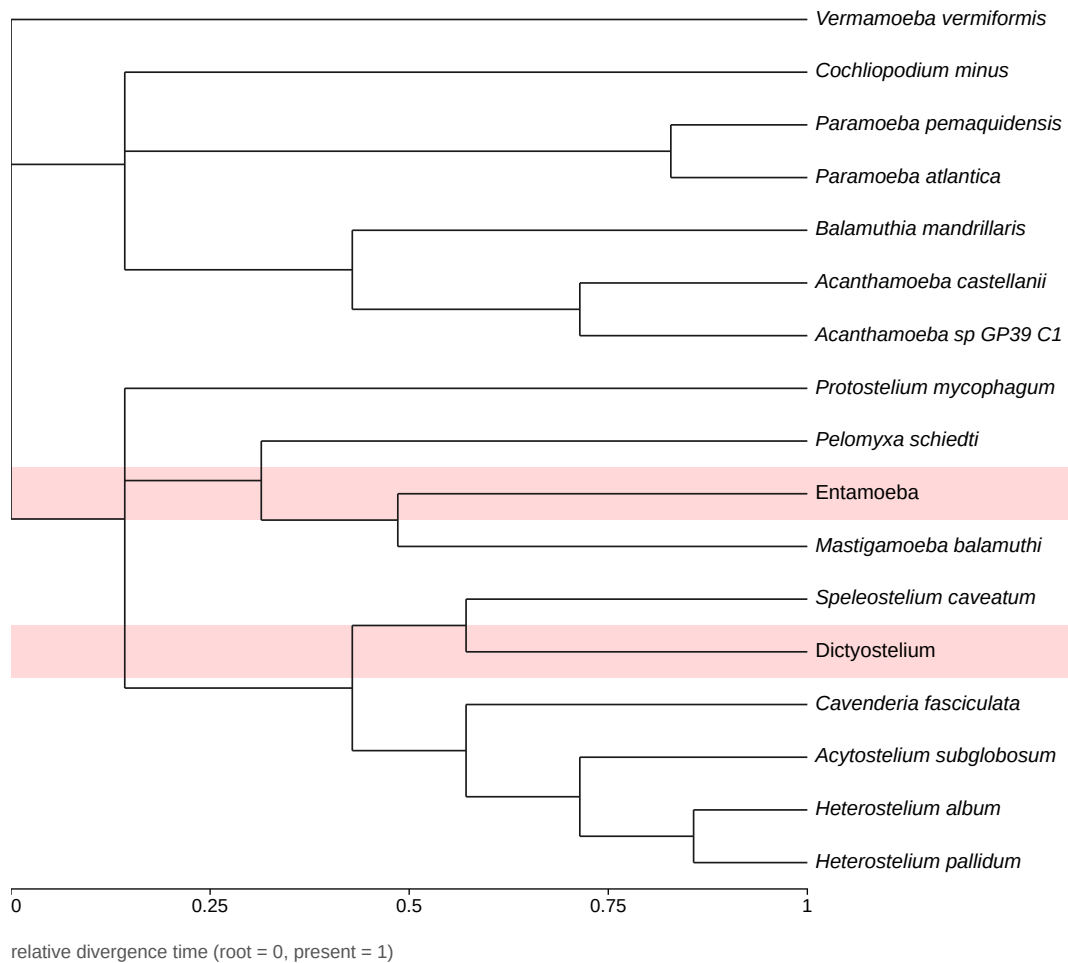

Figure 4: Clock-like tree of the 15 species sampled from **AMOEBOZOA** (NCBI taxid 554915), with 2 held-out clades, shaded.

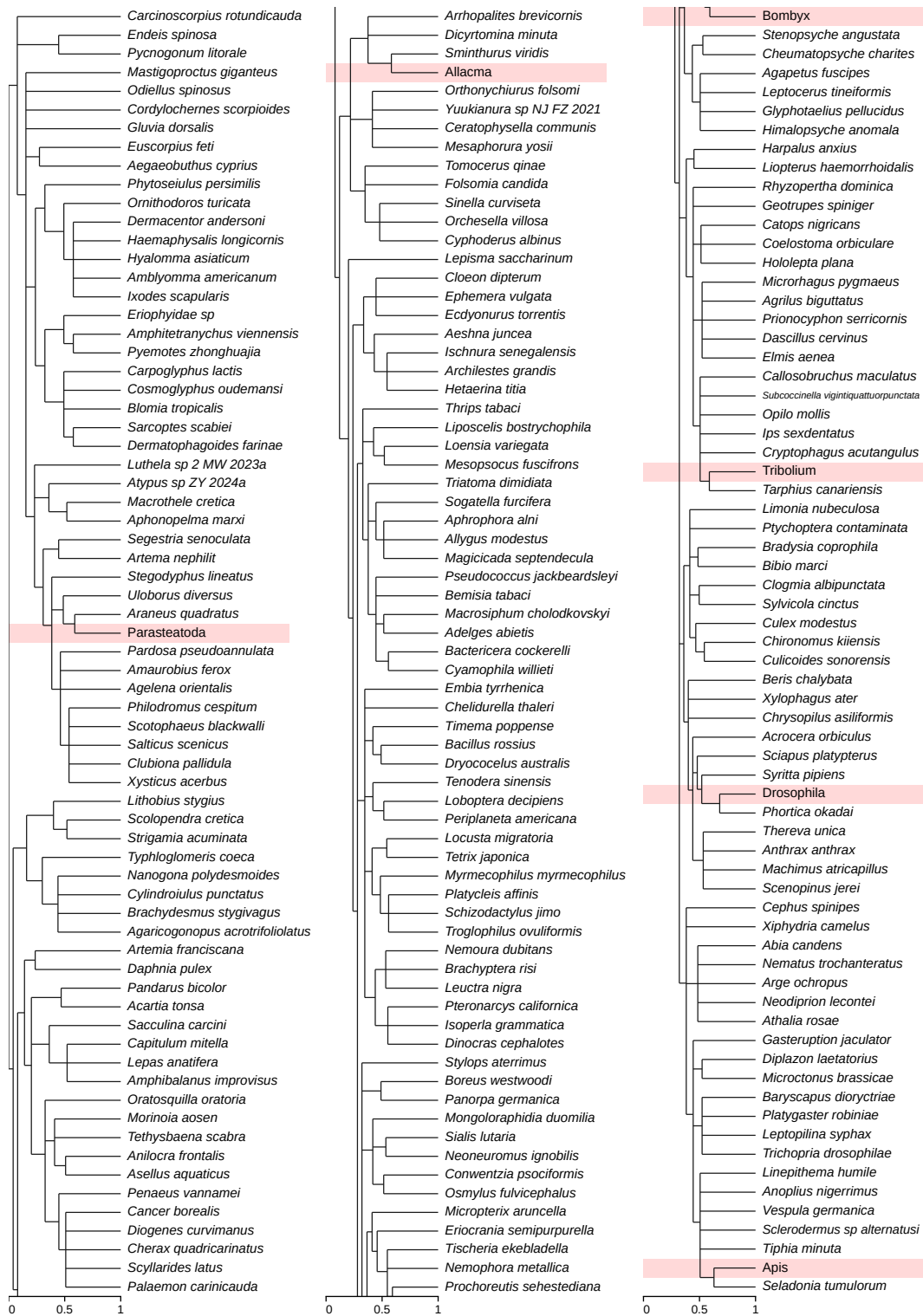

Figure 5: Clock-like tree of the 200 species sampled from **ARTHROPODA** (NCBI taxid 6656), with 6 held-out clades, shaded. The tree is continued across 3 columns, read left to right.

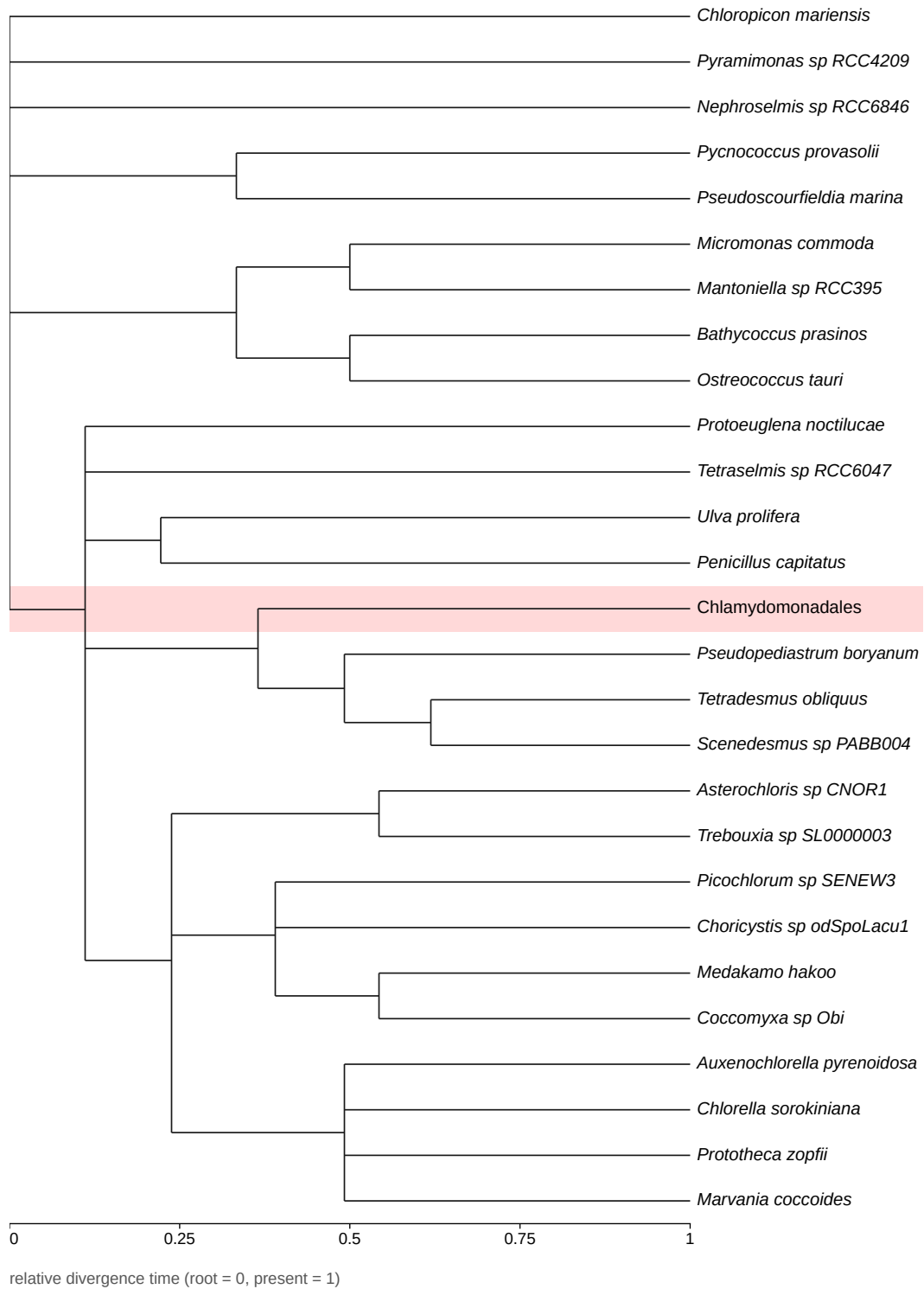

Figure 6: Clock-like tree of the 26 species sampled from **CHLOROPHYTA** (NCBI taxid 3041), with one held-out clade, shaded.

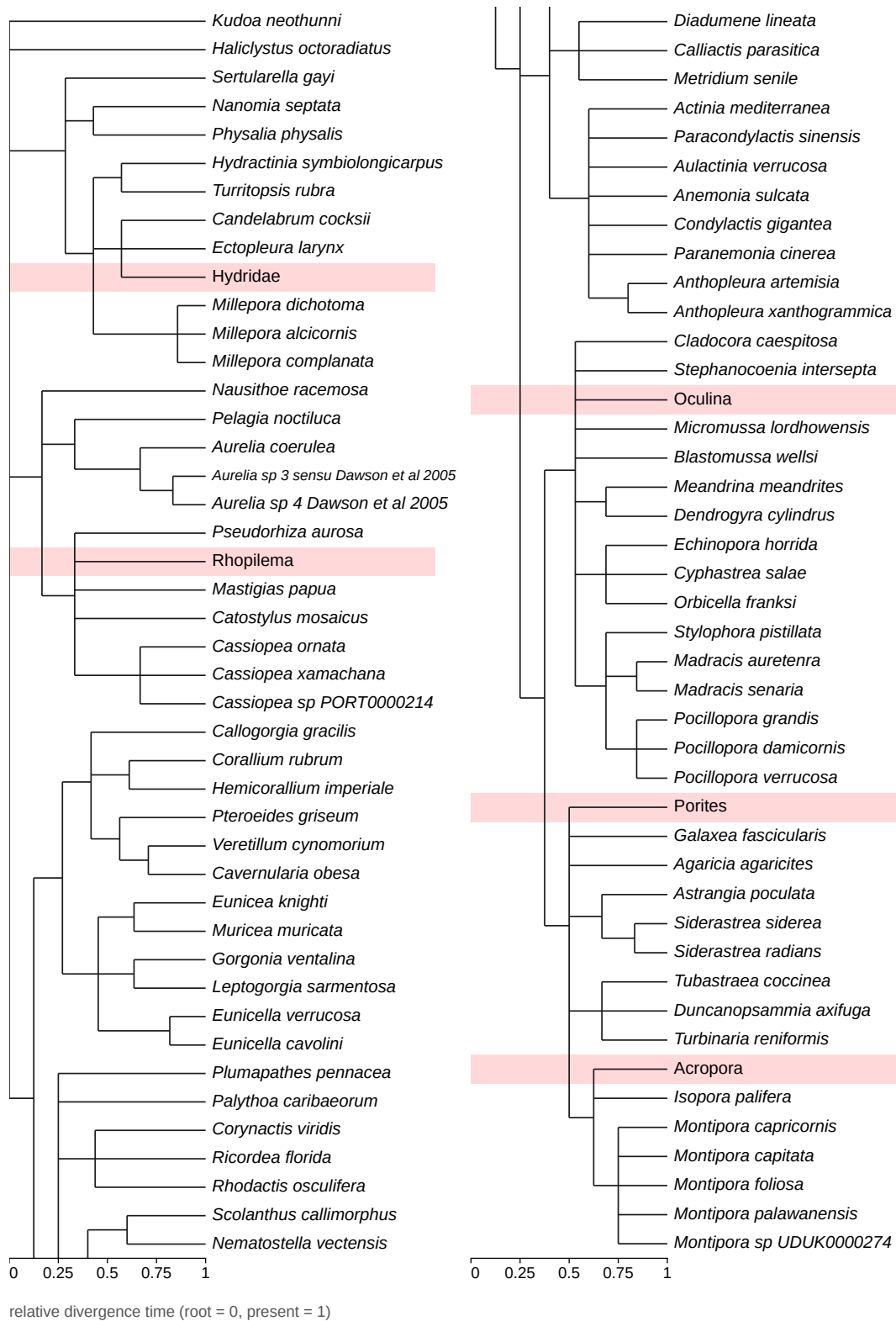

Figure 7: Clock-like tree of the 82 species sampled from **Cnidaria** (NCBI taxid 6073), with 5 held-out clades, shaded. The tree is continued across 2 columns, read left to right.

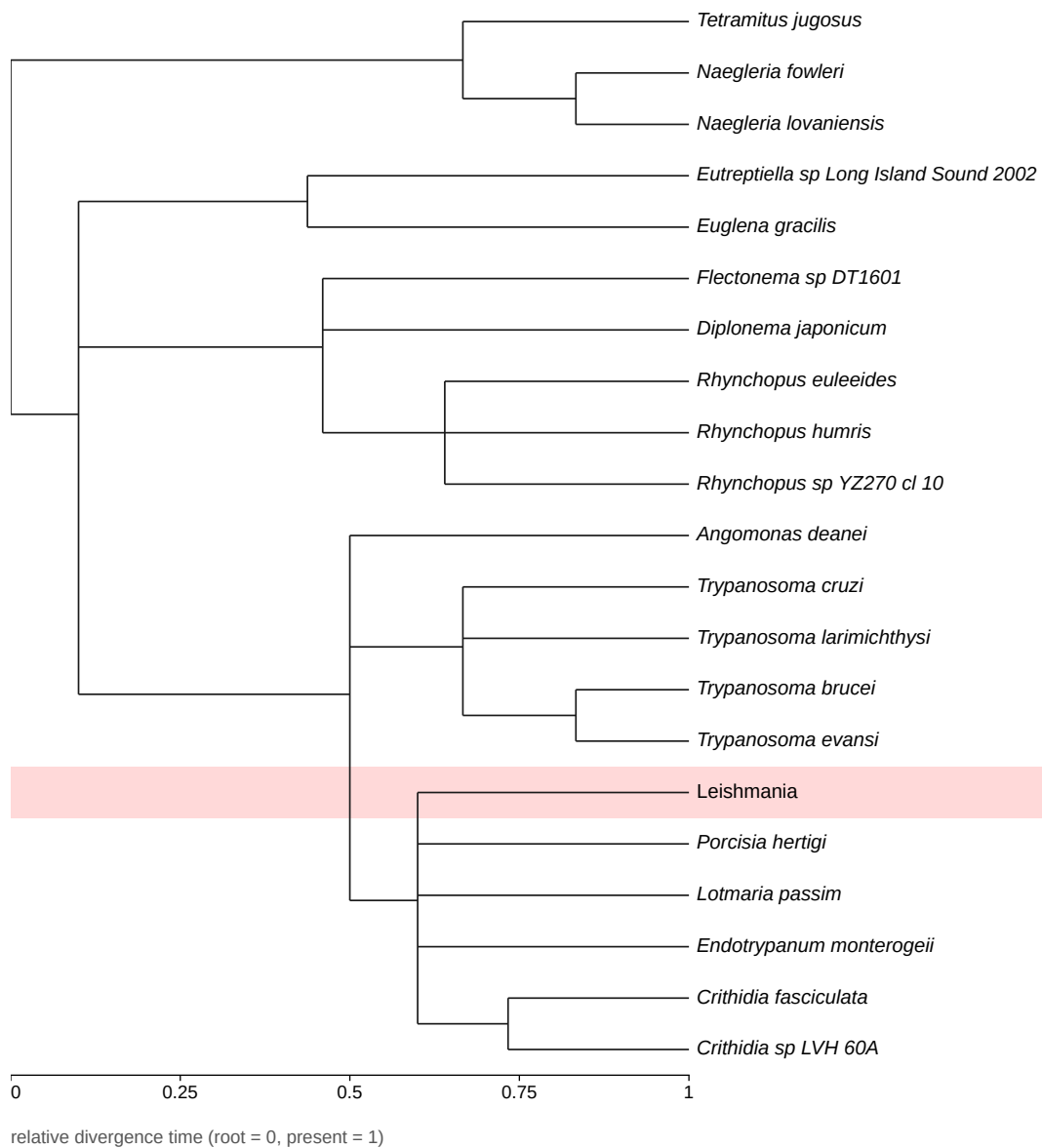

Figure 8: Clock-like tree of the 20 species sampled from **DISCOBA** (NCBI taxid 2611352), with one held-out clade, shaded.

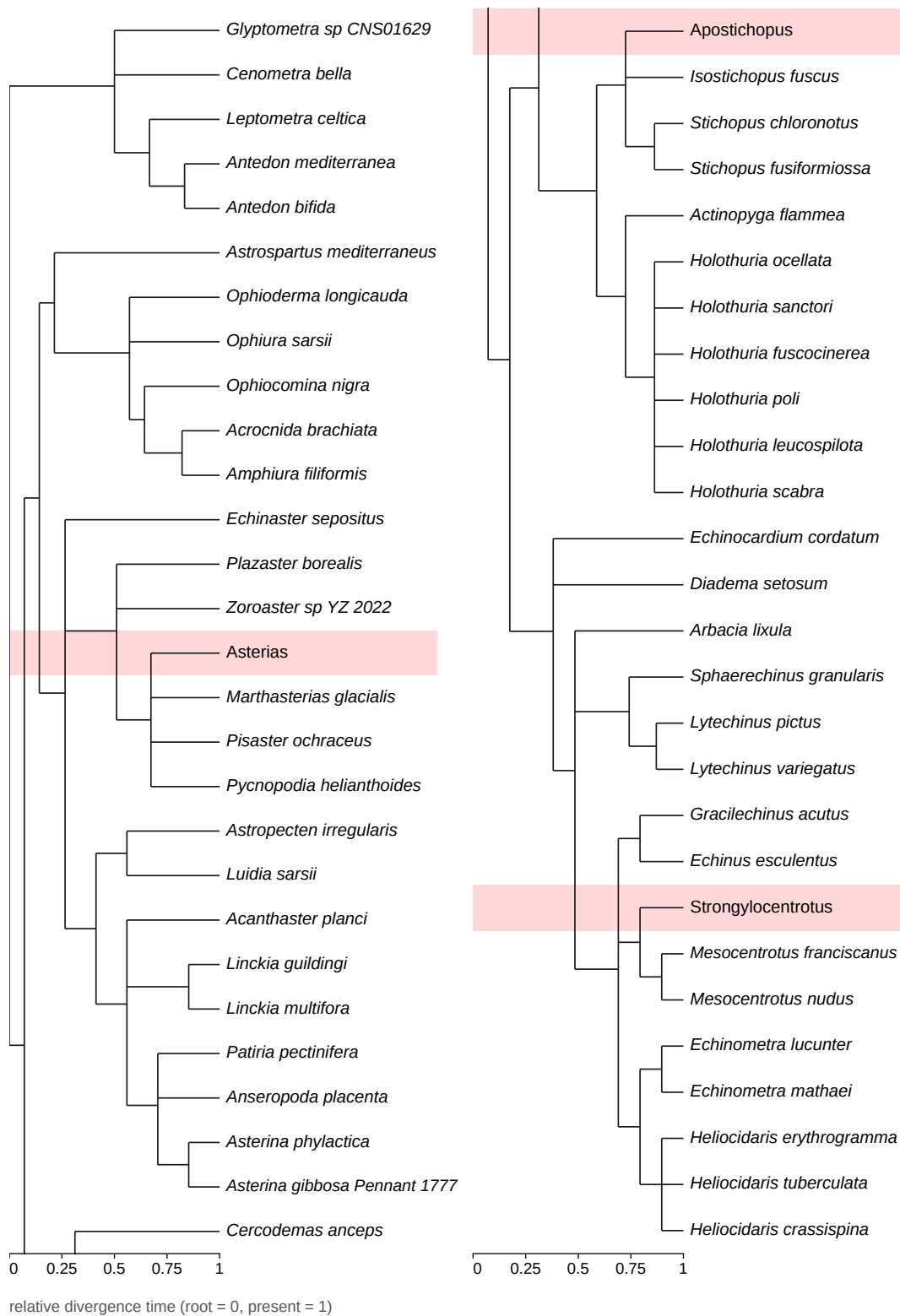

Figure 9: Clock-like tree of the 52 species sampled from ECHINODERMATA (NCBI taxid 7586), with 3 held-out clades, shaded. The tree is continued across 2 columns, read left to right.

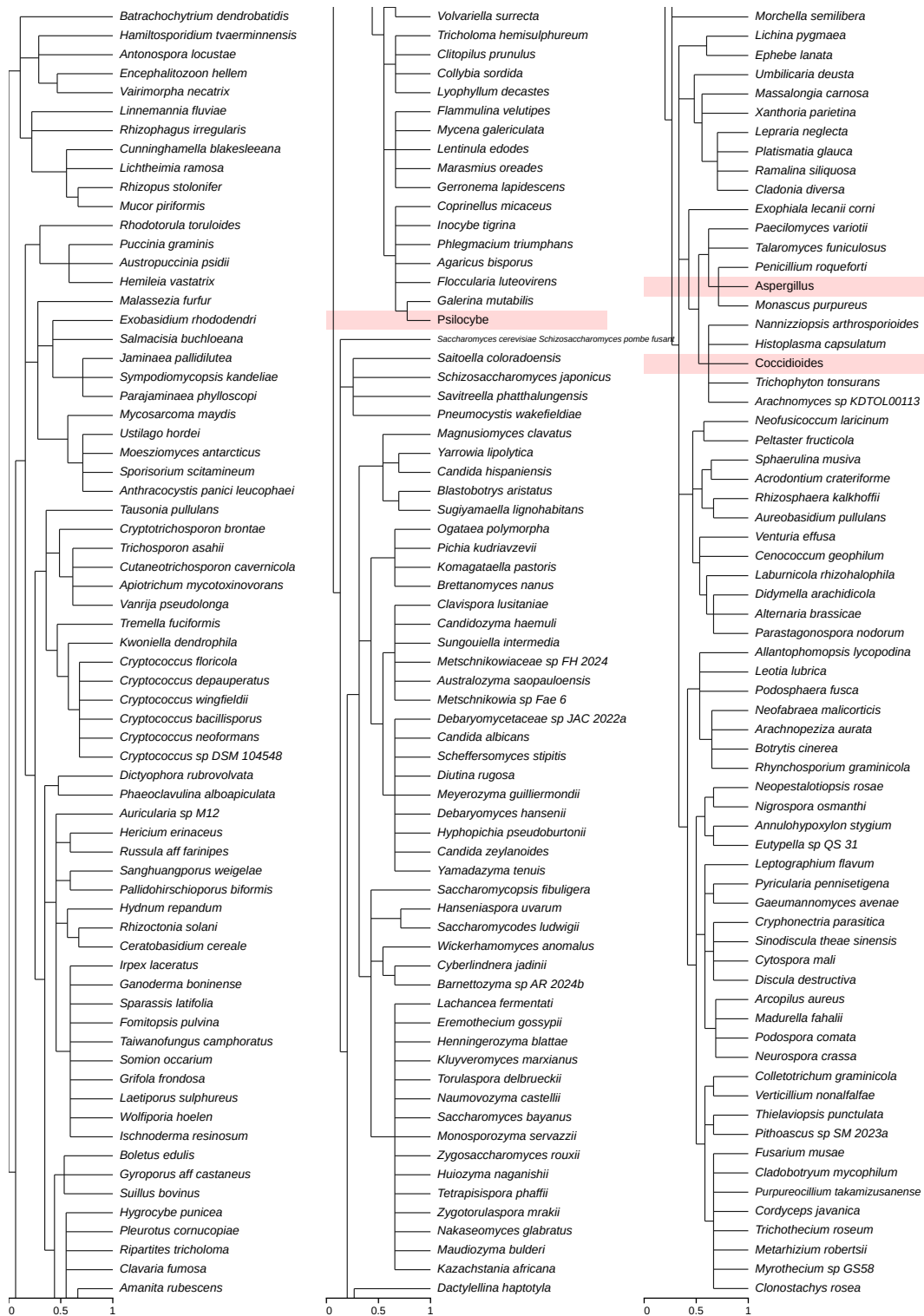

Figure 10: Clock-like tree of the 200 species sampled from FUNGI (NCBI taxid 4751), with 3 held-out clades, shaded. The tree is continued across 3 columns, read left to right.

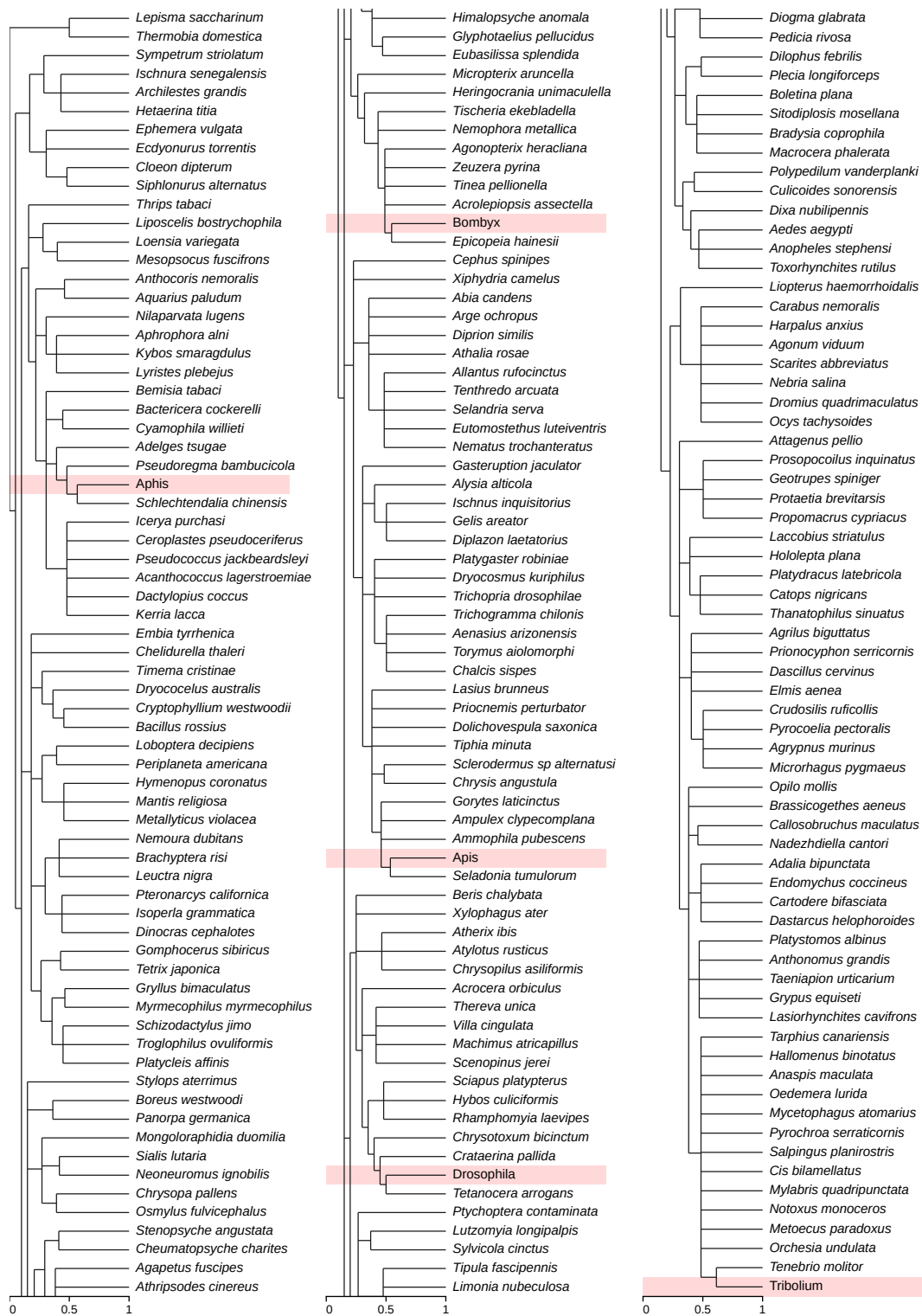

relative divergence time (root = 0, present = 1)

Figure 11: Clock-like tree of the 200 species sampled from INSECTA (NCBI taxid 50557), with 5 held-out clades, shaded. The tree is continued across 3 columns, read left to right.

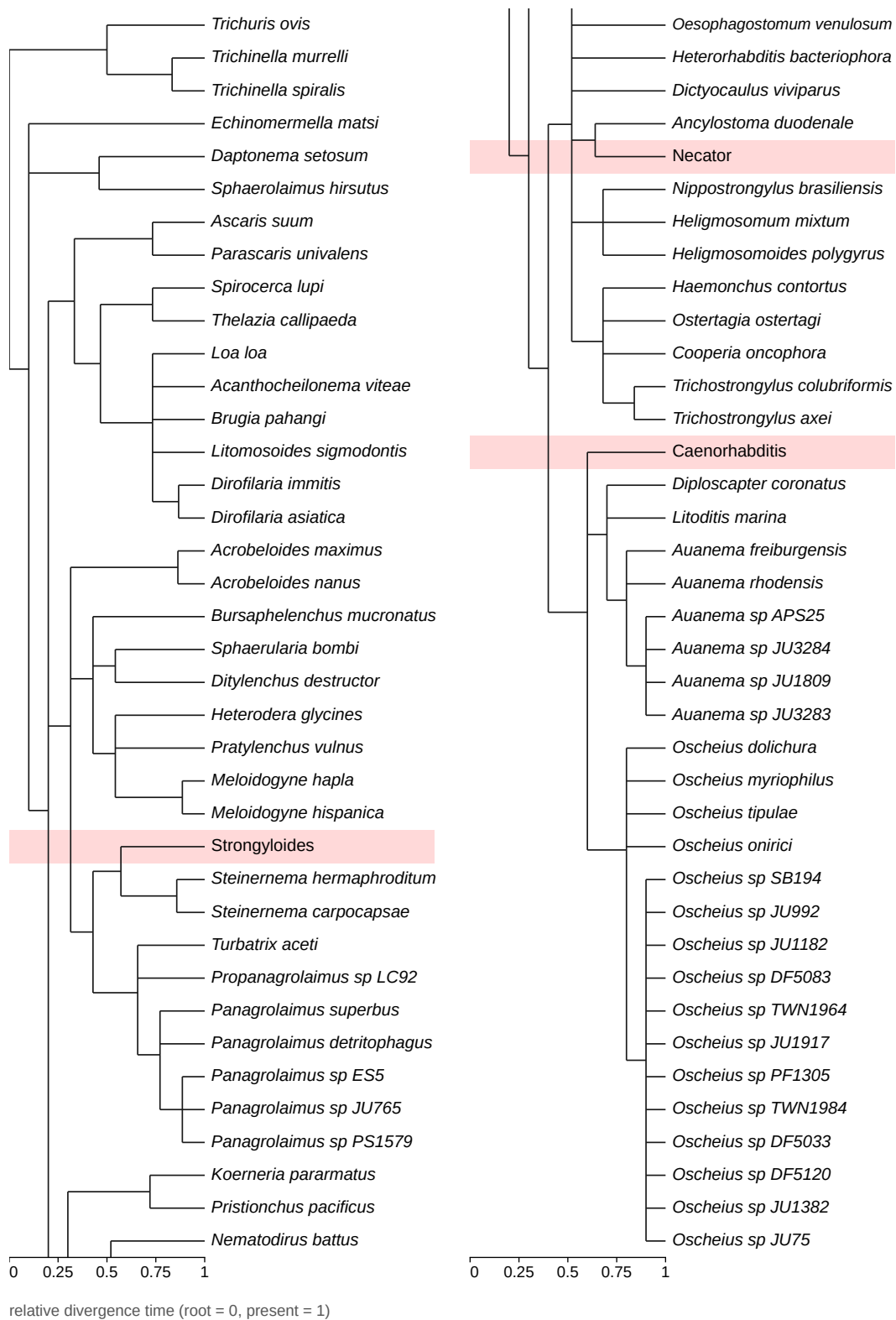

Figure 12: Clock-like tree of the 73 species sampled from **NEMATODA** (NCBI taxid 6231), with 3 held-out clades, shaded. The tree is continued across 2 columns, read left to right.

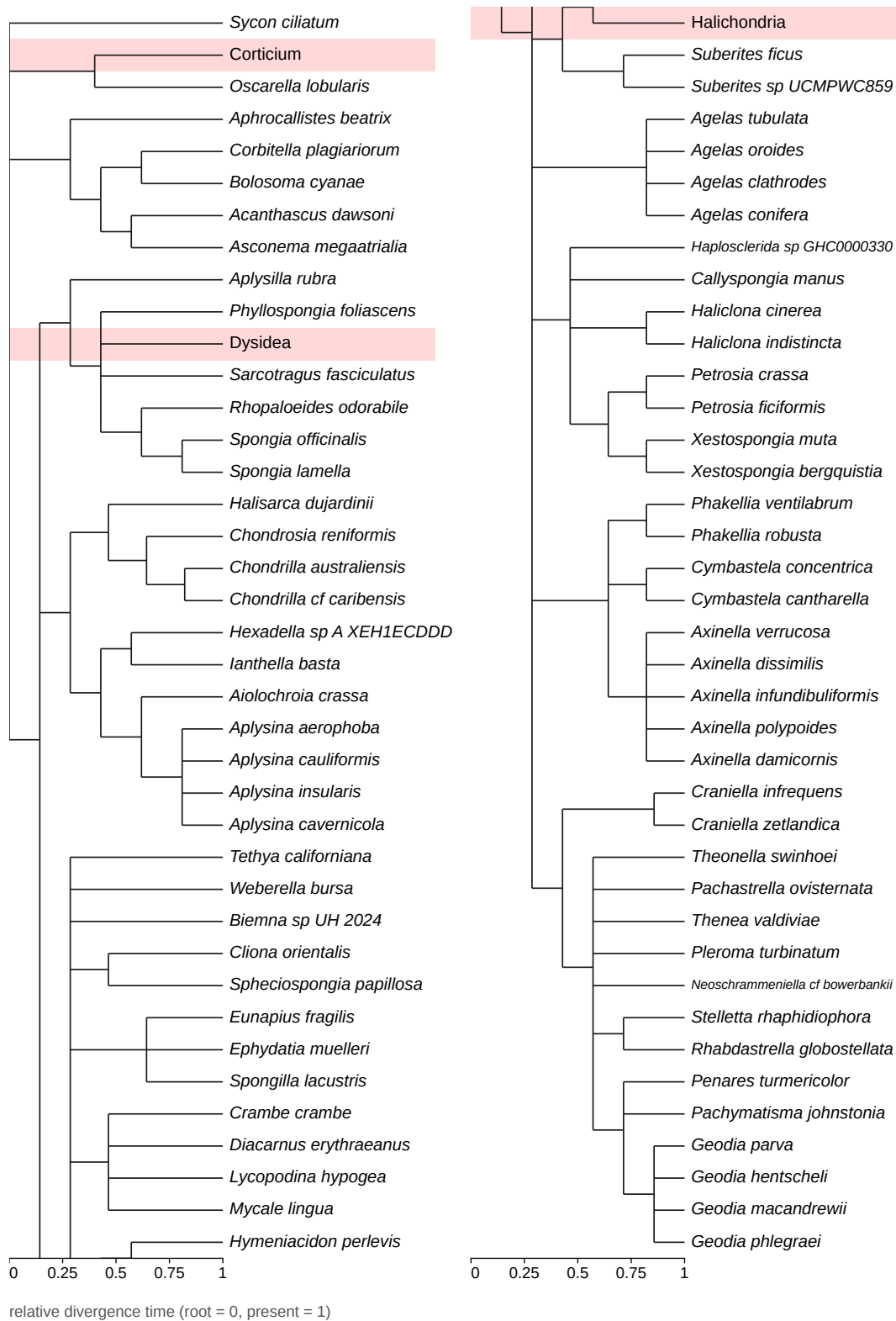

Figure 13: Clock-like tree of the 75 species sampled from **PORIFERA** (NCBI taxid 6040), with 3 held-out clades, shaded. The tree is continued across 2 columns, read left to right.

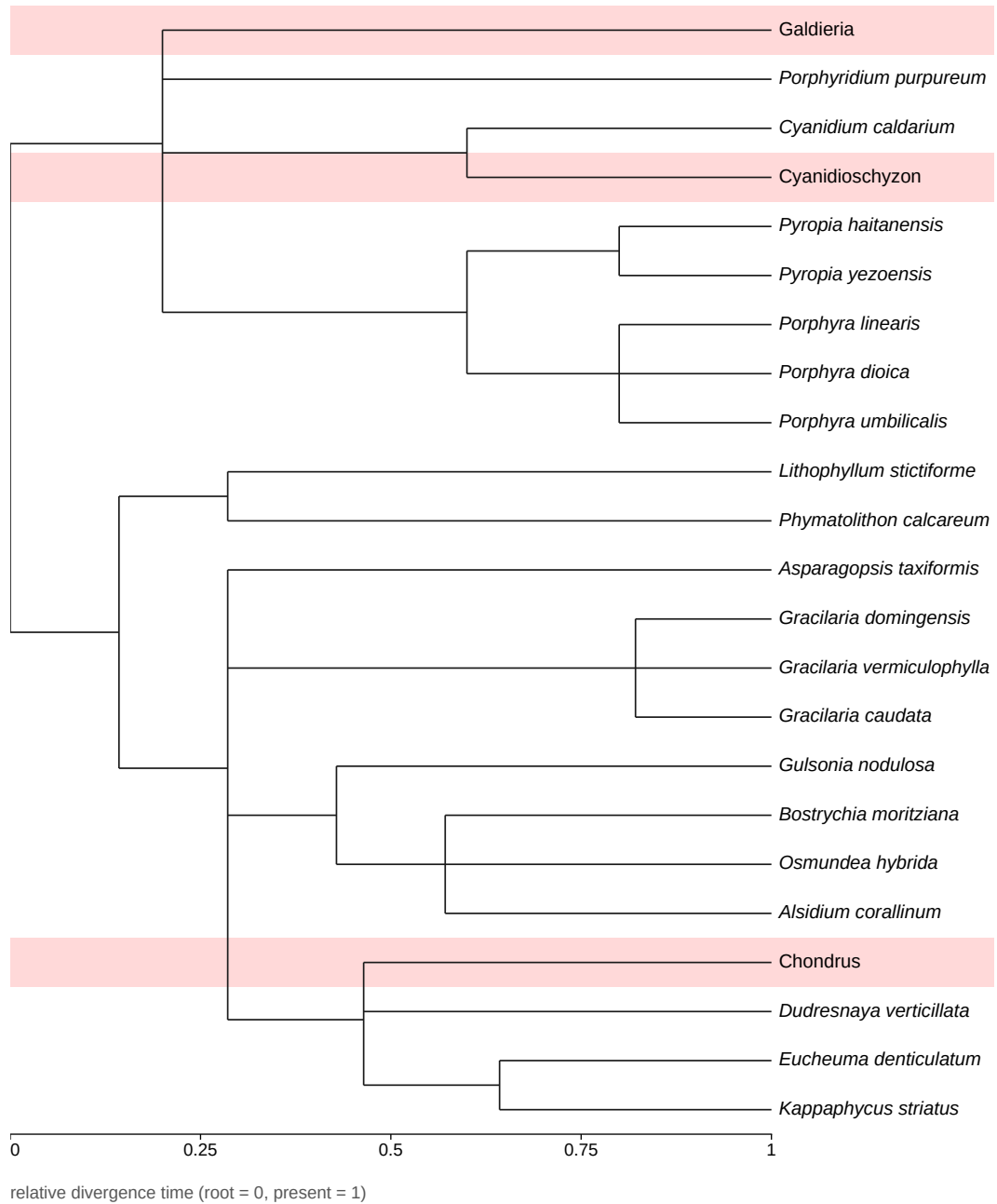

Figure 14: Clock-like tree of the 20 species sampled from **RHODOPHYTA** (NCBI taxid 2763), with 3 held-out clades, shaded.

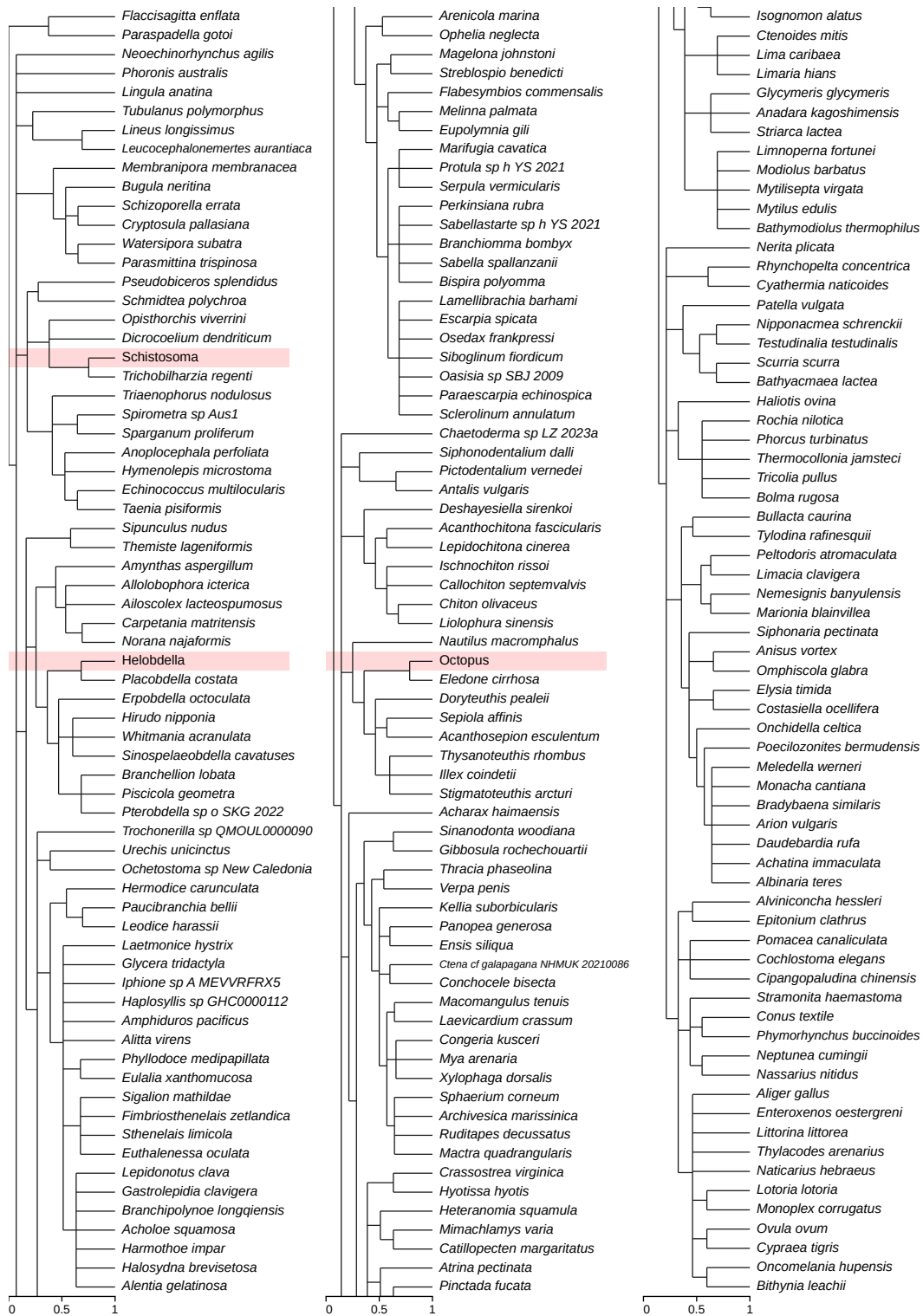

Figure 15: Clock-like tree of the 200 species sampled from **SPIRALIA** (NCBI taxid 2697495), with 3 held-out clades, shaded. The tree is continued across 3 columns, read left to right.

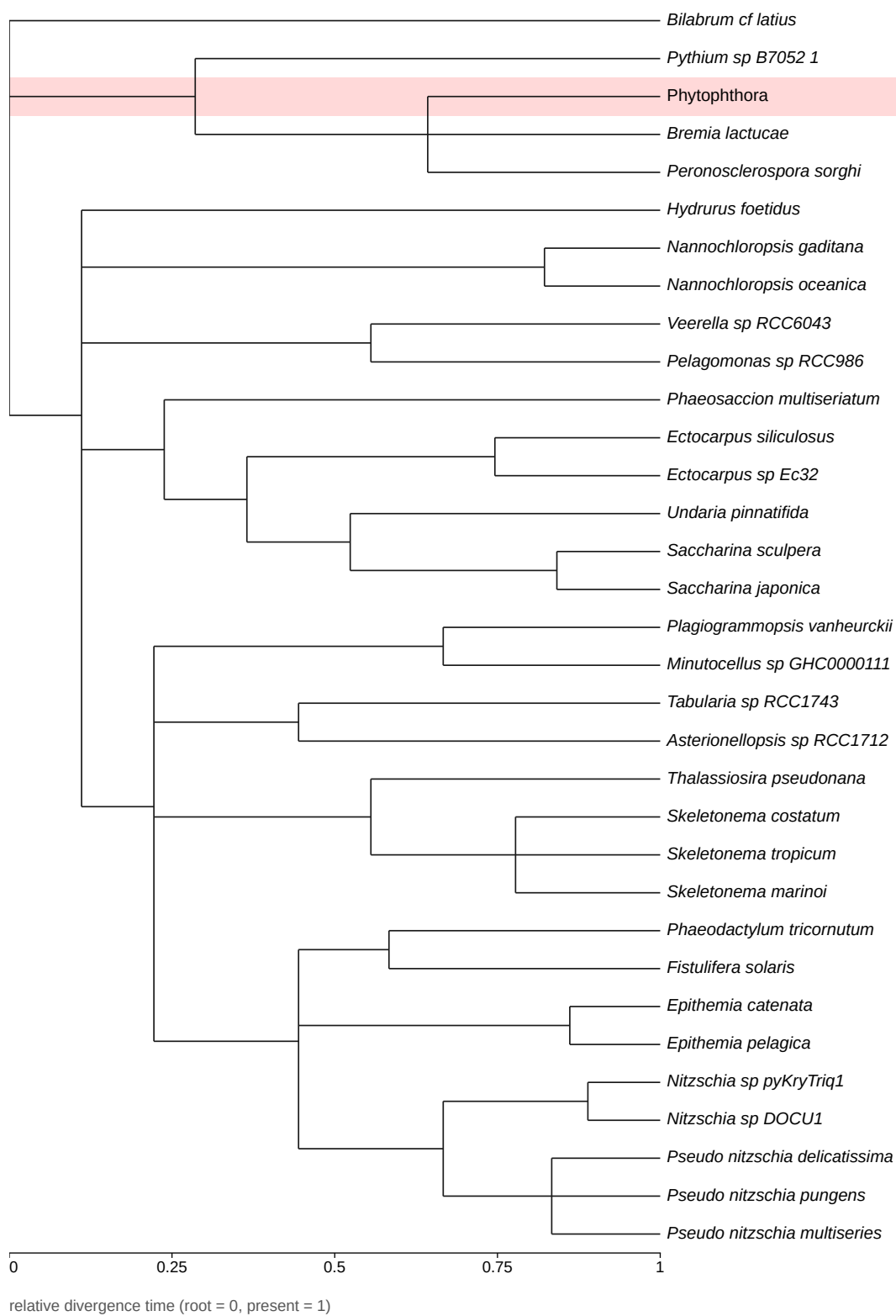

Figure 16: Clock-like tree of the 32 species sampled from **STRAMENOPILES** (NCBI taxid 33634), with one held-out clade, shaded.

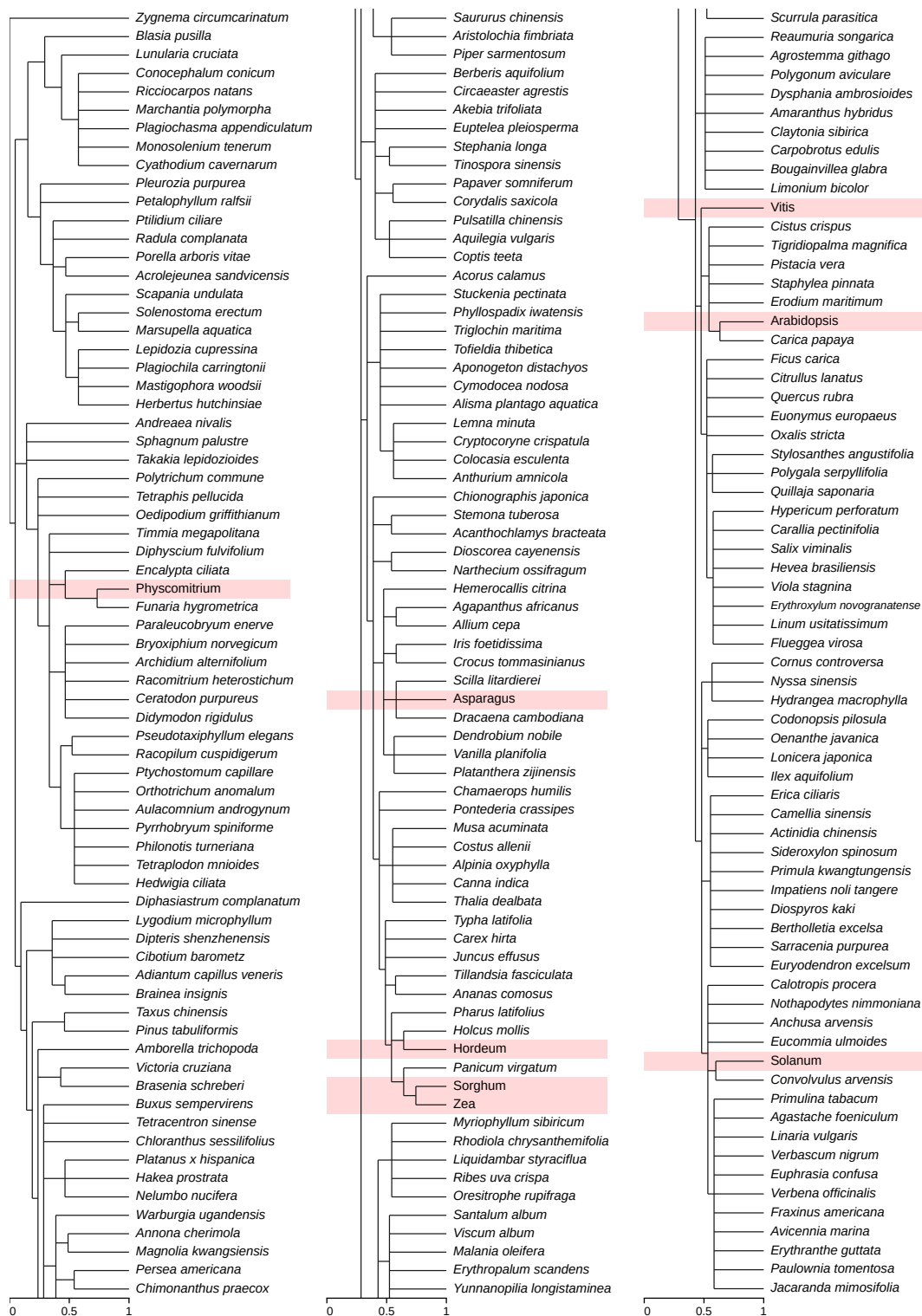

relative divergence time (root = 0, present = 1)

Figure 17: Clock-like tree of the 200 species sampled from **STREPTOPHYTA** (NCBI taxid 35493), with 8 held-out clades, shaded. The tree is continued across 3 columns, read left to right.

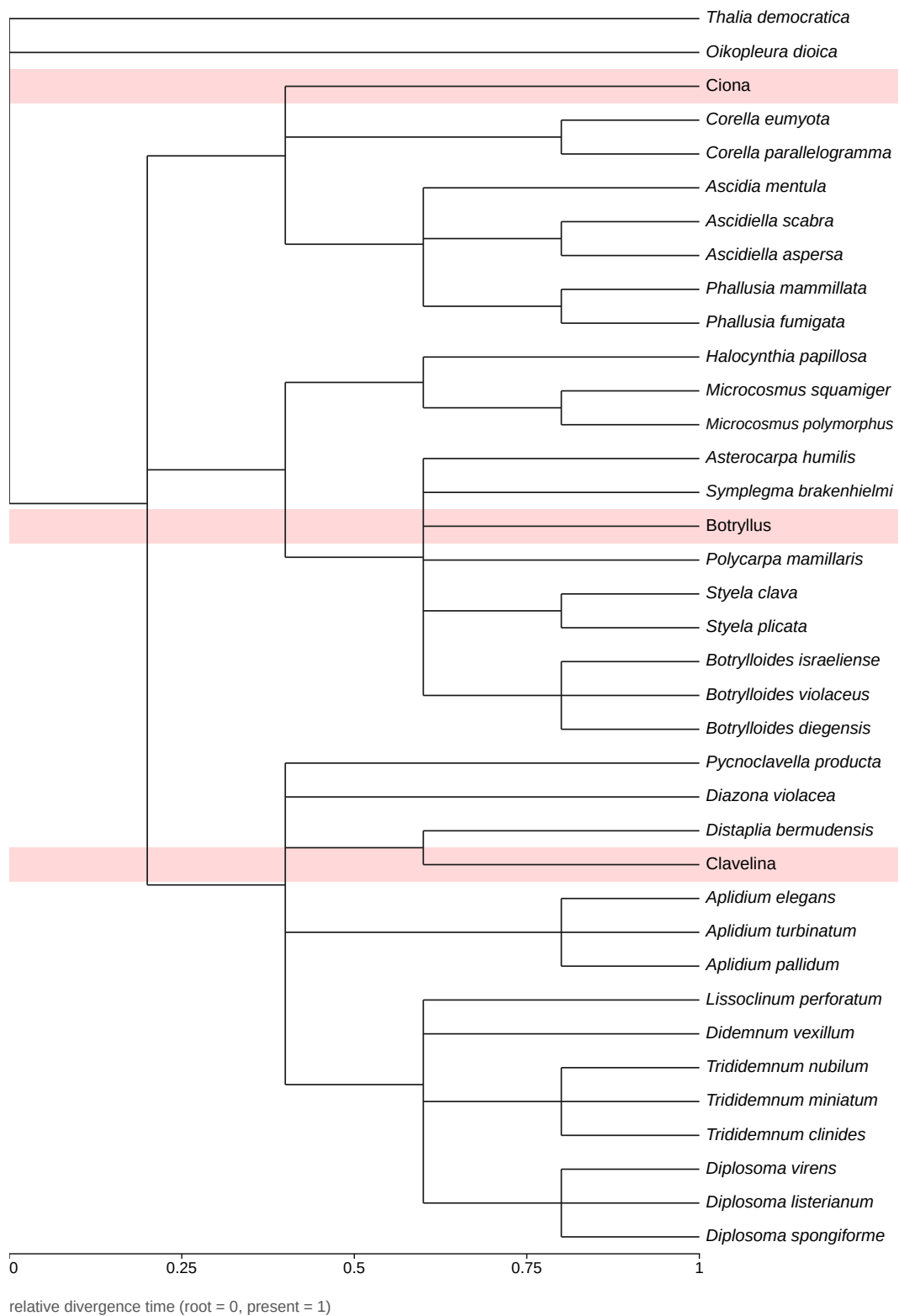

Figure 18: Clock-like tree of the 34 species sampled from TUNICATA (NCBI taxid 7712), with 3 held-out clades, shaded.

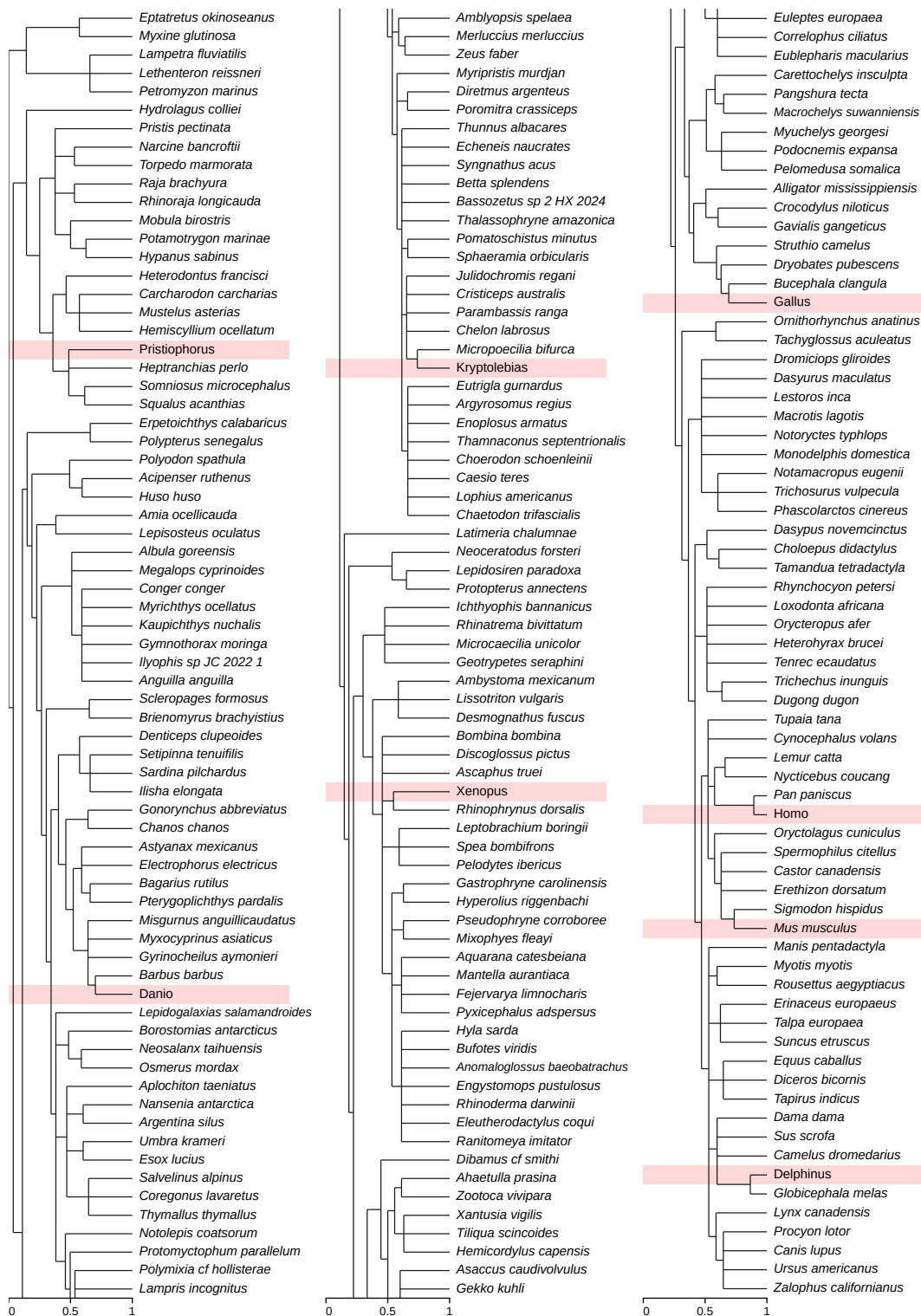

relative divergence time (root = 0, present = 1)

Figure 19: Clock-like tree of the 200 species sampled from VERTEBRATA (NCBI taxid 7742), with 8 held-out clades, shaded. The tree is continued across 3 columns, read left to right.

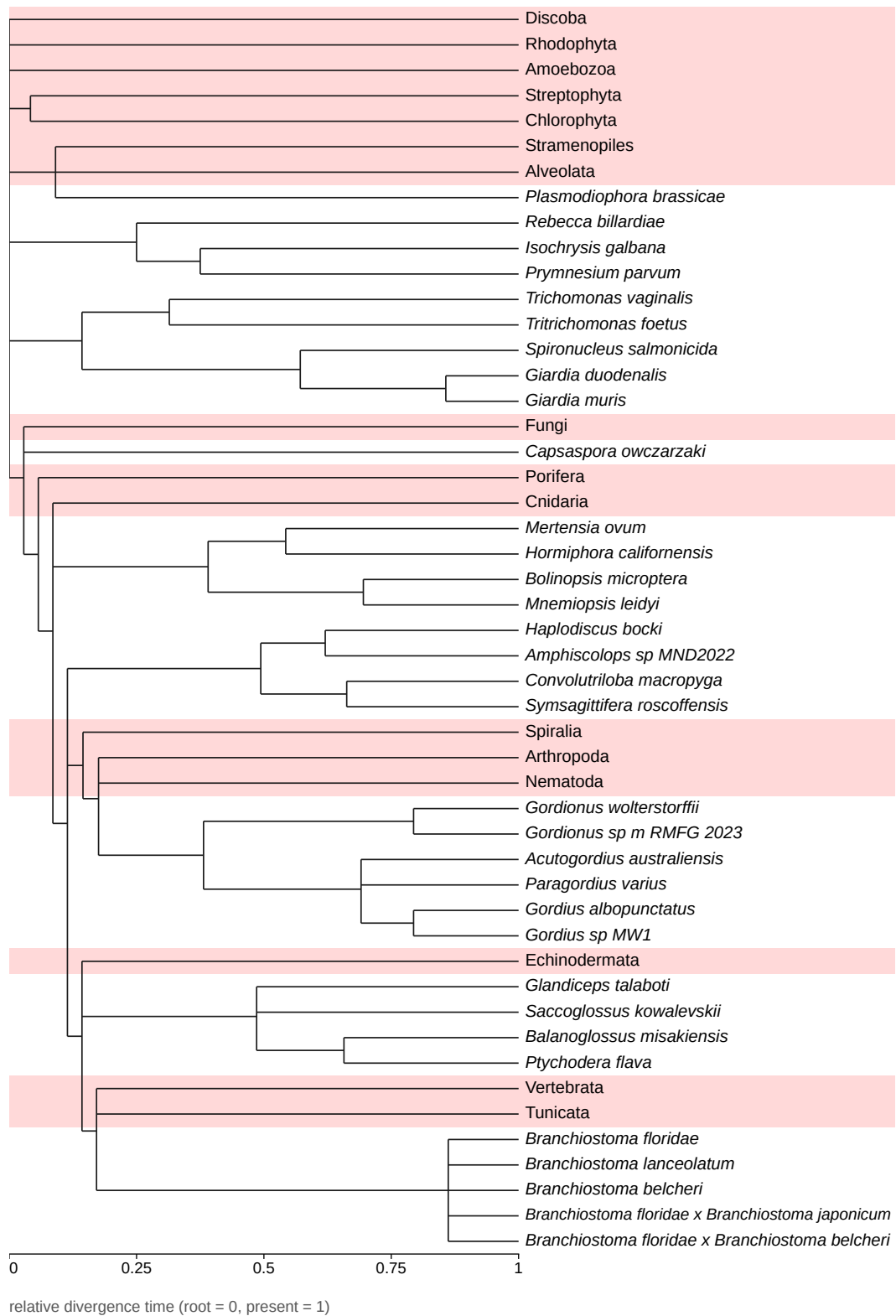

Figure 20: Clock-like tree of the 33 species sampled from **OTHER EUKARYOTES** (NCBI taxid 2759), with 16 held-out clades, shaded.
